# Patterns of recombination in snakes reveal a tug of war between PRDM9 and promoter-like features

**DOI:** 10.1101/2023.07.11.548536

**Authors:** Carla Hoge, Marc de Manuel, Mohamed Mahgoub, Naima Okami, Zachary Fuller, Shreya Banerjee, Zachary Baker, Morgan McNulty, Peter Andolfatto, Todd S. Macfarlan, Molly Schumer, Athanasia C. Tzika, Molly Przeworski

**Author notes:** These authors contributed equally to this work (Carla Hoge, Marc de Manuel). Division of Oncology, Department of Medicine, Washington University School of Medicine, St. Louis, MO. Center for Population Biology, University of California, Davis, CA. 23andMe (Zachary Fuller). Dept. of Genetics, Cambridge University (Zachary Baker).

## Abstract

In vertebrates, there are two known mechanisms by which meiotic recombination is directed to the genome: in humans, mice, and other mammals, recombination occurs almost exclusively where the protein PRDM9 binds, while in species lacking an intact *PRDM9*, such as birds and canids, recombination rates are elevated near promoter-like features. To test if PRDM9 also directs recombination in non-mammalian vertebrates, we focused on an exemplar species, the corn snake (*Pantherophis guttatus*). Unlike birds, this species possesses a single, intact *PRDM9* ortholog. By inferring historical recombination rates along the genome from patterns of linkage disequilibrium and identifying crossovers in pedigrees, we found that PRDM9 specifies the location of recombination events outside of mammals. However, we also detected an independent effect of promoter-like features on recombination, which is more pronounced on macro- than microchromosomes. Thus, our findings reveal that the uses of PRDM9 and promoter-like features are not mutually-exclusive, and instead reflect a tug of war, which varies in strength along the genome and is more lopsided in some species than others.

**One sentence summary:** While the localization of meiotic recombination in vertebrates was previously thought to occur using one of two distinct mechanisms, our analysis of recombination in corn snakes reveals that they and likely other vertebrates use both of these mechanisms.

## Main text

Meiotic recombination begins with the intentional creation of double strand breaks (DSB) in the genome, the repair of which results in crossovers and non-crossovers (*1*). In vertebrates as well as other taxa, such DSBs are concentrated in short genomic segments (typically 1-2 kilobases), known as “recombination hotspots.” The mechanisms that specify the genomic location of hotspots differ among species. In mice, humans, and other mammals, the gene *PRDM9* (PR/SET Domain 9) plays a key role (*2*–*6*). In mouse and rat, outside the pseudoautosomal region (*6, 7*), almost all hotspot locations are determined by PRDM9 binding: for instance, congenic mouse strains with different *Prdm9* alleles share only ∼1% of their DSB hotspots (*7*) and introducing a human *PRDM9* allele into B6 mice shifts the location of over 97% of DSB hotspots (*8*).

PRDM9 is a zinc-finger protein that binds DNA with sequence specificity and tri-methylates histone H3 lysine 4 (H3K4me3) and H3 lysine 36 (H3K36me3) (*9*). In mice, the SET domain that carries out these modifications is necessary for PRDM9-dependent DSB formation (*9*). PRDM9 also has two N-terminal domains that are essential for its role in recombination, but whose specific functions remain unclear (*10*–*12*). This four domain gene structure originated before the ancestor of vertebrates, but has been lost, in whole or in part, at least 13 times independently (*13*).

In species that carry a pseudogene for *PRDM9* (e.g., canids; (*14*)), are missing N-terminal domains (e.g., several lineages of ray-finned fish; (*12, 13*)) or lack *PRDM9* entirely (e.g., birds; (*15, 16*)), crossover rates are elevated near promoter-like features, notably CpG islands and transcription start sites (TSS) (*12, 16, 17*). These features are typically marked by H3K4me3–but not H3K36me3 (*18*)–through molecular processes that serve purposes other than initiating meiotic recombination (*19*). A possible explanation is that, in the absence of the marks made by PRDM9, the machinery responsible for making DSBs defaults to residual sites of H3K4me3 (*7*), either because the mark causes its recruitment–say via a reader protein–or simply because such regions are more accessible. Consistent with the notion that species lacking *PRDM9* rely on other H3K4me3 marks, when *Prdm9* is knocked-out in B6 mice or SHR rats, ∼92% and ∼99% of DSBs occur at sites of H3K4me3, respectively (*6, 7*). The knock-out experiments in mice and rats, alongside the report of a human mother homozygous for a non-functional *PRDM9* (*20*), further indicate that, in species with a complete *PRDM9*, there exists an alternative mechanism of recombination that can serve as a back-up, albeit a less efficient one (*7, 8, 21*).

In mammals in which PRDM9 is known to direct recombination, the zinc finger domain of the protein shows evidence of positive selection at residues in contact with DNA and the binding affinity is rapidly evolving (*3, 12*). Modeling work suggests that this rapid evolution is an expected consequence of the roles of PRDM9 in recombination (*22, 23*). Intriguingly then, similar evidence for positive selection is also seen in all the non-mammalian vertebrates with a complete PRDM9 ortholog that have been surveyed to date (and only in those species with a complete ortholog) (*12, 13*). These findings suggest that PRDM9 directs recombination outside of mammals (*12*). We tested this prediction by examining recombination patterns in an exemplar species with a single, complete PRDM9 ortholog, the corn snake *Pantherophis guttatus*. To this end, we inferred historical recombination rates from patterns of linkage disequilibrium (LD) across the genome and inferred crossover events from pedigrees.

### Genome sequencing

As a first step, we improved the corn snake reference genome to nearly chromosome-scale (scaffold N50: 63 Mb) by complementing an existing hybrid assembly (*24*) with PacBio long read sequencing (see Methods). We further collected Pacbio RNA-seq data and bisulfite sequencing data from two testis samples to better annotate transcription start sites and CpG islands, respectively (see Methods).

In order to infer historical recombination rates along the genome and identify crossovers in families, we resequenced whole genomes from a total of 37 individuals at medium to high coverage (13-48X, median 23X) using Illumina sequencing (Table S1). Our samples include 19 wild-caught individuals collected from across the southeastern United States (Figure 1A). To those, we added 18 individuals from a lab colony, of which 16 come from two nuclear families and five are not close relatives (Figure 1B). Although samples from these two sets form distinct clusters in the first principal component of a principal component analysis (Figure S1), Fst (*25*) between the sets is relatively low (mean across 10 kb windows = 0.075). Combining the samples, we called single nucleotide polymorphisms (SNPs) in non-repetitive genomic regions of autosomal scaffolds, using inheritance patterns in the families to exclude sites at which alleles violated rules of Mendelian segregation (0.31% of polymorphic sites) as well as any region harboring an unusually large number of such errors (see Methods). Among the 24 individuals who are not close relatives (henceforth “unrelated”), we identified >11 million SNPs, corresponding to a mean heterozygosity (π) of 0.0032 per site across 10 kb windows (Figure S2A).

**Figure 1:**
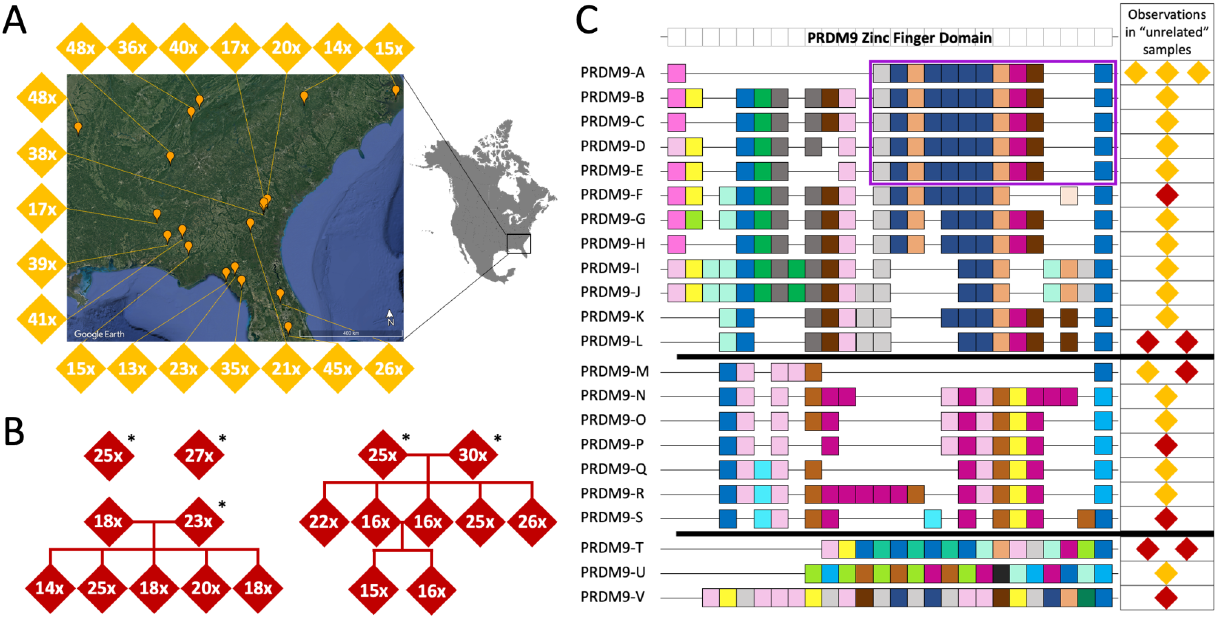
Genome sequences of corn snakes and PRDM9 zinc finger alleles in our samples. A) Sample collection locations for wild-caught individuals are shown for the 19 individuals depicted by a diamond. The number in each diamond indicates the mean fold-coverage of whole genome sequencing. B) The pedigree structures for samples from the colony, also including “unrelated” individuals, indicated with an asterisk. The number in each diamond indicates the mean fold-coverage of genome sequencing. C) PRDM9 zinc finger domain structure for 22 PRDM9 alleles, grouped and aligned by the similarity of their computationally-predicted binding affinity. Zinc fingers with distinct predictions for their binding affinities are shown in different colors; loosely, more similar colors represent zinc fingers with more similar computationally-predicted binding affinities (Figure S3). Each observation of a given allele is shown in the table; gold diamonds indicate wild samples and red diamonds colony samples. If the same allele was identified multiple times in closely related individuals, it is only shown once. The purple box highlights a succession of 11 zinc fingers (Shared 11-ZF) that are shared among five different alleles, including the only allele seen more than twice in the sample, PRDM9-A.

### PRDM9 diversity

The PRDM9 zinc finger has a minisatellite-like structure that makes amplifying, sequencing, and assembling this genomic region challenging (*26, 27*). To characterize the diversity of PRDM9 zinc finger alleles in our sample, we therefore used a combination of Sanger, Oxford Nanopore, and PacBio sequencing (see Methods). Ultimately, we identified 22 distinct PRDM9 alleles in 25 individuals (Figure 1C, Figure S3, Table S2). Strikingly, only four alleles are found more than once in our sample and no allele is present more than three times, reflecting the immense diversity of PRDM9 zinc finger alleles segregating in corn snakes (Figure 1C). Moreover, the zinc finger domains vary in their lengths and the identities of the residues in contact with DNA, leading to enormous variation in computationally predicted binding affinities (Figure S3). In contrast, the rest of the PRDM9 protein is conserved between mammals and snakes (Figure S4A). Rapid evolution of the residues that specify binding affinity is expected from a role of PRDM9 in directing recombination (*22, 23*).

### The population recombination rate increases both near PRDM9 binding sites and near promoter-like features

In our unrelated sample of corn snakes, expected pairwise linkage disequilibrium (LD) decays on the order of 10 kb (Figure S5). Given the levels of heterozygosity, this scale of LD decay suggests that the ratio of mutation to recombination is comparable to, or perhaps a bit lower than, what is found in humans and other species in which recombination rates have been reliably estimated from patterns of LD (*28*). Using the genetic variation found in the 24 unrelated individuals, we therefore proceeded to infer LD-based recombination maps, using the programs LDhelmet (*28*) and Pyrho (*29*). A comparison of the genetic maps obtained by the two approaches suggested that estimates based on LDhelmet may be more reliable (see Methods, Figure S6). In what follows, we therefore relied on results from LDHelmet, after excluding any 10 kb region in which the recombination rates estimated by the two methods differ by >10-fold (2.2% of the autosomal genome); qualitative results are similar if based on the Pyrho map instead (Figure S7). At a 10 kb scale, the resulting map has a mean estimated population recombination rate of 0.0068 per bp and a median of 0.0027 per bp (Figure S8A). As expected, recombination rates increase with GC content (Figure S8B).

LD-based inferences provide an estimate of the population recombination rate, which reflects an average over the many generations ancestral to the sample (*30*). In that regard, the immense diversity of PRDM9 zinc finger alleles presents an interpretative challenge, as each individual allele may be too rare or have been too short-lived to have left a discernible footprint in the decay of LD. Nonetheless, the set of computationally predicted PRDM9 binding sites is associated with a significant increase in the mean population recombination rate, in genomic regions far (>10kb) from a promoter-like feature (Figure 2A, Figure S9).

**Figure 2:**
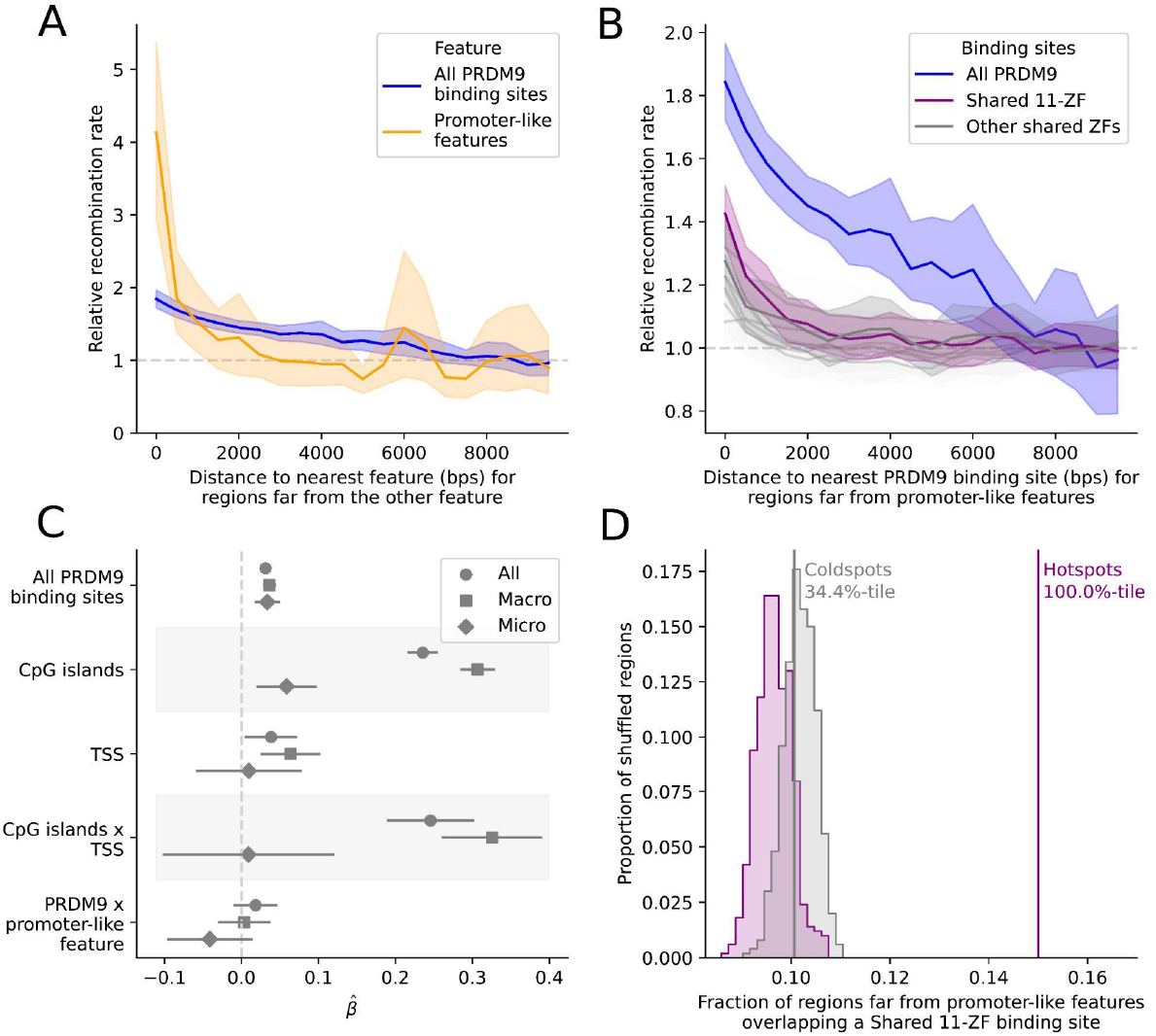
An increase in the population recombination rate is seen around *both* PRDM9 binding sites and promoter-like features. A) Mean population recombination rate in 100 bp windows as a function of distance to the nearest predicted PRDM9 binding site (blue) or promoter-like feature (orange). When considering one feature, we condition on windows >10 kb from the other feature; thus, when focusing on predicted PRDM9 binding sites, we only consider windows that are >10 kb from promoter-like features (i.e., TSS or CpG island). The recombination rate is relative to the mean rate 8-10 kb away. Shaded regions represent the central 95% confidence interval obtained by bootstrapping (see Methods). B) Mean population recombination in 100 bp windows as a function of distance from predicted binding sites for sets of zinc fingers shared among PRDM9 alleles. This plot is conditional on the windows being far from promoter-like features. The “Shared 11-ZF” allele shared among five PRDM9 alleles is shown in purple and the set of all PRDM9 alleles, equivalent to the curve in (A), is shown in blue. C) Point estimate and 95% CI for the coefficients of a linear model, in which the response variable is the (log) recombination rate in 1 kb windows (thinned to be 10 kb apart) and the predictors are the (binary) presence or absence of one or more predicted PRDM9 binding sites, TSS, or CpG islands. Covariates include the background recombination rate (1 Mb scale) and GC content (see Methods). Results are reported for data from the autosomes (circles), only scaffolds assigned to macrochromosomes (squares), and only microchromosomes (diamonds). D) Overlap of hotspots (purple) and matched coldspots (gray) far from a promoter-like feature with the predicted binding sites for the Shared 11-ZF allele. The observed values are shown with solid lines. The overlap expected by chance is shown with the shaded distribution and is based on 500 replicates, in which each hotspot was placed at random within 5 Mb of the original location conditional on there not being a gap in the genome sequence (see Methods). We note that while hotspots and coldspots are matched for base composition (see Methods), that need no longer be the case once we condition on them laying far from a promoter-like feature, driving the slight difference between the null distributions.

Despite their diversity, corn snake zinc finger alleles of PRDM9 sometimes share a subset of fingers in common (Figure 1C). To hone in on such shared features, we considered all sets of five or more consecutive fingers observed in at least five alleles (Figure S10). The predicted binding sites of each of these “shared zinc fingers” are associated with increased recombination, and in many cases, the decay with distance is steeper than it is for the union of binding sites from all zinc finger alleles (Figure 2B). Of particular note is the clear elevation in the mean rate around binding sites of a sequence of 11 zinc fingers that is found in five different PRDM9 alleles (henceforth, the “Shared 11-ZF” allele), including PRDM9-A, the most common allele in our sample (Figure 2B). Thus, despite the diversity of PRDM9 zinc finger alleles, patterns of LD support the use of PRDM9 binding in directing recombination in snakes.

Intriguingly, however, there is also a strong signal of increased recombination rate near promoter-like features that are far (>10 kb) from any predicted PRDM9 binding site (Figure 2A). The magnitude of the increase dwarfs that at PRDM9 but is in fact not directly comparable: CpG islands and TSS are readily identified compared to true PRDM9 binding sites and are relatively stable over evolutionary timescales. By contrast, PRDM9 binding sites vary by allele and are difficult to predict computationally (*2, 31, 32*). What is clear is that the signals of PRDM9 binding and promoter-like features are distinct.

In fact, the effects of the two features are largely independent: in a linear model, both putative PRDM9 binding sites and promoter-like features–notably CpG islands–are highly significant predictors of the population recombination rate in 1 kb bins (Figure 2C; p < 10^−22^ for PRDM9 binding sites and CpG islands and p=0.02 for TSS, see Methods), and their interaction term is not significantly different from 0 (p=0.18). The model further reveals an apparent difference between genomic regions on one of the eight macrochromosomes (defined as chromosomes >35 Mb in length) (*33*) versus one of the 10 microchromosomes (see Methods): despite a high density of genes on microchromosomes (*33*) there appears to be a much weaker effect of CpG islands and TSS, but a comparable effect of predicted PRDM9 binding sites (Figure 2C).

These findings stand in contrast to the conclusion reached in an analysis of LD patterns in the prairie rattlesnake (*Crotalus viridis)*, where, based on three of the zinc fingers of the *PRDM9* reference genome sequence, the authors concluded that PRDM9 directs recombination to promoter-like features (*34*). We re-analyzed their genomic data after collecting six complete prairie rattlesnake *PRDM9* alleles from three individuals (Figure S11). While we confirmed that rattlesnake PRDM9 alleles are predicted to bind GC rich genomic regions (Figure S12), such as CpG islands, we found separable effects of PRDM9 binding sites and promoter-like features on population recombination rates, as in corn snakes (Figure S13). In the course of our analyses, however, we noted that the prairie rattlesnake genome assembly may be unreliable around CpG islands (Figure S14), so a firm conclusion for this second snake species awaits further validation.

We also used the LD-based genetic map in corn snakes to infer the presence of 13,580 autosomal recombination hotspots, defined as genomic regions of >2kb with recombination rates at least 5-fold the background rate of the flanking regions (see Methods). Given these criteria, the median estimated heat is 8.6-fold and the median estimated length ∼3 kb (Figure S15). Hotspots far (>10 kb) from promoter-like features overlap with predicted PRDM9 binding sites more often than expected by chance (p=0.002, as assessed by reshuffling hotspot locations 500 times within 5 Mb of the starting location to control for background recombination rate; Figure S16A). In contrast, GC-matched coldspots far from promoter-like features overlap with PRDM9 binding sites no more than expected by chance (p=0.61, Figure S16A).

Thus, the overlap of hotspots with predicted PRDM9 binding sites cannot be explained simply by local base composition effects. The same finding is obtained for sites predicted to be bound by the Shared 11-ZF allele (p=0.002, Figure 2C). Analogously, hotspots overlap promoter-like features that are far from any PRDM9 binding site significantly more than expected by chance (p=0.002), when the same is not true for GC-matched coldspots (p=0.34, Figure S16B). Thus, as expected from the analysis of population recombination rates, hotspot locations inferred from LD are associated with both predicted PRDM9 binding sites and promoter-like features.

### Divergence at PRDM9 binding sites

Because the chromosome on which a DSB occurs is repaired based on information from its homolog, recombination converts strong PRDM9 binding sites to weaker PRDM9 binding sites in heterozygotes (*35, 36*). In the absence of any countervailing force, this process leads to the more rapid loss of PRDM9 binding sites over evolutionary time than expected under neutrality (*3, 37*). To look for a signal of this phenomenon in the lineage ancestral to corn snake, we used a multi-species whole genome alignment that we had previously generated, which includes corn snake and the closely related black rat snake, *Pantherophis obsoletus* (*38*). First, we focused on predicted binding sites of the Shared-11ZF allele, as it shows a clear association with increased population recombination rates, suggesting it may be old enough to also have left a footprint in divergence data. There are more losses than gains of the binding sites of this allele in the corn snake lineage, but the difference is not statistically significant (Figure S17). A similar excess of losses over gains is seen for 16 of the 22 PRDM9 alleles (if we consider them to be independent, p=0.019, using a one-tailed Wilcoxon signed-rank test). Next, we identified eight kmers significantly over-represented in recombination hotspots relative to coldspots (see Methods); for seven of the eight, there were more losses than gains in the corn snake lineage (p=0.007, Figure 3A). Moreover, three of the kmers are close matches to subsets of the binding motifs of extant PRDM9 alleles (Figure S18). Together, these results indicate that putative PRDM9 binding motifs are being lost on the corn snake lineage faster than expected by chance, lending further support to the role of PRDM9 binding in specifying DSB locations.

**Figure 3:**
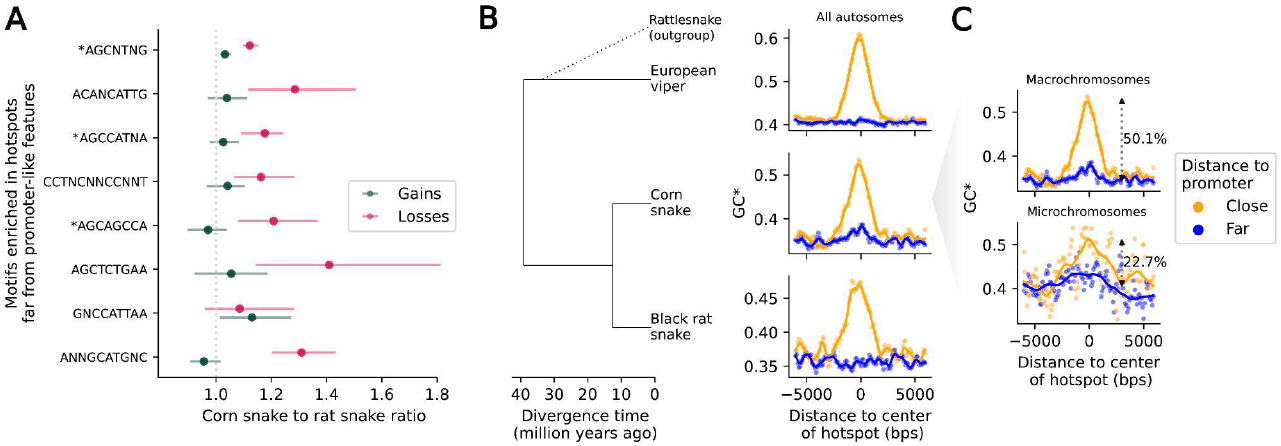
Footprints of recombination in divergence data. A) The ratio of losses in the corn snake lineage relative to the black rat snake (in magenta) and the ratio of gains in the corn snake lineage relative to the black rat snake (in green), for motifs enriched in recombination hotspots far from promoter-like features (>10 kb), ordered by motif enrichment significance (top to bottom). The 95% confidence intervals are obtained by bootstrapping over 5 Mb regions with at least one gain or loss event in either of the two lineages. B) Increased flux to GC (GC*) as a function of distance from corn snake autosomal hotspots, in sliding windows of 500 bp with a 100 bp offset, for the lineages leading to European viper (top), corn snake (middle) and black rat snake (bottom). Note that rattlesnakes, a sister species to European viper, were included only to infer the ancestral state in substitutions (see Methods). GC* around hotspots that are close (<500 bp) to a promoter-like feature (orange) and far from any promoter-like feature (blue) are shown. Local regression curves are shown for a span of 0.05. C) GC* in the lineage leading to corn snakes, for hotspots in macro-(top) and microchromosomes (bottom), using the same color scheme as in B. Local regression curves are shown for a span of 0.05 and 0.2 in macro- and microchromosomes, respectively. The percentage increase in the mean GC* for sites >3 kb away from the center of the hotspot relative to that within 200 bps of the center are shown.

### Conservation of hotspots at promoter-like features

One implication of these findings is that only a subset of hotspots–those dependent on PRDM9–are expected to evolve rapidly and to differ among closely related snake species, whereas hotspots near promoters may be largely conserved, as they are in species lacking PRDM9 (*16, 39*). To evaluate this possibility, we relied on GC* to quantify the excess flux towards GC-substitutions in a lineage, which results from GC-biased gene conversion associated with recombination (*40*). We found evidence for increased GC* at corn snake hotspots near promoter-like features in the lineage leading to corn snakes as well as in orthologous regions in the black rat snake and european viper branches (Figure 3B), when no such elevation is seen for coldspots (Figure S19). Considering that the three species last shared a common ancestor more than 30 MYA (*41*) (Figure 3B), these findings point to a remarkable stability of recombination hotspots at promoter-like features, as previously reported in birds (*16*) and yeast (*39*).

In contrast, recombination hotspots that are far from promoter-like features, a subset of which are likely specified by PRDM9 binding in corn snakes, show a much weaker signal of elevated GC* in the corn snake branch, and no discernible GC* peaks in the other two snake lineages (Figure 3B). This observation is consistent with corn snakes having two classes of hotspots: those near promoter-like features (and possibly other sites of H3K4me3), which are conserved among distantly related species, and those at PRDM9 binding sites, which are rapidly repositioned because of changes in the binding affinity of PRDM9. In that regard, it is interesting to note that the signal of GC* at hotspots overlapping promoter-like features is less pronounced on micro- than macrochromosomes (Figure 3C), consistent with a reduced use of promoter-like features on microchromosomes (Figure 2C).

### Crossovers occur at PRDM9 binding sites and CpG islands

LD-based genetic maps represent a mixture of the effects of many different PRDM9 zinc finger alleles, past and present, and LD-based estimates can be biased in the presence of population structure or changes in population size (*28, 42, 43*) as well as by subtle errors in the reference genome (Figure S14). We therefore sought to confirm our findings of a dual usage of PRDM9 and promoter-like features using a more direct and robust approach. To that end, we collected genome sequences from two corn snake pedigrees with five F1s each, as well as two F2s from one of the pedigrees, thereby obtaining information about 24 meioses (Figure 1B).

Calling crossovers from phase switches of informative markers (following (*44*); see Methods), we identified 324 crossover events, of which 172 could be delimited to within an interval of <5 kb. 177 occurred in mothers and 147 in fathers, a sex ratio not significantly different from 1:1 (by a two-tailed Binomial test, p=0.10). At the scaffold level, population recombination rates and crossover rates are highly correlated (Figure S20B) and, as expected, crossover events overlap LD hotspots more often than expected by chance (Figure S20A, p=0.0003). Moreover, the observed overlap (19.2%) is in good agreement with what we expect based on the estimated widths and heats of hotspots (18.4%, see Methods).

Crossovers (delimited to intervals of less than 5 kb) far from promoter-like features overlap predicted PRDM9 binding sites of the parent in which they occurred more frequently than expected by chance (p = 0.0006, Figure 4A). Inversely, crossovers far from the predicted PRDM9 binding sites of the focal parent overlap promoter-like features more often than expected by chance (p = 0.007, Figure 4B). Qualitative conclusions are the same if a larger set of 239 crossovers with a resolution <20 kb are used instead (Figure S21). Moreover, there are no significant differences between sexes in the usage of PRDM9 binding sites (by Fisher’s exact test, p=0.85) or promoter-like features (p=0.61, Figure S22). Thus, the analysis of crossovers in pedigrees confirms our findings based on LD: both promoter-like features and PRDM9 binding sites are used to direct recombination in corn snakes, seemingly in both sexes.

**Figure 4:**
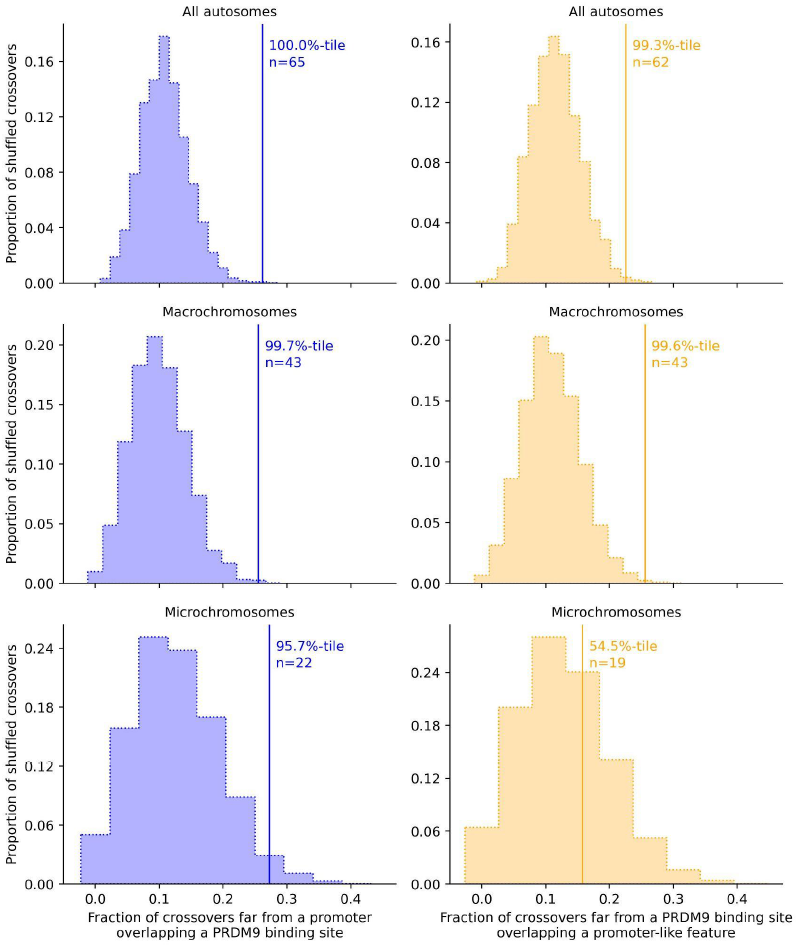
Crossovers identified from pedigree resequencing overlap both predicted PRDM9 binding sites and promoter-like features significantly more often than expected by chance in corn snakes. Shown is the overlap for crossovers resolved to within 5 kb. In the first column (blue) is the overlap of crossovers with the predicted binding sites of the PRDM9 alleles carried by the parent in which the crossover is inferred to have occurred, for binding sites far from a promoter-like feature (>10 kb). In the second column (orange) is the overlap of crossovers with promoter-like features far (>10kb) from the PRDM9 binding sites of alleles carried by the parents. The solid line is the observed overlap for the subset of crossovers, n, that satisfy the criteria. The frequency distribution represents the overlap for 3000 sets of simulated crossovers obtained by placing the observed interval lengths down at random within 5 Mb of the original crossover interval, conditional on it containing at least two informative markers and there not being a gap in the genome sequence at that location (see Methods). The rank of the observation relative to realizations under the null is given as a percentile (*47*). The three rows present results for all scaffolds, only those assigned to macrochromosomes and only those assigned to microchromosomes (see Methods).

Controlling for distance to the other feature and accounting for overlap by chance, we estimate that across the autosomes, 15.4% of crossovers (95% CI: 7.7-21.5%) overlap with PRDM9 binding sites whereas 11.3% (95% CI: 3.2-17.7%) overlap promoter-like features (see Methods). The estimated overlap with PRDM9 binding sites is almost certainly an underestimate, given the difficulty of computationally predicting binding sites. The estimate for promoter-like features is likely to be less downwardly biased. Nonetheless, if we assume that promoter-like features are not causal but only associated with a causal feature (e.g., sites of H3K4me3), the estimate for promoter-like features likely only captures a fraction of crossovers that occur through PRDM9-independent mechanisms.

As in analyses of LD, we once again find evidence that promoter-like features are not used to the same extent on microchromosomes: in contrast to what is seen for macrochromosomes (p=0.004, Figure 4), there is no significant overlap of crossovers with predicted promoter features far from PRDM9 binding sites (p = 0.45, Figure 4). This difference persists when accounting for the reduced number of crossovers on microchromosomes (Figure S23). Thus, a smaller proportion of crossovers are associated with promoter-like features on micro- than macrochromosomes, consistent with the evidence from the linear model (Figure 2C) and patterns of GC* (Figure 3C). One possibility is that the proportion of DSBs at promoter-like features versus PRDM9 binding sites is similar, but PRDM9 binding sites are more likely to result in a crossover (rather than a non-crossover) on microchromosomes. Alternatively, the greater use of PRDM9-independent hotspots on later-replicating macrochromosomes (*45*) could reflect a relative delay in DSB formation, as proposed for the pseudo-autosomal region in mice (*46*).

### Changes downstream of PRDM9 are likely to underlie the different recombination landscape of snakes

In mammals studied to date, there appear to be two distinct strategies to directing recombination to the genome: either the vast majority of DSBs occur at PRDM9 binding sites or, in species that lack an intact PRDM9, recombination rates are elevated in promoter regions and other sites of H3K4me3. In contrast, in corn snakes, recombination is more evenly distributed between PRDM9 binding sites and promoter-like features. Since the key catalytic residues in the SET domain of PRDM9 are unchanged between mammals and snakes (Figure S4A), it seems likely that PRDM9 makes the same histone modifications and that changes to downstream partners instead underlie the use of PRDM9-independent DSBs in corn snakes.

Among possible candidates are the two other genes present in almost all vertebrate species that carry PRDM9, including corn snakes, and absent in most lineages without an intact PRDM9 (*13*). The first, ZCWPW1, binds H3K4me3 and H3K36me3 marks through its zinc finger CW domain (zf-CW) and PWWP domain, respectively (*48*–*50*). In mice, ZCWPW1 is essential for the efficient repair of PRDM9-dependent DSBs, but does not affect their localization (*51*). The second gene is its paralog ZCWPW2, which also has zf-CW and PWWP domains (*13*). As we show in an *in vitro* binding assay, the ZCWPW2 protein from mice binds both marks, more strongly so when the two are in combination than when either one is alone (Figure 5A). Based on its domain structure and its co-evolution with *PRDM9*, we previously suggested that ZCWPW2 may be a missing link between PRDM9 binding and the recruitment of the DSB machinery (*13*). If so, changes to ZCWPW2 could underlie the greater use of promoter-like features in corn snakes.

**Figure 5:**
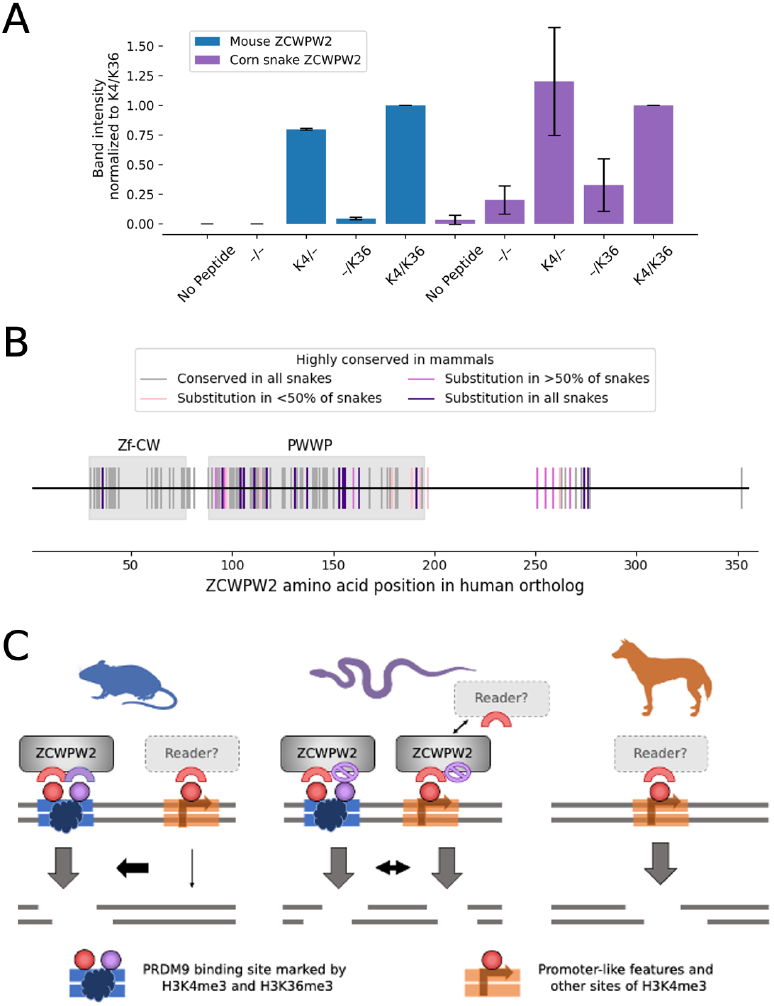
Possible role for ZCWPW2 in the tug of war between promoter-like features and PRDM9. A) In vitro binding affinity of mouse (blue) and snake (purple) ZCWPW2 for histone peptides methylated with H3K4me3 (K4) and H3K36me3 (K36). Plotted are the band intensities of mouse and snake HA-tagged ZCWPW2 in Western Blotting following H3-peptide pull-down (see Methods). For each experimental replicate, the band intensities were normalized to the band intensity of Zcwpw2 pulled down using the K4/K36 labeled peptide. The experiment was performed once using corn-snake Zcwpw2-HA alone, and twice using both mouse and corn-snake Zcwpw2-HA. Whiskers denote two standard deviations. B) Conservation of ZCWPW2 amino acids among 137 mammalian and nine snake species. Each tick mark denotes a site at which >95% mammals carry the same amino acid. Amino acids shown as gray ticks are completely conserved in the nine snakes; pink are conserved in all but 1-4 snakes, maroon are changed in 5-8 snakes, and purple amino acids are changed in all 9 snakes. Of the 20 sites in the Zf-CW domain that are highly conserved in mammals, there is a substitution in all snakes at 1 site (5%), whereas of 58 such sites in the PWWP, 12 (21%) have changed in all snakes. C) Proposed schema by which decreased binding affinity of ZCWPW2 to H3K4me36 in snakes could lead to DSBs and resulting crossovers at both PRDM9 binding sites and promoter-like features, in contrast to their mutually exclusive use in mammals studied to date.

*ZCWPW2* is evolving under purifying selection in snakes (dN/dS=0.54, p=2×10^−5^; estimated using PAML (*52*)), including in the corn snake lineage (p=0.018), indicating that it plays a functional role in this taxon (Table S3). Yet the protein differs at a number of amino acid positions that are highly conserved across 137 mammalian species with *PRDM9*–particularly in the PWWP domain that binds H3K36me3 (Figure 5B). The substitutions in the PWWP domain of ZCWPW2 may be functionally important: in our *in vitro* binding experiment, the corn snake ZCWPW2 is not detectably better at binding both marks together than H3K4me3 alone (Figure 5A). A possible explanation is that the changes to the PWWP domain of snakes lead to a comparable affinity of ZCWPW2 for H3 at PRDM9 binding sites with dual H3K4me3/H3K36me3 marks and for single H3K4me3 marks found in other locations. If so, the DSB machinery may be recruited by ZCWPW2 both to sites of PRDM9 binding and to promoter-like regions, or the PRDM9/ZCWPW2 system may be more readily outcompeted by an alternative recruitment mechanism to promoter-like regions (Figure 5C).

While these findings are intriguing, changes to *ZCWPW2* need not be the original cause, or the only cause, of the shift in the recombination landscape towards promoter-like features in corn snakes. Notably, the dual usage of PRDM9 binding sites and promoter-like features is reminiscent of what is seen in *Ankrd31*-/-mice, in which DSBs are distributed between PRDM9-dependent hotspots and promoter regions (*46, 53, 54*). While *ANKRD31* is present in snakes, including corn snakes, and is evolving under purifying selection (Table S3), more subtle changes to the protein may act as hypomorphic alleles (*54*). Regardless of the cause, changes in DSB localization are likely to also have been accompanied by remodeling of later stages of recombination; in this regard, it is interesting to note that there are a number of substitutions in the PWWP domain of *ZCWPW1* in snakes as well (Figure S4B).

### A tug of war between the histone modifications made by PRDM9 and H3K4me3 sites elsewhere in the genome in vertebrates

In light of these results, we propose that across vertebrates (and possibly beyond; (*12, 15*)), the distribution of recombination events along the genome results from a competition between the recruitment of the DSB machinery to histone marks laid down by PRDM9 and the recruitment to sites of H3K4me3 unrelated to recombination (*7, 9*). In mammals with *PRDM9* studied to date, the competition is won almost entirely by PRDM9, and the alternative mechanism of DSB initiation is revealed only in knock-outs or in hybrid mice, in which repair at PRDM9-dependent hotspots is compromised (*55*). In corn snakes, in contrast, PRDM9-dependent recombination is not as dominant a mechanism and promoter-like features are used to a greater extent, especially on macrochromosomes. The explanation may lie in changes to the binding affinity of ZCWPW2 or in alternative molecular paths. In any case, our findings highlight that uses of PRDM9 and promoter-like features in vertebrates are not mutually-exclusive, and instead reflect a tug of war between competing signals that plays out differently along the genome and has more lopsided outcomes in some species than others.

These findings imply that the recombination landscapes of vertebrates may lie in a continuum between the quasi-exclusive use of PRDM9-dependent hotspots on the one hand and the reliance on signals associated with promoter-regions on the other, with implications for evolutionary genetics as well as genome editing (*56*–*58*). Moreover, they indicate that natural selection can shape the recombination landscape of a species by tuning the extent to which these two strategies are used, likely through dials downstream of PRDM9.

## Acknowledgements

We thank Jeff Spence for helpful discussions, as well as Akanksha Bhat, Djivan Prentout and other members of the Andolfatto, Przeworski, and Sella labs; Ed Myers and Frank Burbrink for providing samples of corn snakes and rattlesnakes; and Yun Deng for help with initial attempts to sequence PRDM9 alleles. An unpublished genome assembly for corn snakes was used with permission from the DNA Zoo Consortium (dnazoo.org).

## Supplementary Materials

Materials and Methods

Figs. S1 to S27

Tables S1 to S7

## References

1. B. de Massy, Initiation of meiotic recombination: how and where? Conservation and specificities among eukaryotes. Annu. Rev. Genet. 47, 563–599 (2013).

2. F. Baudat, J. Buard, C. Grey, A. Fledel-Alon, C. Ober, M. Przeworski, G. Coop, B. de Massy, PRDM9 is a major determinant of meiotic recombination hotspots in humans and mice. Science. 327, 836–840 (2010).

3. S. Myers, R. Bowden, A. Tumian, R. E. Bontrop, C. Freeman, T. S. MacFie, G. McVean, P. Donnelly, Drive against hotspot motifs in primates implicates the PRDM9 gene in meiotic recombination. Science. 327, 876–879 (2010).

4. Y. Zhou, B. Shen, J. Jiang, A. Padhi, K.-E. Park, A. Oswalt, C. G. Sattler, B. P. Telugu, H. Chen, J. B. Cole, G. E. Liu, L. Ma, Construction of PRDM9 allele-specific recombination maps in cattle using large-scale pedigree analysis and genome-wide single sperm genomics. DNA Res. 25, 183–194 (2018).

5. L. S. Stevison, A. E. Woerner, J. M. Kidd, J. L. Kelley, K. R. Veeramah, K. F. McManus, Great Ape Genome Project, C. D. Bustamante, M. F. Hammer, J. D. Wall, The Time Scale of Recombination Rate Evolution in Great Apes. Mol. Biol. Evol. 33, 928–945 (2016).

6. O. Mihola, V. Landa, F. Pratto, K. Brick, T. Kobets, F. Kusari, S. Gasic, F. Smagulova, C. Grey, P. Flachs, V. Gergelits, K. Tresnak, J. Silhavy, P. Mlejnek, R. D. Camerini-Otero, M. Pravenec, G. V. Petukhova, Z. Trachtulec, Rat PRDM9 shapes recombination landscapes, duration of meiosis, gametogenesis, and age of fertility. BMC Biol. 19, 86 (2021).

7. K. Brick, F. Smagulova, P. Khil, R. D. Camerini-Otero, G. V. Petukhova, Genetic recombination is directed away from functional genomic elements in mice. Nature. 485, 642–645 (2012).

8. B. Davies, E. Hatton, N. Altemose, J. G. Hussin, F. Pratto, G. Zhang, A. G. Hinch, D. Moralli, D. Biggs, R. Diaz, C. Preece, R. Li, E. Bitoun, K. Brick, C. M. Green, R. D. Camerini-Otero, S. R. Myers, P. Donnelly, Re-engineering the zinc fingers of PRDM9 reverses hybrid sterility in mice. Nature. 530, 171–176 (2016).

9. B. Diagouraga, J. A. J. Clément, L. Duret, J. Kadlec, B. de Massy, F. Baudat, PRDM9 Methyltransferase Activity Is Essential for Meiotic DNA Double-Strand Break Formation at Its Binding Sites. Mol. Cell. 69, 853–865.e6 (2018).

10. Y. Imai, F. Baudat, M. Taillepierre, M. Stanzione, A. Toth, B. de Massy, The PRDM9 KRAB domain is required for meiosis and involved in protein interactions. Chromosoma. 126, 681–695 (2017).

11. S. Thibault-Sennett, Q. Yu, F. Smagulova, J. Cloutier, K. Brick, R. D. Camerini-Otero, G. V. Petukhova, Interrogating the Functions of PRDM9 Domains in Meiosis. Genetics. 209, 475–487 (2018).

12. Z. Baker, M. Schumer, Y. Haba, L. Bashkirova, C. Holland, G. G. Rosenthal, M. Przeworski, Repeated losses of PRDM9-directed recombination despite the conservation of PRDM9 across vertebrates. Elife. 6, e24133 (2017).

13. M. I. A. Cavassim, Z. Baker, C. Hoge, M. H. Schierup, M. Schumer, M. Przeworski, PRDM9 losses in vertebrates are coupled to those of paralogs ZCWPW1 and ZCWPW2. Proceedings of the National Academy of Sciences. 119, e2114401119 (2022).

14. V. Muñoz-Fuentes, A. Di Rienzo, C. Vilà, Prdm9, a major determinant of meiotic recombination hotspots, is not functional in dogs and their wild relatives, wolves and coyotes. PLoS One. 6, e25498 (2011).

15. P. L. Oliver, L. Goodstadt, J. J. Bayes, Z. Birtle, K. C. Roach, N. Phadnis, S. A. Beatson, G. Lunter, H. S. Malik, C. P. Ponting, Accelerated evolution of the Prdm9 speciation gene across diverse metazoan taxa. PLoS Genet. 5, e1000753 (2009).

16. S. Singhal, E. M. Leffler, K. Sannareddy, I. Turner, O. Venn, D. M. Hooper, A. I. Strand, Q. Li, B. Raney, C. N. Balakrishnan, S. C. Griffith, G. McVean, M. Przeworski, Stable recombination hotspots in birds. Science. 350, 928–932 (2015).

17. A. Auton, Y. Rui Li, J. Kidd, K. Oliveira, J. Nadel, J. K. Holloway, J. J. Hayward, P. E. Cohen, J. M. Greally, J. Wang, C. D. Bustamante, A. R. Boyko, Genetic recombination is targeted towards gene promoter regions in dogs. PLoS Genet. 9, e1003984 (2013).

18. N. R. Powers, E. D. Parvanov, C. L. Baker, M. Walker, P. M. Petkov, K. Paigen, The Meiotic Recombination Activator PRDM9 Trimethylates Both H3K36 and H3K4 at Recombination Hotspots In Vivo. PLoS Genet. 12, e1006146 (2016).

19. A. L. Hughes, J. R. Kelley, R. J. Klose, Understanding the interplay between CpG island-associated gene promoters and H3K4 methylation. Biochim. Biophys. Acta Gene Regul. Mech. 1863, 194567 (2020).

20. V. M. Narasimhan, K. A. Hunt, D. Mason, C. L. Baker, K. J. Karczewski, M. R. Barnes, A. H. Barnett, C. Bates, S. Bellary, N. A. Bockett, K. Giorda, C. J. Griffiths, H. Hemingway, Z. Jia, M. A. Kelly, H. A. Khawaja, M. Lek, S. McCarthy, R. McEachan, A. O’Donnell-Luria, K. Paigen, C. A. Parisinos, E. Sheridan, L. Southgate, L. Tee, M. Thomas, Y. Xue, M. Schnall-Levin, P. M. Petkov, C. Tyler-Smith, E. R. Maher, R. C. Trembath, D. G. MacArthur, J. Wright, R. Durbin, D. A. van Heel, Health and population effects of rare gene knockouts in adult humans with related parents. Science. 352, 474–477 (2016).

21. O. Mihola, F. Pratto, K. Brick, E. Linhartova, T. Kobets, P. Flachs, C. L. Baker, R. Sedlacek, K. Paigen, P. M. Petkov, R. D. Camerini-Otero, Z. Trachtulec, Histone methyltransferase PRDM9 is not essential for meiosis in male mice. Genome Res. 29, 1078–1086 (2019).

22. Z. Baker, M. Przeworski, G. Sella, Down the Penrose stairs: How selection for fewer recombination hotspots maintains their existence. bioRxiv (2022), p. 2022.09.27.509707.

23. A. Genestier, L. Duret, N. Lartillot, Bridging the gap between the evolutionary dynamics and the molecular mechanisms of meiosis: a model based exploration of the PRDM9 intra-genomic Red Queen. bioRxiv (2023), p. 2023.03.08.531712.

24. A. Ullate-Agote, I. Burgelin, A. Debry, C. Langrez, F. Montange, R. Peraldi, J. Daraspe, H. Kaessmann, M. C. Milinkovitch, A. C. Tzika, Genome mapping of a LYST mutation in corn snakes indicates that vertebrate chromatophore vesicles are lysosome-related organelles. Proceedings of the National Academy of Sciences. 117, 26307–26317 (2020).

25. B. S. Weir, C. C. Cockerham, ESTIMATING F-STATISTICS FOR THE ANALYSIS OF POPULATION STRUCTURE. Evolution. 38, 1358–1370 (1984).

26. J. J. Schwartz, D. J. Roach, J. H. Thomas, J. Shendure, Primate evolution of the recombination regulator PRDM9. Nat. Commun. 5, 4370 (2014).

27. J. Buard, E. Rivals, D. Dunoyer de Segonzac, C. Garres, P. Caminade, B. de Massy, P. Boursot, Diversity of Prdm9 zinc finger array in wild mice unravels new facets of the evolutionary turnover of this coding minisatellite. PLoS One. 9, e85021 (2014).

28. A. H. Chan, P. A. Jenkins, Y. S. Song, Genome-wide fine-scale recombination rate variation in Drosophila melanogaster. PLoS Genet. 8, e1003090 (2012).

29. J. P. Spence, Y. S. Song, Inference and analysis of population-specific fine-scale recombination maps across 26 diverse human populations. Science Advances. 5, eaaw9206 (2019).

30. G. Hellenthal, M. Stephens, Insights into recombination from population genetic variation. Curr. Opin. Genet. Dev. 16, 565–572 (2006).

31. N. Altemose, N. Noor, E. Bitoun, A. Tumian, M. Imbeault, J. R. Chapman, A. R. Aricescu, S. R. Myers, A map of human PRDM9 binding provides evidence for novel behaviors of PRDM9 and other zinc-finger proteins in meiosis. Elife. 6, e28383 (2017).

32. A. V. Persikov, M. Singh, De novo prediction of DNA-binding specificities for Cys2His2 zinc finger proteins. Nucleic Acids Res. 42, 97–108 (2014).

33. P. D. Waters, H. R. Patel, A. Ruiz-Herrera, L. Álvarez-González, N. C. Lister, O. Simakov, T. Ezaz, P. Kaur, C. Frere, F. Grützner, A. Georges, J. A. M. Graves, Microchromosomes are building blocks of bird, reptile, and mammal chromosomes. Proc. Natl. Acad. Sci. U. S. A. 118 (2021), doi:10.1073/pnas.2112494118.

34. D. R. Schield, G. I. M. Pasquesi, B. W. Perry, R. H. Adams, Z. L. Nikolakis, A. K. Westfall, R. W. Orton, J. M. Meik, S. P. Mackessy, T. A. Castoe, Snake Recombination Landscapes Are Concentrated in Functional Regions despite PRDM9. Mol. Biol. Evol. 37, 1272–1294 (2020).

35. T. Nagylaki, T. D. Petes, Intrachromosomal gene conversion and the maintenance of sequence homogeneity among repeated genes. Genetics. 100, 315–337 (1982).

36. A. Boulton, R. S. Myers, R. J. Redfield, The hotspot conversion paradox and the evolution of meiotic recombination. Proceedings of the National Academy of Sciences. 94, 8058–8063 (1997).

37. G. Coop, S. R. Myers, Live hot, die young: transmission distortion in recombination hotspots. PLoS Genet. 3, e35 (2007).

38. M. de Manuel, F. L. Wu, M. Przeworski, A paternal bias in germline mutation is widespread in amniotes and can arise independently of cell division numbers. Elife. 11 (2022), doi:10.7554/eLife.80008.

39. I. Lam, S. Keeney, Nonparadoxical evolutionary stability of the recombination initiation landscape in yeast. Science. 350, 932–937 (2015).

40. J. Meunier, L. Duret, Recombination Drives the Evolution of GC-Content in the Human Genome. Mol. Biol. Evol. 21, 984–990 (2004).

41. S. Kumar, G. Stecher, M. Suleski, S. B. Hedges, TimeTree: A Resource for Timelines, Timetrees, and Divergence Times. Mol. Biol. Evol. 34, 1812–1819 (2017).

42. M. P. H. Stumpf, G. A. T. McVean, Estimating recombination rates from population-genetic data. Nat. Rev. Genet. 4, 959–968 (2003).

43. A. G. Hinch, A. Tandon, N. Patterson, Y. Song, N. Rohland, C. D. Palmer, G. K. Chen, K. Wang, S. G. Buxbaum, E. L. Akylbekova, M. C. Aldrich, C. B. Ambrosone, C. Amos, E. V. Bandera, S. I. Berndt, L. Bernstein, W. J. Blot, C. H. Bock, E. Boerwinkle, Q. Cai, N. Caporaso, G. Casey, L. A. Cupples, S. L. Deming, W. R. Diver, J. Divers, M. Fornage, E. M. Gillanders, J. Glessner, C. C. Harris, J. J. Hu, S. A. Ingles, W. Isaacs, E. M. John, W. H. L. Kao, B. Keating, R. A. Kittles, L. N. Kolonel, E. Larkin, L. Le Marchand, L. H. McNeill, R. C. Millikan, A. Murphy, S. Musani, C. Neslund-Dudas, S. Nyante, G. J. Papanicolaou, M. F. Press, B. M. Psaty, A. P. Reiner, S. S. Rich, J. L. Rodriguez-Gil, J. I. Rotter, B. A. Rybicki, A. G. Schwartz, L. B. Signorello, M. Spitz, S. S. Strom, M. J. Thun, M. A. Tucker, Z. Wang, J. K. Wiencke, J. S. Witte, M. Wrensch, X. Wu, Y. Yamamura, K. A. Zanetti, W. Zheng, R. G. Ziegler, X. Zhu, S. Redline, J. N. Hirschhorn, B. E. Henderson, H. A. Taylor Jr, A. L. Price, H. Hakonarson, S. J. Chanock, C. A. Haiman, J. G. Wilson, D. Reich, S. R. Myers, The landscape of recombination in African Americans. Nature. 476, 170–175 (2011).

44. G. Coop, X. Wen, C. Ober, J. K. Pritchard, M. Przeworski, High-resolution mapping of crossovers reveals extensive variation in fine-scale recombination patterns among humans. Science. 319, 1395–1398 (2008).

45. K. Srikulnath, S. F. Ahmad, W. Singchat, T. Panthum, Why Do Some Vertebrates Have Microchromosomes? Cells. 10 (2021), doi:10.3390/cells10092182.

46. F. Papanikos, J. A. J. Clément, E. Testa, R. Ravindranathan, C. Grey, I. Dereli, A. Bondarieva, S. Valerio-Cabrera, M. Stanzione, A. Schleiffer, P. Jansa, D. Lustyk, J.-F. Fei, I. R. Adams, J. Forejt, M. Barchi, B. de Massy, A. Toth, Mouse ANKRD31 Regulates Spatiotemporal Patterning of Meiotic Recombination Initiation and Ensures Recombination between X and Y Sex Chromosomes. Mol. Cell. 74, 1069–1085.e11 (2019).

47. B. V. North, D. Curtis, P. C. Sham, A note on the calculation of empirical P values from Monte Carlo procedures. Am. J. Hum. Genet. 71, 439–441 (2002).

48. T. Huang, S. Yuan, L. Gao, M. Li, X. Yu, J. Zhan, Y. Yin, C. Liu, C. Zhang, G. Lu, W. Li, J. Liu, Z.-J. Chen, H. Liu, The histone modification reader ZCWPW1 links histone methylation to PRDM9-induced double-strand break repair. Elife. 9 (2020), doi:10.7554/eLife.53459.

49. M. Mahgoub, J. Paiano, M. Bruno, W. Wu, S. Pathuri, X. Zhang, S. Ralls, X. Cheng, A. Nussenzweig, T. S. Macfarlan, Dual histone methyl reader ZCWPW1 facilitates repair of meiotic double strand breaks in male mice. Elife. 9, e53360 (2020).

50. D. Wells, E. Bitoun, D. Moralli, G. Zhang, A. Hinch, J. Jankowska, P. Donnelly, C. Green, S. R. Myers, ZCWPW1 is recruited to recombination hotspots by PRDM9 and is essential for meiotic double strand break repair. Elife. 9, e53392 (2020).

51. E. Damm, L. Odenthal-Hesse, Orchestrating recombination initiation in mice and men. Curr. Top. Dev. Biol. 151, 27–42 (2023).

52. Z. Yang, PAML 4: phylogenetic analysis by maximum likelihood. Mol. Biol. Evol. 24, 1586–1591 (2007).

53. M. Boekhout, M. E. Karasu, J. Wang, L. Acquaviva, F. Pratto, K. Brick, D. Y. Eng, J. Xu, R. D. Camerini-Otero, D. J. Patel, S. Keeney, REC114 Partner ANKRD31 Controls Number, Timing, and Location of Meiotic DNA Breaks. Mol. Cell. 74, 1053–1068.e8 (2019).

54. J. Xu, T. Li, S. Kim, M. Boekhout, S. Keeney, Essential roles of the ANKRD31-REC114 interaction in meiotic recombination and mouse spermatogenesis. bioRxiv (2023), doi:10.1101/2023.04.27.538541.

55. F. Smagulova, K. Brick, Y. Pu, R. D. Camerini-Otero, G. V. Petukhova, The evolutionary turnover of recombination hot spots contributes to speciation in mice. Genes Dev. 30, 266–280 (2016).

56. M. Schumer, C. Xu, D. L. Powell, A. Durvasula, L. Skov, C. Holland, J. C. Blazier, S. Sankararaman, P. Andolfatto, G. G. Rosenthal, M. Przeworski, Natural selection interacts with recombination to shape the evolution of hybrid genomes. Science. 360, 656–660 (2018).

57. E. Chen, E. Lin-Shiao, M. Trinidad, M. S. Doost, D. Colognori, J. A. Doudna, Decorating chromatin for enhanced genome editing using CRISPR-Cas9. Proceedings of the National Academy of Sciences. 119, e2204259119 (2022).

58. C. Veller, N. B. Edelman, P. Muralidhar, M. A. Nowak, Recombination and selection against introgressed DNA. Evolution. 77, 1131–1144 (2023).

