## Supplemental Materials for "Patterns of recombination in snakes reveal a tug of war between PRDM9 and promoter-like features"

### Supplemental information

### Methods

**1 Corn snake genome assembly and annotation** **3**

### 1 Corn snake genome assembly and annotation

We sought to improve the *Pantherophis guttatus* genome assembly and annotation published in Ullate-Agote *et. al.* (2020), using long-read and whole-genome bisulfite sequencing data[1].

#### 1.1 Animal experimentation

The colony of corn snakes was housed and bred at the LANE animal facility (University of Geneva) running under a veterinary cantonal permit. The individuals were sampled following Swiss law regulations and under experimentation permits GE24/33145 approved by the Geneva cantonal authorities.

#### 1.2 Genome assembly

The Ullate-Agote (2020)[1] genome assembly is 1.71 Gb long and was constructed using a combination of 10X Chromium reads and BioNano optical maps; it has a final scaffold N50 of 16.8 Mb and contig N50 of 38 kb. To improve on this assembly, we performed PacBio whole genome sequencing using a sample from a corn snake colony maintained by one of us (A.T.) at the University of Geneva; DNA from the same individual was used to generate previous versions of the genome[1,2]. Whole blood was extracted and stored in EDTA tubes at $-80°$C. The sample was shipped to Cold Spring Harbor Labs for the extraction of high molecular weight DNA, continuous long read library construction, and sequencing on a PacBio Sequel II platform. All genome assembly steps were performed on Columbia University's High Performance Computing (HPC) Terremoto cluster.

##### 1.2.1 Whole genome PacBio sequencing and assembly

The raw PacBio subreads were converted from `.bam` format to `.fastq` using the `bam2fastx` tool available from the `pbbioconda` package. We generated an initial PacBio-only assembly using `canu`[3] with the following command:

```
canu -p CS -pacbio-raw CS_genome.fastq.gz \\
    --gridOptions="-A palab" --gridOptionsovs="--mem-per-cpu=64G" \\
    --gridOptionsCNS="--mem-per-cpu=64G" genomeSize=1.7g
```

The initial PacBio `canu` assembly generated 23,086 contigs with an N50 of 325 Kb and a total sequence length of 2.72 Gb. We hypothesized that this increase in assembled sequence length relative to the Ullate-Agote (2020) genome was the result of unresolved haplotigs in the `canu` assembly. We therefore used `Purge Haplotigs`[4] to identify and realign allelic contigs. To prepare input data for the pipeline, we first used `minimap2`[5] to align trimmed PacBio subreads (output from an intermediate step in `canu`) to the initial `canu` assembly.

```
minimap2 -t 4 -ax map-pb ${REF} ${READS} --secondary=no > cs_aln.sam
```

After sorting and indexing the resulting alignment, we ran the three steps of the `Purge Haplotigs` pipeline:

```
purge_haplotigs hist -b ${INPUT} -g ${REF} -t 4

purge_haplotigs cov -i ${INPUT} -l 10 -m 55 -h 180 -o coverage_stats.csv \\
    -j 80 -s 80

purge_haplotigs purge -g ${REF} -c ${INPUT} -t 4
```

After running Purge Haplotigs, the total size of the assembly was substantially reduced, to 1.67 Gb, and the N50 increased to 958 Kb, indicating that there were indeed a large number of unresolved haplotigs in the initial canu assembly. The total number of contigs decreased to 3,656, of which the largest was 14.60 Mb.

Next, we performed a series of scaffolding, gap closing, and polishing steps. To scaffold the assembled contigs, we used LRScaf[6]. We prepared the input data by mapping trimmed and corrected PacBio subreads output from canu to the curated genome assembly from Purge Haplotigs. We then ran LRScaf on the resulting alignment:

```
INPUT=CS.correctedReads.fasta.gz
REF=curated.fasta

minimap2 -t 8 ${REF} ${INPUT} > intrm.corrected.aln.mm

java -jar LRScaf-1.1.11.jar -c curated.fasta -a intrm.corrected.aln.mm -t 4 \\
    -o lrscaf_corr_output
```

Once the scaffolding was completed, this intermediate assembly had an N50 of 1.021 Mb and a total sequence length of 1.74 Gb, consisting of 3,139 scaffolds. To fill in gaps in the assembly, we applied TGSGapcloser[7], using the corrected PacBio reads from canu:

```
$REF = scaffolds.fasta
$READS = CS.correctedReads.fasta.gz

tgsgapcloser --thread 8 --scaff ${REF} --ne --reads ${READS} --tgstype pb \\
    --output cs_gapfilled
```

The resulting assembly had an N50 of 1.048 Mb and a total length of 1.74 Gb, including 3,191 scaffolds. As a final step for the PacBio assembly, we performed polishing and error correction using high coverage Illumina paired end reads generated from the original Ullate-Agote (2020) assembly (SRA Accession Number: SRR9596760).

```
${REF} = cs_gapfilled.fasta
polca.sh -a ${REF} \\
    -r 'SRR9596760_1.fastq.trim.c1.fq.gz SRR9596760_2.fastq.trim.c2.fq.gz' -t 12
```

##### 1.2.2 Merging assemblies

To construct the final genome assembly, we used the tool quickmerge[8] in order to perform an all-vs-all alignment of the polished PacBio assembly and the Ullate-Agote (2020) assembly. We used the PacBio assembly as the "hybrid" input and the original assembly as the "self" input:

```
/quickmerge/merge_wrapper.py ${SELF_INPUT} ${HYBRID_INPUT} -l 100 \\
    --prefix ${PREFIX}
```

As a final step, we performed one more round of gap-filling using the same parameters for TGSGapcloser and the corrected PacBio reads.

Lastly, we used the Bionano optical maps from Ullate-Agote (2020) for a final round of scaffolding and to correct any misassemblies. To construct Bionano superscaffolds, we used the stitch.pl workflow available from the Irys Scaffolding pipeline (https://github.com/i5K-KINBRE-script-share/Irys-scaffolding)[9]. First, an in silico map was generated for the assembly with cut sites for the enzyme *BspQI* with the fa2cmap_multi.pl script. The in silico map was then used as input for the sewing_machine.pl script along with the merged assembly:

```
fa2cmap_multi.pl -v -i ${REF} -e BspQI

stitch/sewing_machine.pl -o sewing-machine-out -g CS.cmap -p ${REF} \\
    -r CS.BspQI.cmap
```

The final statistics of the merged and Bionano optically mapped assembly were:

```
Main genome scaffold total:          466
Main genome contig total:            4019
Main genome scaffold sequence total: 1708.106 MB
Main genome contig sequence total:   1649.955 MB   3.404% gap
Main genome scaffold N/L50:          10/63.229 MB
Main genome contig N/L50:            142/3.031 MB
Main genome scaffold N/L90:          54/3.485 MB
Main genome contig N/L90:            828/332.618 KB
Max scaffold length:                 141.592 MB
Max contig length:                   17.177 MB
Number of scaffolds > 50 KB:         360
% main genome in scaffolds > 50 KB:  99.78%
```

##### 1.3 Repeat annotation

Masking and annotating repeats was done following an approach similar to one found in Card, et al. 2019[10]. In short, we generated a de novo repeat library using RepeatModeler[11], curated the results to identify elements classified as unknown, and masked the genome using RepeatMasker[12] The Dfam TETools[13] singularity container v.1.5 was used to build a database from the genome assembly and to run RepeatModeler, including the long terminal repeat (LTR) discovery pipeline.

```
singularity exec docker://dfam/tetools:latest BuildDatabase \\
    -name Pantherophis_guttatus Pantherophis_guttatus_genome.fa

singularity exec docker://dfam/tetools:latest RepeatModeler \\
    -database Pantherophis_guttatus -pa 16 -LTRStruct
```

We confirmed that no protein-coding sequences were erroneously identified as repetitive elements by conducting a blastx[14] search against the nr protein database from the *Serpentes* taxon. Reassuringly, we found no hits.

```
blastx -db blastdb/nr -query repeat_modeler_families.fa \\
    -out protein_check.out -taxids 8570 -word_size 6
```

Many of the elements marked as unknown by RepeatClassifier were similar to elements found in an existing library of repetitive elements identified in 12 snake species[10]. We conducted a blastn search of our unknown de novo elements against the existing snake library and classified unknown elements as a given repetitive element family if over 50% of Blast hits for the unknown element were from one repeat family. In the absence of a majority consensus among the Blast hits, the de novo family remained classified as unknown. By this approach, we were able to assign 78.60% of the unknown elements to a family. We then used RepeatMasker to identify coordinates of repetitive elements and mask them using an iterative approach that prioritizes curated and classified elements. RepeatMasker was run using a BovB-CR1 library provided by Card *et. al.* (2019), the Dfam tetrapoda library[15], the classified elements from our RepeatModeler results, and finally the remaining unclassified elements. These steps masked 6.88%, 2.39%, 39.08%, and 2.96% of the genome, respectively.

```
singularity exec docker://dfam/tetools:latest RepeatMasker -pa 16 -gff \\
    -lib [Repeat_library.fa] Pantherophis_guttatus_genome.fa
```

The full pipeline identifies 48.90% of the genome as comprised of repetitive elements; 44.74% are transposable elements, the majority of which are retroelements (Table S4).

#### 1.4 Transcriptome

To annotate our genome assembly, we used a combination of PacBio Iso-seq reads that we collected from testes mRNA and publicly available Illumina RNA-seq data from a variety of tissues, as detailed in Table S6.

##### 1.4.1 PacBio RNA-seq

To accurately identify transcription start sites (TSS), especially for genes expressed in corn snake testes, we collected PacBio RNA-seq from two corn snake testis samples, provided to us from a colony of corn snakes maintained by one of us (A. T.) at the University of Geneva. The testes were dissected from two adult male corn snakes and stored in RNAlater (Thermo Scientific) at −80°C. RNA was extracted using TRIzol extraction and cleaned up using the RNAeasy Plus Mini kit (Qiagen). Roughly 30mg of tissue was placed in TRIzol and homogenized using a tissue lyser. The addition of chloroform followed by centrifugation was used to separate the phases, and the aqueous phase was saved. An equal volume of 70% ethanol was added before the solution was transferred to an RNeasy spin column and sample cleanup continued from step 7 of the kit protocol. The DNA from the eluted sample was removed using a TURBO DNA-free Kit (Invitrogen) following the manufacturer's protocol, the total RNA was quantified using a Qubit fluorometer (Thermo Scientific), and the sample as sent to Novogene, where PacBio Iso-Seq SMRTbell libraries were generated and sequenced on a Sequel II system.

For each sample, the .bam files containing subreads were first merged into a single file. Circular consensus sequence calling was then performed on the merged .bam files using PacBio ccs software under default parameters. Primer removal and barcode identification was performed with PacBio lima software, using the --isoseq option and --peek-guess to remove potential spurious false positive signals as recommended. Finally, PacBio isoseq software was used to refine, cluster, and polish the reads.

```
ccs ${INFILE} ${PREFIX}.ccs.bam --log-level INFO \\
    --report-json ${PREFIX}.report.json \\
    --hifi-summary-json ${PREFIX}.hifi_summary.json \\
    --log-file ${PREFIX}.ccs.log --report-file ${PREFIX}.report.txt \\
    --metrics-json ${PREFIX}.zmw_metrics.json.gz

lima --isoseq --dump-clips ${PREFIX}.ccs.bam primers.fasta ${PREFIX}.fl.bam \\
    --peek-guess --log-file lima.log

isoseq3 refine --require-polya ${PREFIX}.fl.NEB_5p --NEB_Clontech_3p.bam \\
    primers.fasta ${PREFIX}.flnc.bam
```

```
bamtools convert -format fasta -in ${PREFIX}.flnc.bam > \\
    ${PREFIX}.flnc.fasta

isoseq3 cluster ${PREFIX}.flnc.bam ${PREFIX}.polished.bam --verbose --use-qvs
```

The resulting high-quality (HQ), polished `.hq.bam` files were then merged into a single file, and aligned to the genome assembly using `minimap2` software. Identical isoforms were removed and redundant transcript models were collapsed using TAMA.

```
minimap2 -t 8 -R "@RG\tID:Sample\tSM:hs\tLB:ga\tPL:PacBio" --MD \\
    -ax splice:hq -uf --secondary=no ${REF} ${INFILE}> ${PREFIX}.aligned.sam

samtools sort -@ 8 -O BAM ${PREFIX}.aligned.sam -o ${PREFIX}.aligned.sort.bam

samtools index ${PREFIX}.aligned.sort.bam

python tama_collapse.py -s ${PREFIX}.aligned.sort.bam -f ${REF}
```

###### 1.4.2 Short-read transcriptome assembly

We also used publicly available short read RNA-seq data sets from several corn snake tissues in our annotation pipeline; see Table S6 for the NCBI accession information.

Using these data, we generated a de novo transcriptome assembly using `Trinity`[16]. As the data contained both paired and unpaired reads, the –left and –right flags were used for paired reads and the –single flag was used for unpaired reads.

```
Trinity --full_cleanup --seqType fq --left {*R1.fq.gz} --right {*R2.fq.gz} \\
    --single {*unpaired.fq.gz} --output {Output_Folder}
```

We aligned the de novo transcriptome to the updated reference genome using `GMap`[17] using default settings except that we did not set a limit on reported paths (–npaths=0), we removed failed alignments (–nofails), and we pruned poor and repetitive alignments (-p 3).

```
gmap_build -D {Database_Directory} -d {Genome_Name} {REFERENCE}

gmap -D {Database_Directory/Genome_Name} -d {Genome_Name} --npaths=0 \\
    --quality-protocol=illumina --format=gff3_gene --nofails -p 3 \\
    {Trinity_output.fasta} > {Trinity.gff3}
```

###### 1.4.3 Combining transcriptome data

We combined gene model predictions and transcript assemblies from both the short-read and PacBio long-read data in an iterative process using `PASA`[18] and `EvidenceModeler`[19] software to generate a final genome annotation. First, gene model prediction was performed on the short read transcriptome assembly alone with the `Braker`[20] pipeline. Full-length long read IsoSeq transcripts were mapped to the genome assembly with `PASA` (v.2.2) annotation software, using both `blat`[21] and `GMap` aligners, in order to generate an initial long-read only trancript database. We then integrated the short-read gene model predictions by loading the transcript annotations into the long-read `PASA` database to generate a first-round merged `GFF` file.

To update and refine the predicted gene models, we incorporated additional coding information from other sources with `EvidenceModeler` after producing an initial merged annotation. Protein sequences from the garter snake genome assembly (cite) were aligned to the updated reference genome with `exonerate`. We predicted open reading frames on the polished IsoSeq reads using `transdecoder` and additionally included the predicted transcripts from the initial merged `PASA` assembly. Finally, we included transcript sequences from the Ullate-Agote (2020) genome assembly. For each data source, we applied the following weights before running `EvidenceModeler`.

```
PROTEIN Exonerate_predictions 5
TRANSCRIPT pasa 8
ABINITIO_PREDICTION Braker.hmm.gff 10
TRANSCRIPT Ullate-Agote_transcripts 10
TRANSCRIPT IsoSeq 12
OTHER_PREDICTION transdecoder 1
```

The merged output from `EvidenceModeler` was then imported back in the `PASA` transcript database. Following the authors' recommendations (https://github.com/PASApipeline/PASApipeline/wiki/PASA_genome_annotation), two subsequent iterations of gene model comparison and updating were performed before the final `GFF` file was generated. The final annotation contains 18,822 protein-coding genes.

```
#Short read Braker run
braker2-2.1.6-hdfd78af_5/bin/braker.pl  --species=Cornsnake \\
    --genome=CS_superscaffold.renamed.fasta.masked.fasta --bam=CS.rna_seq.bam \\
    --softmasking --cores=24 --useexisting

#First long read-only PASA run
module load singularity
singularity exec ${PWD}/pasapipeline_latest.sif /bin/bash & export LANG=C & \\
    /usr/local/src/PASApipeline/Launch_PASA_pipeline.pl \\
    -c alignAssembly.config -C -R --transcribed_is_aligned_orient
    -g CS_superscaffold.renamed.fasta.masked.fasta --ALIGNERS blat,gmap \\
    -t CS.isoseq_collapsed.renamed.fasta --TDN isoseq.cleaned.accs \\

#Load short-read data in long-read database
/usr/local/src/PASApipeline/scripts/Load_Current_Gene_Annotations.dbi \\
    -c alignAssembly.config \\
    -g CS_superscaffold.renamed.fasta.masked.fasta -P braker.hmm.gff"

#Transdecoder
TransDecoder.LongOrfs -t CS.isoseq_collapsed.fasta

TransDecoder.Predict -t CS.isoseq_collapsed.fasta

#Exonerate
exonerate-2.4.0-h7c8e0dd_4/bin/exonerate -c 1 --model p2g --showvulgar no \\
    --showalignment no --showquerygff no --showtargetgff yes --percent 80 \\
    --ryo "AveragePercentIdentity: %pi\n" ${INFILE} ${REF} > ${INFILE}.gff

#EvidenceModeler
EVidenceModeler-1.1.1/EvmUtils/partition_EVM_inputs.pl \\
    --genome CS_superscaffold.renamed.fasta.masked.fasta \\
    --gene_predictions gene_predictions.gff \\
    --protein_alignments ThaEle_exonerate.renamed.gff \\
    --transcript_alignments all.transcript_alignments.gff3 \\
```

```
    --segmentSize 100000 --overlapSize 10000 \\
    --partition_listing partitions_list.out

EVidenceModeler-1.1.1/EvmUtils/write_EVM_commands.pl \\
    --genome CS_superscaffold.renamed.fasta.masked.fasta \\
    --weights `pwd`/weights.txt --gene_predictions gene_predictions.gff \\
    --protein_alignments ThaEle_exonerate.renamed.gff
    --transcript_alignments all.transcript_alignments.gff3 \\
    --output_file_name evm.out --partitions partitions_list.out > \\
    commands.list

EVidenceModeler-1.1.1/EvmUtils/execute_EVM_commands.pl commands.list \\
    | tee run.log

EVidenceModeler-1.1.1/EvmUtils/recombine_EVM_partial_outputs.pl \\
    --partitions partitions_list.out --output_file_name evm.out

EVidenceModeler-1.1.1/EvmUtils/convert_EVM_outputs_to_GFF3.pl \\
    --partitions partitions_list.out --output evm.out  \\
    --genome CS_superscaffold.renamed.fasta.masked.fasta
```

#### 1.5 CpG islands

##### 1.5.1 DNA methylation

Testes from two young adult *P. guttatus* samples from the corn snake colony at the University of Geneva (A.T.) were dissected, immediately flash-frozen in liquid nitrogen, and stored at -80°C. They were shipped on dry ice to Novogene, where the DNA was sheared using a Covaris S220 and used to prepare a methylation sequencing library. The library underwent Bisulfite Treatment using a EZ DNA Methylation Gold Kit (Zymo Research) and was then sequenced on a NovaSeq 6000 using PE150 sequencing (Illumina).

We made methylation calls for both samples in a single step, using the `methylseq` pipeline (https://zenodo.org/record/2555454) using `Nextflow` (https://nf-co.re/methylseq):

```
nextflow run methylseq -profile singularity --input ${FASTQ} --fasta ${REF}
```

We used the `coverage2cytosine` script in `Bismark`[22] to combine the methylation calls for the C and G in a CpG site into a single call:

```
coverage2cytosine -genome_folder {REF_FOLDER} --merge_CpG \\
    -o ${CPG_MERGED_OUT} ${METHYLSEQ_OUT} --gzip --zero_based
```

##### 1.5.2 CpG island identification

CpG islands are regions in the genome that contain a large number of hypomethylated CpG dinucleotide repeats and often serve as sites of transcription initiation[23]. In order to identify CpG islands in the corn snake genome, we initially used the `cpgplot`[24] tool in `EMBOSS:6.6.0.0` to detect regions where, over an average of ten windows of 100 bases and a minimum of 250 consecutive bases, the GC content is more than 50% and the calculated observed-to-expected ratio is > 0.6:

```
cpgplot -sequence ${REF} -outfile ${CPGPLOT_OUT} -window 50 -minlen 250 \\
    -minoe 0.6 -minpc 50 -graph png -outfeat ${CPGI.GFF} -plot No
```

To expand on the set identified by `cpgplot`, we combined this information with DNA methylation data from bisulfate sequencing (see 1.5.1). To this end, we defined hypomethylated CpG sites as positions where <50% of the reads support a methylated cytosine (note that the distribution of the fraction of methylated reads is bimodal, with peaks around ∼0% and ∼100%; FigureS27). We then implemented a hidden Markov model with six emissions, corresponding to the four 'standard' nucleotides, as well as a hypomethylated C and a hypomethylated G –cytosine in the complementary strand– and eight hidden states (corresponding to an 'island' and 'non-island' state for each 'standard' nucleotide). Since we do not have a gold standard set of CpG islands in cornsnakes, to set the transition and emission probabilities, we based ourselves on the CpG islands identified by `cpgplot`. We then used the function `MultinomialHMM` in the Python package `hmmlearn` to identify positions in the corn snake genome supporting the 'island' states. By this approach, the CpG islands identified contain the vast majority of those identified by `cpgplot` plus some islands that were presumably too short or missed for other reasons. In total, by this procedure we called 42,582 CpG islands, on par with numbers for other vertebrate species[25,26].

#### 1.6 Chromosome identification

##### 1.6.1 Sex chromosomes

We identified scaffolds in the corn snake assembly that likely belong to the sex chromosomes by analyzing depth of coverage patterns in females versus males. In corn snakes, females are the heterogametic sex (i.e., they carry a single Z chromosome), so in scaffolds belonging to the Z, the depth of coverage should be two times higher in males than in females on average. We obtained the depth of coverage for each scaffold in each parent in the corn snakes pedigrees, for which we know their sex:

```
samtools coverage -q20 -Q20 ${INDV.BAM} -r ${SCAFFOLD}
```

Consistent with the expectation, we found a few scaffolds deviating from the rest and showing patterns in depth of coverage that strongly suggested they belong to the sex chromosomes (FigureS24). We classified scaffolds as non-autosomal if the female-to-male depth of coverage ratio was ≤0.7 (putatively belonging to the Z) or ≥1.3 (putatively belonging to the W). This procedure identified 17 scaffolds (TableS5 and FigureS24), and for the candidates in the Z chromosome, their length added up to ∼135Mb, which is similar to the Z chromosome length in the high-quality assembly of *Thamnophis elegans* (∼145Mb, `https://www.ncbi.nlm.nih.gov/assembly/GCF_009769535.1`). Putative sex chromosomes were excluded from downstream analyses.

##### 1.6.2 Macro- and microchromosomes

During preparation of our manuscript, we became aware of another *P. guttatus* assembly update, made publicly available at `https://www.dnazoo.org/assemblies/Pantherophis_guttatus`[27]. This assembly is also a hybrid assembly, generated using the Ullate-Agote (2020) assembly and additional Hi-C data. While the DNA Zoo assembly has a low contig N50 (38 kb) compared to our genome assembly (3031 kb), it assembles 92% of the genome into 18 chromosomes that match the *P. guttatus* karyotype: 8 macrochromosomes and 10 microchromosomes. We used the DNAZoo genome assembly to assign our scaffolds to macro- and microchromosomes using `MUMmer4`; the alignment was run using the `nucmer` function with anchors that are unique in both genomes (`--mum`), filtered using `delta-filter`, and analyzed using `show-coords`.

```
nucmer --mum --delta {DNAZoo_Vs_NewGenome.delta} \\
    {DNAZoo_Assembly.fa} {Updated_Hybrid_Assembly.fa}

delta-filter -g {DNAZoo_Vs_NewGenome.delta} -l 100000 > \\
    {DNAZoo_Vs_NewGenome_filtered.delta}
```

```
show-coords {DNAZoo_Vs_NewGenome_filtered.delta} > \\
    {DNAZoo_Vs_NewGenome_filtered.txt}
```

In order to assign scaffolds from our genome assembly as macro- or microchromosomes, we only considered scaffolds where at least 50% of the bases map to the DNAZoo genome. Scaffolds where more than 75% of the aligned bases fall on a macrochromosome are assigned as a macrochromosome, and the same for microchromosomes. This assigns 95% of autosomal scaffolds larger than 100kb to either macro- or microchromosomes (85% and 11%, respectively). The remaining 5% of the scaffolds were excluded from analyses of macro- vs. microchromosomes.

### 2 Whole genome polymorphism data collection

#### 2.1 Whole genome sample resequencing

*P. guttatus* samples for whole genome sequencing came from two different sources: 25 wild-caught corn snakes kindly provided to us by Dr. Edward Myers at the the American Museum of Natural History, and 19 samples, including unrelated and pedigree samples, from the corn snake colony at maintained by A.T. at the University of Geneva.

DNA was extracted and sequenced for six colony samples (identified in Table S1) as in Ullate et al., 2020[1].

For the remaining 37 samples, we extracted genomic DNA from the provided tissue samples using the Qiagen DNeasy Blood & Tissue kit, using the protocol for "Purification of Total DNA from Animal Tissues" and quantified the DNA concentration using a Qubit fluorometer (Thermo Scientific, Wilmington, DE). We prepared genomic libraries for high coverage sequencing following Quail *et al.* (2009)[28]. Briefly, we sheared approximately 500 ng of DNA to ∼400 bp with a QSonica sonicator (QSonica Sonicators, Newton, Connecticut). We mixed DNA with dNTPs and T4 DNA polymerase, Klenow DNA polymerase and T4 PNK and incubated at room temperature for 30 minutes (NEB, Ipswich, MA) to repair the sheared ends, and then purified the reaction with the Qiagen QIAquick PCR purification kit (Qiagen, Valencia, CA). A-tails were added by mixing the purified end-repaired DNA with dATPs and Klenow exonuclease and incubating at 37° C for 30 minutes (NEB, Ipswich, MA) and then purified the reaction with the Qiagen QIAquick PCR purification kit (Qiagen, Valencia, CA). We performed adapter ligation reaction followed by purification with the Qiagen QIAquick PCR purification kit (Qiagen, Valencia, CA), amplified the adapter ligated DNA using singly indexed primers in individual Phusion PCR reactions for 12 cycles and then purified the libraries using 18% SPRI beads.

We quantified libraries with a Qubit fluorometer (Thermo Scientific, Wilmington, DE) and assessed library size distribution and quality using Agilent 4200 Tapestation (Agilent, Santa Clara, CA). We sent libraries for sequencing on an Illumina Novoseq 6000 with 2 x 150 cycles at the New York Genome Center, New York, NY or an Illumina HiSeq 4000 with 2 x 150 cycles at Admera Health Services, South Plainfield, NJ (see Table S1).

#### 2.2 Read alignment and processing

Reads were trimmed using `trimmomatic`[29] to remove adaptors and low quality bases. We aligned the reads to the genome using `bwa mem`[30] and sorted the resulting alignments using `samtools`[31]. Read groups identifying the library and sequencing lane were added using `picard AddOrReplaceReadGroups`[32] and reads from the same library were then merged using `samtools merge`. We identified duplicate reads from each library separately using `picard MarkDuplicates`, specifying an optical distance of 2500 and 12000 for libraries sequenced on Hiseq machines and Novoseq machines, respectively. All libraries from each sample were merged using `samtools merge` and sample genome-wide coverage was assessed using `picard CollectWgsMetrics`.

```
java -jar {PATH_TO_TRIMMOMATIC.jar} PE -threads 6 \\
```

```
    {SAMPLE_LIBRARY_LANE_R1.fq.gz} {SAMPLE_LIBRARY_LANE_R2.fq.gz} -baseout \\
    {SAMPLE_LIBRARY_LANE_TRIMMED.fq.gz} MINLEN:40 \\
    ILLUMINACLIP:{PATH_TO_ADAPTERS.fa}:2:30:10:8:true SLIDINGWINDOW:4:15

bwa mem -t 6 {REFERENCE} {SAMPLE_LIBRARY_LANE_TRIMMED_1P.fq.gz} \\
    {SAMPLE_LIBRARY_LANE_TRIMMED_2P.fq.gz} | samtools view -b -h -@ 6 \\
    | samtools sort -@ 6 -O BAM -o {SAMPLE_LIBRARY_LANE.bam}

java -Xmx4G -jar {PATH_TO_PICARD.jar} AddOrReplaceReadGroups \\
    I={SAMPLE_LIBRARY_LANE.bam} O={SAMPLE_LIBRARY_LANE_RG.bam} \\
    RGID={SAMPLE_LIBRARY_LANE} RGLB={SAMPLE_LIBRARY} \\
    RGPU={SAMPLE_LIBRARY_LANE} RGPL=illumina RGSM={SAMPLE}

samtools merge -@ 4 {SAMPLE_LIBRARY.bam} {SAMPLE_LIBRARY_*_RG.bam}

java -Xmx6g -jar {PATH_TO_PICARD.jar} MarkDuplicates ASO=coordinate \\
    OPTICAL_DUPLICATE_PIXEL_DISTANCE={DISTANCE} I={SAMPLE_LIBRARY.bam} \\
    O={SAMPLE_LIBRARY_MD.bam} M={SAMPLE_LIBRARY_MD.txt} USE_JDK_DEFLATER=true

samtools merge {SAMPLE.bam} {SAMPLE_*_MD.bam}

java -Xmx2G -jar {PATH_TO_PICARD.jar} CollectWgsMetrics I={SAMPLE.bam} \\
    O={SAMPLE_WGSMETRICS.txt} R={REFERENCE}
```

#### 2.3 Removal of contaminated samples

After preparing the `bam` files, we generated a preliminary set of variants using `GATK`[33] for variant calling and following GATK best practices for filtering. We used `vcftools relatedness2`[34,35] to assess relatedness between samples in this preliminary variant call set and noticed that two of the high coverage samples showed inflated levels of relatedness with many other samples (Figure S25).

```
vcftools --gzvcf {HC_HWE_Unrelated.recode.vcf.gz} --relatedness2 \\
    --out {Relatedness_File}
```

By looking at the fraction of sequencing reads supporting the alternative allele at heterozygous positions (as per the AD field output by GATK's HaplotypeCaller), we found that these same libraries showed abnormal patterns of allelic balance (Figure S25)). We used `GATK GetPileupSummaries` and `CalculateContamination` to calculate the amount of cross-sample contamination present in each library-specific `.bam` file.

```
gatk GetPileupSummaries -I {SAMPLE_LIBRARY_MD.bam} -V {Filtered_SNPs.vcf.gz} \\
    -L {Filtered_SNPs.vcf.gz} -O {SAMPLE_LIBRARY.table}

gatk CalculateContamination -I {SAMPLE_LIBRARY.table} -O {SAMPLE_LIBRARY.txt}
```

The same two libraries that showed inflated relatedness values and abnormal allelic balance patterns were also outliers for the estimated fraction of contaminated reads; 20-30% of these libraries were estimated to be the result of cross-sample contamination. We removed these libraries from all subsequent analyses. One additional library contained approximately 10% contaminated reads; out of caution, we also removed this library. We recalculated the coverage with the remaining libraries (Table S1) and proceeded with variant calling (section 2.4) using only the other libraries and re-calculated relatedness to confirm that removing the contaminated libraries greatly reduced the estimated relatedness for the affected samples (Figure S25)).

#### 2.4 Variant calling

Variants were called using GATK; specifically, HaplotypeCaller was used to generate per sample GVCF files for each autosomal scaffold larger than 100 kb. The GVCFs were validated using ValidateVariants and no problems were detected. For each scaffold, we then combined GVCFs from each sample using GenomicsDBIImport and genotyped using GenotypeVCFs, including non-variant sites and using a heterozygosity of .003 per base-pair, which we estimated based on a preliminary variant calling run.

```
gatk HaplotypeCaller -L {SCAFFOLD} -R {REFERENCE} -I {SAMPLE.bam} \\
    -O {SAMPLE_SCAFFOLD.gvcf} -ERC GVCF

gatk ValidateVariants -gvcf -L {SCAFFOLD} -R {REFERENCE} \\
    -V {SAMPLE_SCAFFOLD.gvcf}

gatk GenomicsDBImport --genomicsdb-workspace-path {SCAFFOLD_DATABASE} \\
    --L {SCAFFOLD} -V {Sample1_SCAFFOLD.gvcf.gz} \\
    -V {Sample2_SCAFFOLD.gvcf.gz}...

gatk GenotypeGVCFs --include-non-variant-sites -L {SCAFFOLD} -R {REFERENCE} \\
    -V gendb://{SCAFFOLD_DATABASE} -G StandardAnnotation \\
    --heterozygosity .003 -O {SCAFFOLD.vcf}
```

The resulting VCF files contain 53,261,726 putative SNPs called across 1.52 GB (89.2%) of the genome.

#### 2.5 Filtering

Variants were filtered following GATK best practices[36] with some additional filters chosen with our applications in mind.

##### 2.5.1 Mappability

To restrict our analyses to locations where short-read sequences can be confidently mapped, we built a mappability mask following the SNPable pipeline with a read length of 150 base-pairs (http://lh3lh3.users.sourceforge.net/snpable.shtml), complemented with the script makeMappabilityMask.py in https://github.com/stschiff/msmc-tools:

```
splitfa {Fasta} 150 | split -l 20000000 --filter='gzip > \$FILE.gz' \\
    - kmers/{Chunk};

bwa aln -R 1000000 -O 3 -E 3 {fasta} kmers/{Chunk} > bwa/{Chunk}.sai;

bwa samse {Fasta} bwa/{Chunk}.sai kmers/{Chunk} | gzip > bwa/{Chunk}.sam.gz;

gzip -dc bwa/*.sam.gz | perl gen_raw_mask.pl > rawMask_150.fa;

gen_mask -l 150 -r 0.5 rawMask_150.fa > mask_150.fa;

python makeMappabilityMask.py mask_150.fa > {Mappability_mask.bed}
```

The mappability mask ultimately excludes 14.3% of the assembled genome.

##### 2.5.2 Variant filtering

To identify the set of sites where we would likely detect a SNP if present, we used a combination of `bcftools filter`[37] and `bcftools view` to remove all sites within 5 bp of an indel or genotypes with very high or low depth values and keep only invariant sites and SNPs with less than 25% missingness and a maximum of two alleles across individuals. The depth filter set all genotypes with depth lower than 10x or depth higher or lower than two standard deviations from the mean genome-wide coverage of the sample to "missing" (DEPTH_FILTER in the command below); the mean depth and standard deviation were determined by `picard CollectWgsMetrics`. We then used `tabix`[38] to remove repetitive regions annotated in section 1.3 and regions in the mappability mask (see section 2.5.1). The 680,899,915 "callable" sites (40.0% of the genome) remaining after these filtering steps were used to calculate the denominator for diversity statistics.

```
bcftools filter -g 5 --set-GTs . -e '{DEPTH_FILTER} || TYPE~"indel"' -O u \\
    {SCAFFOLD.vcf.gz} | bcftools view -O z -i 'F_MISSING<.25 & \\
    (TYPE="snp" || TYPE="ref")' -V indels,mnps --max-alleles 2 \\
    -o {SCAFFOLD_FILTER1.vcf.gz}

cat {REPEATS.bed} {LOW_MAPPABILITY.bed} | bedtools sort -g {REFERENCE.genome} \\
    | bedtools merge > {MASK_REPEATS_MAPPABILITY.bed}

bedtools complement -i {MASK_REPEATS_MAPPABILITY.bed} \\
    -g {REFERENCE.genome} > {COMPLEMENT_REPEATS_MAPPABILITY.bed}

tabix -h -R {COMPLEMENT_REPEATS_MAPPABILITY.bed} \\
    {SCAFFOLD_FILTER1.vcf.gz} > {SCAFFOLD_FILTERED_ALLSITES.vcf.gz}
```

We then used `GATK SelectVariants` to remove all invariant sites and sites with QD <10.0, MQ <40.0, FS >10.0, SOR >4.0, MQRankSum <-12.5, and ReadPosRankSum <-8.0, and `GATK VariantFiltration` to set sites with GQ <20 as "missing". Genotypes were set to missing if the p-value from a Binomial test on allelic balance was less than $1 \times 10^{-7}$ (as determined by a custom script). We excluded sites with greater than three genotypes that met this criterion and `bcftools view` was used to again remove all invariant sites or sites with missingness >25%. Finally, we used `vcftools` to identify SNPs out of Hardy-Weinbreg equilibrium in unrelated samples; sites with a p-value less than .001 were removed.

```
gatk IndexFeatureFile -F {SCAFFOLD_FILTERED_ALLSITES.vcf.gz}

gatk SelectVariants --selectExpressions 'QD < 10.0 || MQ < 40.0 || FS > 10.0 \\
    || SOR > 4.0 || MQRankSum < -12.5 || ReadPosRankSum < -8.0' \\
    --invertSelect --exclude-non-variants true -R {REFERENCE} \\
    -V {SCAFFOLD_FILTERED_ALLSITES.vcf.gz} -O {SCAFFOLD_SNPS_FILTER1.vcf.gz}

gatk VariantFiltration --genotype-filter-name 'VF' \\
    --set-filtered-genotype-to-no-call --genotype-filter-expression "GQ < 20" \\
    -V {SCAFFOLD_SNPS_FILTER1.vcf.gz} -O {SCAFFOLD_SNPS_FILTER2.vcf.gz}

ipython Remove_AllelicBalance.py {SCAFFOLD_SNPS_FILTER2.vcf.gz} \\
    {SCAFFOLD_SNPS_FILTER3.vcf}

bcftools view -O u -i 'F_MISSING<.25' -f .,PASS {SCAFFOLD_SNPS_FILTER3.vcf.gz} \\
    | bcftools view -c 1:minor -O z -o {SCAFFOLD_SNPS_FILTER4.vcf.gz}

bcftools view -s {Unrelated_Samples} -O v {SCAFFOLD_SNPS_FILTER4.vcf.gz} | \\
```

```
    vcftools --hardy --vcf - --out {SCAFFOLD_HWE_Pvals}

vcftools --recode --gzvcf {SCAFFOLD_SNPS_FILTER4.vcf.gz} --exclude-positions  \\
    {SCAFFOLD_HWE_LowPVals.bed} --out {SCAFFOLD_SNPS_FILTER5}
```

By analyzing the individuals in the two families, we determined the number of instances where a non-singleton autosomal variant violated the principles of Mendelian segregation (0.31% of SNPs in our sample). We masked out all of these SNPs, as well as any genomic regions containing three or more consecutive Mendelian errors (totaling ∼67 Mb, the regions were delimited by the outermost erroneous SNPs in any consecutive stretch) and scaffolds with an abnormally high proportion of Mendelian errors (i.e., ≥5% of their polymorphisms, which identified 14 scaffolds covering a total of ∼2.5 Mb, see S26). The 12,488,572 remaining SNPs are included in our "All Filtered SNPs" set and are used in subsequent analyses.

```
bedtools complement -i {MENDELIAN_ERROR_MASK.bed} -g {REFERENCE.genome} > \\
    {COMPLEMENT_MENDELIAN_ERRORS.bed}

tabix -h -R {COMPLEMENT_MENDELIAN_ERRORS.bed} {SCAFFOLD_SNPS_FILTER5.vcf.gz} > \\
    {SCAFFOLD_ALLFILTERED_SNPS.vcf.gz}
```

For many downstream analyses, we used the subset of variants from the 24 "unrelated" -i.e. not closely related- individuals sequenced to moderate to high (13-48X) coverage. To obtain this set of SNPs, we used `bcftools view` to select the samples and included only SNPs at which fewer than 25% of genotypes among these samples were missing: 11,544,003 SNPs.

```
bcftools view -s {Unrelated_Samples} -O v {SCAFFOLD_ALLFILTERED_SNPS.vcf.gz} | \\
    bcftools view -i 'F_MISSING<.25' -c 1:minor -O z -o \\
    {SCAFFOLD_FILTERED_HICOV_UNREL_SNPS.vcf.gz}
```

We refer to this final variant set as the "high coverage unrelated" one.

#### 2.6 Variant phasing

We phased the variants using both phase-informative reads and statistical phasing, incorporating the familial relationships within our two pedigrees. First, we used the `ExtractPIRs` module from `Shapeit2`[39] to identify phase-informative reads and `Plink`[40] to remove samples with <10x genome-wide sequencing coverage and converted the "All Filtered SNPs" `vcf` file to `bed/map` format. Phasing was then completed with `Shapeit2`, including both the -assemble and -duohmm flags to use the read-aware model and then combine the estimated haplotypes with the pedigree information. We set an estimated effective population size of Ne=250,000 based on our $\boldsymbol{\theta}$ estimate (see section 5.1.1 below), a population recombination rate $\boldsymbol{\rho}$ of 0.0023 per bp, the mean recombination rate from a preliminary run of `LDhelmet`[41], and a window length (-W) of 5 MB based on the suggested settings for `duohmm` when large amounts of IBD sharing are present, as in the pedigrees. Finally, we converted the resulting haplotypes back to vcf format using `shapeit convert`.

```
extractPIRs --bam {Bam_List.txt} --vcf {SCAFFOLD_ALLFILTERED_SNPS.vcf.gz} \\
    --out {SCAFFOLD_PIRs.txt}

plink2 --allow-extra-chr --remove {LowCoverage_Indv_List.txt} --update-parents \\
    {Pedigree_Info.txt} --vcf {SCAFFOLD_ALLFILTERED_SNPS.vcf.gz} --make-bed \\
    --out {SCAFFOLD_ALLFILTERED_SNPS_HICOV}
```

```
shapeit -assemble --input-bed {SCAFFOLD_ALLFILTERED_SNPS_HICOV.bed} --aligned \\
    --input-pir {SCAFFOLD_PIRs.txt} -O {SCAFFOLD_HAPS.txt} --effective-size \\
    250000 --rho 0.0023 --duohmm -W 5

shapeit -convert --aligned --input-haps {SCAFFOLD_HAPS.txt} --output-vcf \\
    {SCAFFOLD_PHASED.vcf}
```

### 3 PRDM9 binding sites

To identify putative PRDM9 binding sites in the *P. guttatus* genome, we followed a similar approach to Baker et al. 2017[42]. In short, we obtained sequences of the PRDM9 zinc fingers, used previously published[43] computational models to predict the position weight matrix for each allele, and then identified putative binding sites in the genome using `Fimo` from `Meme Suite`[44].

#### 3.1 Sequencing PRDM9 zinc finger alleles

The zinc finger array is extremely difficult to sequence, as it has a mini-satellite like structure of variable length[45,46]. It is all the more challenging in corn snakes, which appear to harbor an extremely high diversity of PRDM9 alleles, both in terms of the number of fingers and the identity of the residues in contact with DNA. We therefore used many strategies, including Sanger sequencing and long read sequencing. Table S2 contains a summary of the alleles that we identified.

##### 3.1.1 PRDM9 amplicon sequencing

We amplified the zinc finger array of PRDM9 from genomic DNA isolated from 22 corn snake individuals in section 2.1, using a variety of different PCR primers, as included in Table S7. Specifically, we used OneTaq Hot Start Quick-Load 2X Master Mix with Standard Buffer (NEB) and ran the PCR amplification as instructed by the protocol for this product with a 30s denaturation step, a 30s annealing step at 55-58°C, and a 3 min extension step for 30-35 cycles. To determine whether the PCR amplification was successful and look for individuals carrying two PRDM9 alleles of differing zinc finger lengths, we used gel electrophoresis.

We attempted to Sanger sequence any PCR reaction that yielded a single primary and relatively short band (indicating a PCR product less than 2 kb in length). We cleaned up these PCR products using a DNA Clean & Concentrator-5 kit (Zymo Research) and sent them for Sanger sequencing at Genewiz from Azenta Life Sciences. The primers used for sequencing are included in Table S7. We visually confirmed that the sequencing results represented a single PRDM9 zinc finger allele using the sequencing traces, and discarded any reads that had multiple sites with two prominent peaks, as these reads likely represent two distinct PRDM9 alleles. Next, we aligned forward and reverse Sanger sequencing reads from a given allele using `blastn`, keeping only convincing alignments where three or more zinc fingers overlapped with more than 97% identity. We only report alleles for which we obtained the complete zinc finger array, i.e., with reads containing conserved sequence upstream of the primary zinc finger array as well as the stop codon. In this way, we were able to identified four of the alleles carried by 11 individuals, but were unable to confidently sequence or assemble longer alleles carried by these and other individuals. We also were unable to assemble alleles from heterozygotes with zinc finger arrays of similar length.

##### 3.1.2 MinION Oxford Nanopore sequencing

To obtain the sequence of additional alleles, we turned to Oxford Nanopore sequencing. To this end, we prepared 21 libraries from 18 *P. guttatus* samples and three *Crotalus viridis* samples using the Ligation Sequencing Kit (SQK-LSK110) with the Native Barcoding Expansion 1-12 (EXP-NBD104) (Oxford Nanopore). We followed the community protocol "Ligation sequencing amplicons - native barcoding (SQK-LSK110 with EXP-NBD104 and EXP-NBD114)" `https://community.nanoporetech.com/docs/prepare/`

library_prep_protocols/native-barcoding-amplicons/v/nba_9093_v109_revn_12nov2019 (accessed July 8, 2022).

Briefly, we used PCR to amplify alleles from genomic DNA and cleaned up successful reactions as for Sanger sequencing. We assessed DNA concentration using a Qubit fluorometer (Thermo Scientific) and then end-prepped 150 fmol of each sample using the NEBNext Ultra II End repair/dA-tailing Module (NEB) and ligated the Native barcodes using Blut/TA Ligase Master Mix (NEB). We cleaned up the DNA bewteen each reaction using AMPure XP beads (Beckman Coulter) and, after again checking library concentration, pooled equimolar amounts of each library together. Finally, we ligated the adapters using the Quick T4 DNA Ligase in the NEBNext Quick Ligation Module (NEB) as directed.

We sequenced 12 libraries on a Flongle 9.4.1 flow cell and then 12 libraries on a MinION 9.4.1 flow cell (Oxford Nanopore); the second run included three repeated and nine additional samples. Sequencing was completed using the `MinKNOW` software, followed by `guppy` for base calling and demultiplexing using recommended settings.

```
guppy_basecaller -i {Run_Folder} -s {Run_Folder_fqs} --flowcell FLO-FLG001 \\
    --kit SQK-LSK110 --cpu_threads_per_caller 2 --num_callers 4 -r

guppy_barcoder --barcode_kits EXP-NBD104 -i  {Run_Folder/pass} -s {Run_Folder}
```

Reads from the three individuals that we sequenced twice were combined. We then plotted the read lengths for each individual. For 16 of the 18 samples, the distribution of the lengths of the reads had one or two peaks between 1500bp and 3000bp. To isolate the reads from these 16 samples, which were most likely to contain the complete zinc finger array, we selected reads within 42 bases (half of the length of a zinc finger) of the top two modes of the length distribution. For two of the samples (annotated in Table S2), we additionally selected a peak between 2 and 3 kb, which visually looked as though it might have represented a long allele, but did not contain a mode. We then ran `Canu`[3] to assemble the reads from each mode into contigs. This procedure assembled complete zinc finger arrays for two distinct alleles in three individuals; these individuals were among those with many reads falling into two distinct peaks of read length.

```
canu -p {Sample_ModeA} corMinCoverage=5 minOverlapLength=300 ovsThreads=1 \\
    useGrid=false -d {Sample_ModeA} genomeSize=3k maxInputCoverage=100 \\
    stopOnLowCoverage=5 minInputCoverage=5 -nanopore {Sample_ModeA_reads.fq}
```

For each of the 13 remaining samples, we identified "assembled" and "unassembled" contigs or corrected reads from the output of `canu` and used the `MAFFT`[47] web server (https://mafft.cbrc.jp/alignment/server) to generate multiple sequence alignments and phylogenetic trees with these data using automatic settings (except that we allowed the alignment to consider both forward and reverse sequence orientation). For three samples, the resulting trees were monophyletic with very few differences between sequences; either these samples are homozygous or we failed to amplify or sequence one of the alleles. The initial `canu` run assembled one complete zinc finger for each of these samples. The trees for the remaining 10 samples contained exactly two groups of sequences, representing two different PRDM9 alleles. We selected one sequence from each group and tried to isolate raw reads that had originated from the two alleles by aligning all of the Nanopore reads from the sample to these two sequences using `minimap2` and filtering the resulting `.bam` file to include only uniquely mapping, high quality alignments using `samtools`. Finally, we extracted the raw reads associated with the filtered alignments and re-ran `canu` as above. This procedure successfully identified two complete zinc finger arrays for nine of the samples.

```
minimap2 -ax map-ont {Sample_Sequences_from_tree.fa} {Sample_reads.fq} | \\
    samtools sort | samtools view -b -F 2304 -m 300 -o {Sample.bam}

samtools view {Sample.bam} {SequenceA_name} | cut -f1 > \\
```

```
    {Sample_SequenceA_readnames.txt}

cat {Sample_reads.fq} | grep --no-group-separator -A 3 \\
    -f {Sample_SequenceA_readnames.txt} > {Sample_SequenceA.fq}

canu -p {Sample_SequenceA} corMinCoverage=5 minOverlapLength=300 ovsThreads=1 \\
    useGrid=false -d {Sample_SequenceA} genomeSize=3k maxInputCoverage=100 \\
    stopOnLowCoverage=5 minInputCoverage=5 -nanopore {Sample_SequenceA.fq}
```

The final three samples, of which two did not have obvious peaks in the distribution of read lengths, were analyzed by hand by visually grouping similar reads. This process identified one additional *P. guttatus* allele (PRDM9-I) from one sample and two additional *C. viridis* alleles (CV-E and CV-F) from one sample (annotated in Table S2). We note that the inclusion of these alleles does not qualitatively change our results, and provide the raw reads for all samples for reproducibility.

Finally, we confirmed that there were no obvious, shared discrepancies between the raw reads and the final alleles by aligning the raw reads to the final alleles as above and visualizing the results using `IGV`. In total, we identify 17 alleles from 14 *P. guttatus* individuals and six alleles from three *C. viridis* individuals. Reassuringly, two alleles that were sequenced both using Sanger sequencing and Oxford Nanopore are identical. All samples and alleles are summarized in Table S2.

###### 3.1.3 Identifying PRDM9 from PacBio reads

For our genome assembly and annotation, we also collected multiple PacBio datasets, which contained additional PRDM9 alleles.

In order to extract the PRDM9 transcripts from the two PacBio Iso-Seq testes RNA samples described in section 1.4.1, we used `pbmm2` to align the the consensus (ccs) reads to the *P. guttatus* PRDM9 predicted transcript, excluding the zinc finger array. The reads associated with the resulting alignments contained three PRDM9 alleles, including one that was also identified in another individual using Sanger sequencing.

```
pbmm2 align {PRDM9_Nterm.fa} {Sample_ccs.bam} {Sample_PRDM9_ccs.bam} --sort
```

For the whole genome PacBio continuous long reads from a single sample described in section 1.2, we again identified reads containing PRDM9 by aligning the raw reads to the predicted PRDM9 N-terminal domains. We then followed the same protocol as with the Oxford Nanopore reads above and we were able to successfully assemble two complete PRDM9 alleles, including one also identified using Oxford Nanopore sequencing.

All of these alleles are summarized in Table S2.

##### 3.2 Predicting binding sites

To predict the binding sites of each corn snake PRDM9 allele, we used a computational model from Persikov *et. al.* (2014)[43], using as input the amino acid sequence of the zinc finger array for the "Predict PWMs" feature (`http://zf.princeton.edu/`). For a subset of alleles, the Sanger sequencing reads contain a zinc finger upstream of the primary zinc finger array, which is not involved in binding in mammals[48,49]; we excluded that finger in those cases. We generated the position weight matrix (PWM) for each allele using the Random Forest "RF regression on B1H" model option, downloaded the resulting PWMs, and used `Meme Suite`[50] to identify predicted binding sites for each allele in the *P. guttatus* genome. Specifically, we generated a second order background model for the reference genome using `fasta-get-markov` and ran `uniprobe2meme` to convert the PWM to `meme` format. We then identified predicted binding sites for PRDM9 allele in the corn snake genome using `FIMO`[44], with a p-value cutoff of $1 \times 10^{-5}$.

```
fasta-get-markov -m 2 {REFERENCE} {Ref_Background.txt}

uniprobe2meme -bg {Ref_Background.txt} -pseudo 1 {PWM.txt} > {PWM_meme.txt}

fimo --thresh .00001 -bfile {Ref_Background.txt} -oc {PWM_Folder} \\
    {PWM_meme.txt} {REFERENCE}
```

We used a custom python script to format the results as /texttt.bed files, kept the top 50,000 sites per allele based on the binding score, and merged overlapping sites within a given allele and among alleles using `bedtools`[51].

##### 3.3 Identifying zinc fingers shared among alleles

The population recombination rate estimated from LD data is a complicated average over the fine-scale genetic maps of all the PRDM9 alleles present in the ancestors of our sample. PRDM9 alleles at higher frequency in the population or that have persisted for longer will contribute more to the estimate. Therefore, zinc fingers that are shared among PRDM9 alleles may be more likely to show a signal of increased recombination as compared to the binding sites of any one PRDM9 allele or possibly to the union of all alleles. With this consideration in mind, we used a custom script to divide each allele's position weight matrix into sets of zinc fingers (ZF sequences) of length 5 to 15 and counted the number of observations of each ZF sequence. We then compared the ZF sequences to each other, and removed ZF sequences that were entirely contained within a larger ZF sequence and occurred the same number of times. For example, if three ZF sequences were: ABCDE identified six times, ABCDEF identified six times, and ABCDEFG identified five times, we would remove ABCDE. This procedure identified nine ZF sequences, ranging from five to 12 zinc fingers in length and found in at least five separate alleles, shown in Figure S10.

### 4 Population genetic analyses

##### 4.1 Characterization of population structure in the samples

Relationships among unrelated high coverage samples were assessed using principal component analysis (PCA). Filtered VCFs containing the "high coverage unrelated" set of variants from the largest 95 scaffolds were converted into `map/bed` format by `vcftools`, as `Plink` takes up to 95 scaffolds as input. We used `Plink` to prune SNPs based on pairwise linkage disequilibrium levels and also removed SNPs with a minor allele frequency $< 5\%$ and missingness $> 10\%$. These steps yielded 583,817 remaining SNPs, which we then input to the PCA using `plink --pca`.

```
vcftools --gzvcf {Filtered_HC_Unrelated.vcf.gz} --chrom-map {Chrommap.bed} \\
    --plink {out {Filtered_HC_Unrelated_plinkformat} --chr {Scaffold_1..95}

plink --allow-extra-chr --geno 0.10 --chr-set 95 no-xy no-mt \\
    --indep-pairwise 50 5 0.2 --maf 0.05 --out {SNPs_plinkformat_pruning} \\
    --file {Filtered_HC_Unrelated_plinkformat}

plink --chr-set 95 no-xy no-mt --allow-extra-chr --pca \\
    --file {SNPs_plinkformat} --extract {SNPs_plinkformat_pruning.prune.in} \\
    --out {SNPs_plinkformat_PCA}
```

We also calculated Weir and Cockerham's Fst[52] using the "high coverage unrelated" set of variants from the University of Geneva colony and the wild-caught samples in 10 kb overlapping windows with a 5 kb step, using `vcftools`.

```
vcftools --gzvcf {HC_HWE_Unrelated.recode.vcf.gz}  \\
    --weir-fst-pop {Colony_Samples.txt} --weir-fst-pop {Wild _Samples.txt} \\
    --fst-window-size 10000 --fst-window-step 5000 --out {FST.bed}
```

##### 4.2 Linkage disequilibrium levels

To better understand the distance over which linkage disequilibrium (LD) typically decays in the corn snake genome, we calculated pairwise allelic correlations, $r^2$. Using the same SNPs on the largest 95 scaffolds as in section 4.1, we used `plink` to randomly thin 100 fold the SNPs and calculated the mean $r^2$ between SNPs within up to a distance of 100 kb. We used a custom python script to calculate the distance between each pair of SNPs, group the distances between SNPs in 100 bp bins, and calculate the mean $r^2$ within each bin.

```
plink --chr-set 95 no-x no-y no-xy no-mt --file {SNPs_plinkformat} --thin .01 \\
    --r2 --out {SNPs_plinkformat} --ld-window 100000 --ld-window-kb 100 \\
    --ld-window-r2 0
```

##### 4.3 Heterozygosity

We calculated $\pi$ using the "high coverage unrelated" set of variants in overlapping 10 kb windows with a 5 kb step size, using `vcftools`. We corrected the output from `vcftools` to exclude sites at which we could not call variants using our pipeline. Specifically, rather than use 10 kb for the denominator, we used the number of "callable sites" (see 2.5) within the 10 kb window and excluded windows for which there were fewer than 1000 "callable sites".

```
vcftools --window-pi 10000 --window-pi-step 5000 \\
    --gzvcf {HC_HWE_Unrelated.recode.vcf.gz} --out {HC_HWE_Unrelated_pi}
```

##### 4.4 Inferences of population demographic history

We used `smc++`[53] v1.15.1 to estimate the effective population sizes over time in the lineage leading to extant corn snakes. To distinguish tracts of homozygosity from regions with no data or low depth of coverage in our filtered VCFs, we scanned the median coverage in windows of 5 kb along the autosomes of all individuals using:

```
mosdepth -m -n -x -Q 30 -b 5000 {SAMPLE} {SAMPLE.bam}
```

In each individual, we masked all 5 kb windows with coverage below half the mean value for all autosomal windows.

Input files for `smc++` were prepared from VCF files, excluding all regions that did not meet the coverage threshold and had been determined to be uncallable in section 2.5.1. `smc++` requires one individual in the run to be specified as the "distinguished" individual, with the remaining individuals only contributing allele frequency information. To this end, we generated five sets of input files, using one of the five samples with highest average genome-wide coverage as the "distinguished" individual in each:

```
smc++ vcf2smc {HC_HWE_Unrelated.recode.vcf.gz} {Sample.Scaffold.SMC.input} \\
    {Scaffold} {All_samples} -d {Sample} {Sample} --length {Scaffold_size} \\
    -m {Callability.Mappability.mask.bed}
```

Following the recommendations in https://github.com/popgenmethods/smcpp, we then estimated the demographic history of corn snakes by providing the five input sets with different distinguished individuals, assuming a mutation rate of $4 \times 10^{-9}$ per site per generation. During the preparation of this manuscript, Bergeron et al 2023[54] reported mutation rates estimated from a variety of vertebrates; we note that, while the snake mutation rate reported in that paper is a clear outlier, $4 \times 10^{-9}$ per site per generation is close to the mode of the mutation rates reported in the study.

```
smc++ estimate -o {SMC.output} 4e-9 {*.SMC.input}
```

In addition, to double-check the demographic inference by `smc++`, we ran `msmc` v1.0.0 for 20 corn snake individuals, obtaining similar results. To obtain the input files required by `msmc`, we applied the same callability and mappability mask described above for `smc++`, and used the scripts in `msmc-tools` (https://github.com/stschiff/msmc-tools) to convert VCFs to the required format:

```
python msmc-tools/generate_multihetsep.py {HC_HWE_Sample.recode.vcf.gz} \\
    --negative_mask {Callability.Mappability.mask.bed} --chr {Scaffold} > \\
    {Sample.Scaffold.MSMC.input}
```

We then ran `msmc` for each individual using:

```
msmc -o {Ssample.MSMC.output} {Sample.*.MSMC.input}
```

### 5 Inference of LD-based recombination maps

We tried two distinct methods for inferring LD-based recombination maps: `LDhelmet`[41], which is model-based and assumes a vanilla demographic model, and `Pyrho`[55], a deep learning method that relies on a previously inferred demographic history.

#### 5.1 LDhelmet

`LDhelmet` requires several inputs related to the population genetic parameters of the sample, including an estimate of $\boldsymbol{\theta}$, ancestral allele calls for variants in the population, and a mutation transition matrix.

##### 5.1.1 $\theta$ and long-term effect population size ($N_e$) estimation

We used the "high coverage unrelated" set of variants to estimate $\boldsymbol{\theta}$ using Watterson's estimator[56], $\hat{\boldsymbol{\theta}}_w = K/a_n$, where $K$ is the number of segregating sites, $n$ is the number of haplotypes, and $a_n$ is the number of chromosomes. The number of sites in overlapping 10 kb windows with a 5 kb step size was found using `bedtools`, and this value was divided by the number of "Callable sites" within the 10 kb window, excluding regions for which there were fewer than 1,000 "callable sites" (see section 2.5).

```
bedtools makewindows -s 5000 -g {Fasta.genome} -w 10000 | bedtools coverage \\
    -sorted -a stdin -b {HC_Unrelated_NoSingle.vcf.gz} > \\
    {NumberOfVariableSites_w10kb_s5kb.bed}
```

By this approach, the median $\boldsymbol{\theta}$ estimate was 0.0037 per base-pair and the mean 0.0040 (Figure ??). Because singletons are not considered by LDhelmet, we also calculated this estimator after excluding singletons: the median $\boldsymbol{\theta}$ value was 0.0027. Given a mutation rate of $4 \times 10^{-9}$ per base-pair per generation, this suggests a long-term estimate of the effective population size $N_e$ of 250,000.

##### 5.1.2 Ancestral alleles and mutation transition matrix

LDhelmet takes an optional input file describing the probability of each base (A,C,G,T) being the ancestral allele at each site. These ancestral allele probabilities are typically derived from comparisons to the genomes of closely related species. We used a multi-species alignment that we had previously generated using cactus[57], including corn snakes, the black rat snake, *Pantherophis obsoletus*, the most closely related available species (diverged roughly 11MYA), and seven other snakes[58] to identify the predicted genome sequence of the common ancestor of *P. guttatus* and *P. obosletus* and polarize ancestral and derived alleles. Because the split between these two species was likely >10MYA, we checked the concordance between these ancestral allele calls and the major allele in the sample. The two agree for 80.2% of variants. At these sites, we assigned the base corresponding to the ancestral/major allele a 97% probability of being the ancestral allele and each of the three remaining bases a 3%. For sites at which there is no predicted ancestral sequence from the multi-species alignment (4.3%) or sites where the ancestral sequence and the major allele are not the same (15.5%), we assigned a 47% probability that each of the segregating alleles is the ancestral allele and a 3% probability to the two other bases.

To calculate the mutation transition matrix, we counted the number of times that we identified each mutation type (i.e. the number of observations of ancestral A mutated to derived C, A to G, etc.) and the number of times that we found each base at a "callable site" in the corn snake genome. These values were used to calculate the estimated mutation transition matrix, following[41].

##### 5.1.3 Running LDhelmet

After using bcftools and vcftools to remove singletons and convert the phased variants into the appropriate file formats, LDhelmet was run on all scaffolds larger than 100 kb using our 24 high coverage unrelated samples. We used recommended settings for the preparation steps, except for the range of $\boldsymbol{\rho}$ values in the likelihood table calculation, as we expected $\boldsymbol{\rho}$ to be much lower for corn snakes than for *Drosophila*, for which the recommended range of $\boldsymbol{\rho}$ values were generated. We used a $\boldsymbol{\theta}$ value (without singletons) of .00268 and ran the rjmcmc for $10^6$ steps with a burn in of 100,000 steps and using block penalties of 10, 20, and 50.

```
bcftools view -s {Unrelated_Samples} -O u {SCAFFOLD_PHASED.vcf} | bcftools view \\
    -c 2:minor -O v | vcftools --ldhelmet --chr {Scaffold} --vcf - --out {Scaffold}

ldhelmet find_confs --num_threads 4 -w 50 -o {Output_configurations} \\
    {Scaffold*.snpfile}

ldhelmet table_gen --num_threads 16 -c {Output_configurations} -t 0.00302 \\
    -r 0.0 .0000001 .000001 .000001 .00001 .00001 .0001 .0001 .001 .001 .01 .01 \\
    .1 .1 1.0 1.0 10.0 -o {Output_likelihood}

ldhelmet pade --num_threads 16 -c {Output_configurations} -t 0.00302 -x 11 \\
    -o {Output_pade}

ldhelmet rjmcmc --num_threads 2 -w 50 -l {Output_likelihood} -p Output_pade} \\
    -b {block_penalty} --pos_file {SNP_Positions} --snps_file {SNP_Haplotypes} \\
    -m {Mutation_matrix} --burn_in 100000 -n 1000000 -a {??} -o {RJMCMC_out.post}

ldhelmet post_to_text -m -p 0.025 -p 0.50 -p 0.975 -o {RJMCMC_out.txt} \\
    {RJMCMC_out.post}
```

##### 5.1.4 Filtering the LDhelmet output

The output of LDhelmet was filtered to exclude contig boundaries +/− 1000 bp. We also removed regions of the maps with implausibly high recombination rates, using >1 centimorgan per hotspot as a cut-off. We estimate the long term $N_e$ to be roughly 250,000. For hotspots of length 2 kb, a value of 1 centimorgan per hotspot would correspond to roughly $\boldsymbol{\rho} = 5$ per bp. This filter removed 71 regions, covering 0.0096% of the genome, from the recombination map, and the filtered maps were used to generate mean recombination rates in windows of 100 bp, 1 kb, 10 kb, 100 kb, and 1 Mb.

#### 5.2 Pyrho

Pyrho was run with recommended settings, the demographic history model inferred in 4.4, and a mutation rate of 4 $\times 10^{-9}$ per base-pair. The Pyrho make_table function calculates a lookup table using a Moran population size of the sample; following the manual, which recommends using a value between $1.25x$ and $1.5x$ the sample size and given our sample size of 48 chromosomes (24 samples), we used $48 * 1.35 = 65$. Based on the Pearson correlations between simulations reported by Pyrho hyperparam, we used a window size of 70 and a block penalty of 15 for Pyrho optimize.

```
make_table --samplesize 48 --approx --moran_pop_size 65 --mu .000000004 \\
    --outfile {Pyrho_lookup.txt} --smcpp_file {Demographic_model.csv}

hyperparam --samplesize 48 --tablefile {Pyrho_lookup.txt} --mu .000000004 \\
    --ploidy 2 --smcpp_file {Demographic_model.csv} --outfile {Pyrho_hyperparam.txt}

optimize --vcffile {HC_HWE_Unrelated.recode.vcf.gz}  --windowsize  70 \\
    --blockpenalty 15 --tablefile {Pyrho_lookup.txt} --ploidy 2 --outfile {Pyrho_map.txt}
```

The resulting map was filtered as in 5.1.4.

#### 5.3 Assessing the robustness of the inclusion of two population samples

Out of concern that even the low levels of population differentiation between our colony and wild-caught samples might bias estimated recombination rates, we reran Pyrho(section 5.2) on only the 19 wild-caught samples, after re-inferring the demographic model (section 4.4). We compared the resulting maps to the those generated using the full sample of 24 individuals: the Pearson correlation was 0.863 at the 1 kb scale, 0.846 at the 10 kb scale, 0.874 at the 100 kb scale, and 0.900 at the 1 Mb scale. Given the high concordance, we relied on the full sample of 24 individuals for downstream analysis.

#### 5.4 Calling hotspots

Hotspots were called using a custom python script, which scanned across the genome in 2 kb windows, overlapping by 1 kb. We calculated the mean recombination rate in each window, as well as the background recombination rate, i.e., the mean recombination rate in the 20 kb flanking the window, excluding a buffer region of 1 kb on either side of the window (ref). Any window in which more than half of the central window or more than half of the background had been filtered out was excluded. For each remaining window, we found the "heat" of a hotspot by dividing the mean recombination rate by the mean background recombination rate; hotspots were defined as windows with heat $> 5X$ or $> 10X$; hotspots found within 5 kb of one another were merged. Qualitative results were similar using heat 5X or 10X as a cut-off; to increase the number of hotspots considered, we used 5 as the cutoff in the main analyses.

The number of LDhelmet hotspots identified at a given block penalty and heat were:

| LDhelmet block penalty | Number of hotspots at heat 5 | Number of hotspots at heat 10 |
|---|---|---|
| 10 | 16,923 | 7,172 |
| 20 | 14,434 | 6,133 |
| 50 | 10,323 | 4,334 |

The number of `Pyrho` hotspots identified were:

| Number of hotspots at heat 5 | Number of hotspots at heat 10 |
|---|---|
| 9,272 | 2,970 |

#### 5.5 Comparison of LDhelmet and Pyrho recombination maps

We compared our recombination maps from `LDhelmet` and `Pyrho` using several methods. First, we looked at the Pearson correlation between the `LDhelmet` map and the `Pyrho` map at each scale. The results are reported below.

| `LDhelmet` block penalty | 1 kb scale | 10 kb scale | 100 kb scale | 1MB scale |
|---|---|---|---|---|
| 10 | .326 | .366 | .464 | .495 |
| 20 | .384 | .493 | .709 | .837 |
| 50 | .477 | .645 | .838 | .926 |

We also considered the fraction of hotspots identified in the `LDhelmet` and `Pyrho` maps.

| `LDhelmet` block penalty | Fraction of `LDhelmet` hotspots found in `Pyrho` |
|---|---|
| 10 | .405 |
| 20 | .450 |
| 50 | .549 |

While the /texttttLDhelmet block penalty of 50 resulted in the highest Pearson correlation between the `LDhelmet` and `Pyrho` maps, it also resulted in 30% fewer hotspots than the block penalty of 20. We decided to use the block penalty of 20, which both had a moderately high Pearson correlation with the `Pyrho` maps (.493 at 10 kb scale) and led to the identification of substantially more hotspots. Then, we sought to understand the various properties of hotspots inferred by `LDhelmet` and `Pyrho`. To this end, we used a custom Python script that computes several statistics in every 1 kb window within 15 kb of any hotspot, using the information annotated in the VCF files by `GATK` or based on the genotype data (i.e., mapping quality, rank sum test of REF versus ALT base quality scores, minor allele frequency, depth of coverage, fraction of reads supporting the REF allele, number of SNPs, fraction of unmappable base pairs, nucleotide diversity and fraction of missing genotypes). Some variables, such as allele frequency and $\pi$, are higher in `Pyrho` hotspots than in the surrounding regions, and these relationships are largely absent from hotspots called by `LDhelmet` (Figure S6). While these relationships could be real, a concern is that by `Pyrho` sometimes infers an elevation in the population recombination rate when there is a deep gene genealogy rather than elevated recombination rates. Since `LDhelmet` hotspots do not have those properties to the same extent, and `LDhelmet` infers many more hotspots than `Pyrho`, we decided to rely primarily on the `LDhelmet` map for our downstream analyses. However, we conducted several downstream analyses using both, and found that our results were qualitatively similar (Figure S7).

As an additional filter on the `LDhelmet` recombination maps, we removed 10 kb windows that differed by more than an order of magnitude between `LDhelmet` and `Pyrho` maps; this led to the exclusion of 2.2% of the physical genome. The resulting maps were used for all downstream analyses.

### 6 Relationship between recombination rates and other genomic features

#### 6.1 Heterozygosity and GC content

To assess if our recombination map estimates are sensible, we checked if GC content increases with higher estimated recombination rate, as expected from other species[59]. To do so, we calculated GC content in 10 kb bins using `bedtools`. We then classified each bin into deciles based on GC content and plotted box and whisker plots showing the distribution of recombination rates (Figure S8).

```
bedtools makewindows -g {REFERENCE.genome} -w 10000 > {REFERENCE_10kbBins.bed}

bedtools nuc -fi {REFERENCE} -bed {REFERENCE_10kbBins.bed} > \\
    {{REFERENCE_10kbBins_nuc.bed}
```

#### 6.2 PRDM9 binding sites

To exclude the possibility that the association between computationally predicted PRDM9 binding sites (identified in section 3.2) and recombination rates is driven by confounding effects of CpG islands or transcription start sites (TSS), we restricted our analysis to genomic regions farther than 10 kb from either promoter-like feature, using `bedtools intersect`. We then used `bedtools closest` to find the PRDM9 binding site nearest each genomic location.

```
bedtools intersect -v -a {Filtered_LDhelmet_Output.bed} -b \\
    {CpGIslands_PlusMinus10kb.bed} | bedtools intersect -v -a stdin \\
    -b {TSS_PlusMinus10kb.bed} | bedtools closest -d -a stdin \\
    -b {PRDM9_Binding_Sites.bed} -t first > {DistanceToFiltered_LDhelmet}
```

Specifically, we used a custom python script to group 100 base-pair bins by their distance to the nearest predicted PRDM9 binding site, find the mean recombination rate at each distance, and bootstrap with replacement 100 times. The resulting mean recombination rate versus distance to the nearest predicted PRDM9 binding site curves were smoothed using `Statsmodels lowess`, and the 95% confidence interval was plotted by bootstrap resampling of 100 base-pair windows within a bin. We performed this analysis using the union of all predicted PRDM9 binding sites, as well as for the shared zinc fingers found in section 3.3.

#### 6.3 CpG islands and transcription start sites (TSS)

To examine the association between TSS/CpG islands and recombination rates, we excluded genomic regions that are within 10 kb of a predicted PRDM9 binding site for any PRDM9 allele. We then proceeded analogously to as in section 6.2.

#### 6.4 Recombination modifier model

We followed Spence and Song (2020)[?] in modeling the effects of PRDM9 binding and promoter-like features jointly on recombination rates, with GC content (ref) and background recombination rate as covariates. The effects of these features on recombination rate were estimated using a linear model, where the response variable was the log10 recombination rate in 1 kb bins. These windows were thinned to one every 10 kb, as genomic windows in close proximity to each other are not independent. The predictors were binary variables indicating the presence or absence of a predicted PRDM9 binding site, CpG island, or TSS within each genomic window. Two interaction terms were also included: the first indicating the presence of both a CpG island and a TSS, and the second the presence of a predicted PRDM9 binding site and either promoter-like feature. Finally, two covariates were used: the log10 background recombination rate in the surrounding 1 Mb, as we were interested in features changing the relative recombination rate, and GC content as calculated by `bedtools nuc` as in 6.1.

The model was implemented using the statsmodels Ordinarly Least Squares (OLS) function and was trained on the genome as a whole, on scaffolds aligning to macrochromosomes, and scaffolds on microchromosomes as identified in section 1.6.2.

### 7 Recombination hotspot analyses

#### 7.1 Calling coldspots

To create a set of "coldspots" with which to compare our hotspot regions, we matched each hotspot with a genomic region meeting several criteria: it had to have the same width and GC content (within 1%, as reported by `bedtools nuc`) as the hotspot; be found within 5 MB of the original hotspot location but not within 10 kb of any hotspot or other coldspot, and have an estimated recombination rate less than half of the scaffold-wide average.

#### 7.2 Overlap with PRDM9 binding sites and promoter-like features

To determine the overlap between recombination hotspots (and coldspots) and features of interest, such as predicted PRDM9 binding sites and promoter-like features (CpG islands and TSS), we utilized the following code:

```
bedtools closest -a {Hotspots.bed} -b {PRDM9_binding_sites.bed} -d -t first | \\
bedtools closest -a - -b {Promoter-like.bed} -d -t first \
> {Hotspots_distance2features.bed}
```

We considered the features of interest to overlap with a hotspot when the distance between them was at most 500 base-pairs. We also used these distances to condition on hotspots that are far (> 10kb) from a given feature.

To determine if the observed overlap is different than that expected by chance, we randomly relocated each hotspot within a neighboring 5 Mb region, avoiding any overlap with other hotspots or gaps in the sequence. The code for this procedure is as follows:

```
cat {ContigBoundaries_gaps.bed} {Hotspots.bed} | bedtools sort -i - \\
> {Regions_to_exclude.bed}
```

We then iteratively moved the hotspot coordinates and updated the excluded regions:

```
while read hotspot_interval;do
    slop=$(echo "$hotspot_interval" | bedtools slop -i /dev/stdin -g $scaf_size \
    -b 2.5e6;
    shuf=$(echo "$target_interval" | bedtools shuffle -i - -g $scaf_size \
    -noOverlapping -incl <(echo $slop) -excl {Regions_to_exclude.bed});
    echo -a "$shuf" >> {Regions_to_exclude.bed};
done < {Hotspots.bed} > {Shuffled_Hotspots.bed}
```

We repeated this procedure 500 times and calculated the proportion of shuffled hotspots that overlapped with the features of interest.

#### 7.3 Hotspot motif enrichment analysis

In order to determine whether there are short motifs of 7-13 bases enriched in recombination hotspots compared to coldspots (which would be bound by 2-4 zinc fingers), we used `Streme`[60] from `Meme Suite`, considering only hotspots and coldspots far from CpG islands and TSS as identified in section 7.2.

```
streme --minw 7 --maxw 13 --patience 10 -bfile {Ref_Background.txt} \\
    -oc {Motifs_Hotspots} -n {Coldspots_FarCpGI.fa} -p {Hotspots_FarCpGI.fa}
```

We assessed whether the resulting motifs matched any of our predicted PRDM9 binding motifs using the R package `universalmotif`[61]. We used the `compare_motifs` function to calculate the Pearson correlation coefficient between each motif enriched in hotspots and computationally-predicted PRDM9 binding motif. We required at least seven bases of overlap between motifs. The results were plotted using the `pheatmap` R package https://github.com/raivokolde/pheatmap.

```
library(pheatmap)
library(universalmotif)
m <- read_uniprobe({PRDM9_PWMs.txt})
m2 <- read_meme("Streme_Motifs.txt")
allm <- append(m, m2)
rows <- {List of PRDM9 allele names}
cols <- {List of streme motif names}
mofs = compare_motifs(allm, method="PCC", min.overlap=7)
mofs = mofs[rows,cols]
pheatmap(mofs, treeheight_row = 0, treeheight_col = 0)
```

## 7.4 GC*

We analyzed the nine snake whole genome alignment as in section 5.1.2. For each hotspot (defined as having a heat of at least 5, see 5.4) and matched coldspot (see 7.1) in corn snakes, we generated a Multiple Alignment Format (MAF) file with 5 kb added to each end using the `hal2maf` tool in `halTools`[62]:

```
hal2maf {WG_alignment.hal} {Spot.maf} --refGenome {Pantherophis_guttatus} \
--refSequence {scaffold} --onlyOrthologs --noDupes \
--start {start} --length {length}
```

We then estimated GC*[63], the expected equilibrium GC content, using a custom python script. GC* was calculated in 100 bp windows from the center of the hotspot (or matched coldspot) as:

$$\frac{\frac{AT \to GC}{ancAT}}{\frac{AT \to GC}{ancAT} + \frac{GC \to AT}{ancGC}} \tag{1}$$

where ancAT and ancGC are the number of As and Ts or Gs and Cs, respectively, in the ancestral sequence, as estimated by `cactus`, and AT>GC and GC>AT represent the number of substitutions on a lineage from an A or T to a G or C, or from a G or C to an A or T, respectively. To explore if the conservation level of hotspots is affected by distance to promoter-like features, we classified hotspots as close (<5 kb away) and far (>10 kb away) from TSS and CpG islands in the corn snake genome (see 1.5.2).

### 8 Testing for excess divergence at PRDM9 binding sites in snakes

One way of determining whether PRDM9 is active in localizing recombination events is to look for lineage-specific excess divergence at PRDM9 binding sites, which arises from the over-transmission of sequences bound more weakly by PRDM9 binding sites at heterozygous sites. To look for signals of increased divergence at PRDM9 binding sites along the corn snake lineage, we used the nine snake whole genome alignment as in section 5.1.2, which includes corn snake and the closely related black rat snake (*Pantherophis obsoletus*). We used `hal2fasta` (from `Comparative Genomics Toolkit`) to generate a `fasta` file containing the aligned genomes of corn snakes, rat snakes, and garter snakes (*Thamnophis elegans*), which we used as an outgroup. The aligned genomes were divided into short segments of variable length, which contained uniquely aligned sequence from all three species with no Ns, insertions, or deletions. These genome segments were scanned for PRDM9 binding motifs using `Fimo` from `Meme Suite` as in section 3.2 to identify predicted PRDM9 binding sites for each corn snake PRDM9 allele and for the shared zinc fingers identified in section 3.3.

```
hal2fasta --outFaPath {Ancestral.fa} {Snake_alignment.hal} {Pantherophis_Node_Name}

hal2maf --refTargets {Ancestral_Binding_Sites.bed} --refGenome \\
    {Pantherophis_Node_Name} {Snake_alignment.hal} --targetGenomes \\
    {Pantherophis_obsoletus,Pantherophis_guttatus} {Output.maf}
```

Using a custom python script, we then identified genomic positions at which PRDM9 binding sites were gained or lost in either corn snake or rat snake lineages. For example, a site was classified as gained in corn snake if it was identified as a binding site in corn snake, but not at the corresponding position in the rat or garter snake genomes, while a site was considered lost in corn snakes if it was found in garter and rat snake but not corn snake.

Similarly, we quantified the number of loses and gains for the nine motifs that are significantly enriched in hotspots (see 7.3). For every motif, we replaced any positions within the position weight matrix (PWM) with a maximum weight <80% by an N, i.e., we did not consider such positions when finding matches between the motif and the ancestral, corn snake, or black rat snake sequence. One of the motifs (AAAAAAAAAAAC, using the highest weights in the PWM) showed an extremely low number of hits in all sequences (< 5 instances), most likely explained by the masking of repetitive sequences in the whole-genome alignment of snakes[58]. Given the low numbers, the quantification of gains and loses is extremely noisy and hard to interpret; thus we excluded this motif from the analysis.

### 9 Crossovers

#### 9.1 Calling crossovers

To identify recombination events in corn snake pedigrees, we sequenced the genomes from two nuclear families that provide information about 24 meiosis events (see section 2.1), and implemented a previously described algorithm[64] to call crossovers. The algorithm uses the transmission of "informative markers" (positions that are heterozygous in one parent but not in the other) to phase the haplotypes inherited by the offspring and identify crossover events in the germline of the parents. For each parent in each family, each offspring is considered in turn as the "template" individual. The alleles in the non-template offspring are re-coded to "1" if they match the parental allele transmitted to the template, "2" if they match the untransmitted allele, or "0" if they have a missing genotype. After this re-coding, positions at which a non-template offspring changes from copying one allele to the other are defined as a "switch" (e.g., sequences like "..1 1 2 2.."). We called putative crossovers in the template individual as intervals at which the majority (in practice, usually all) of the non-template individuals with genotype calls showed a switch.

This procedure is susceptible to genotyping errors, e.g., missed heterozygotes in the template individual, which generate changes of phase in close physical succession. To address this issue, we grouped putative crossovers within ten informative markers of each other in each of the template individuals, and kept only those clusters with an odd number of changes of phase. In addition, we also removed crossovers within ten informative markers of the edges of the autosomal scaffolds. Finally, we removed crossovers in different template individuals occurring within 1 kb of each other. While in principle, this step could remove a few true positives, such events are unlikely given the small number of meiosis analyzed, and are much more likely the product of inaccurate genotyping at repetitive regions.

By this procedure, we called 324 crossovers, consistent with what might be expected given 24 meioses (in 10 offspring) and the karyotype of corn snakes (17 autosomes), assuming 1-2 crossovers per chromosome, half of which are visible in transmissions.

##### 9.2 Overlap with features of interest

The overlap between crossovers and features of interest was calculated as described in section 7.2. In addition, we checked that the shuffled crossover locations spans at least two informative markers, such that calling a crossover would in principle be possible.

### 10 Rattlesnakes

Schield et al. 2020[26] previously published an analysis of features correlated with increased recombination rate in the prairie rattlesnake *Crotalus viridis*, reporting increased recombination at CpG islands. The authors further found that the predicted binding motif for three zinc fingers of the PRDM9 reference sequence was enriched at promoter-like features. On that basis, they argued that PRDM9 directs recombination to promoters in rattlesnakes. We sequenced six complete PRDM9 alleles from *C. viridis* and analyzed the binding sites of these alleles in conjunction with the Schield *et al.* (2020) *C. viridis* recombination map. The accession information for the *C. viridis* genome assembly can be found at `https://figshare.com/projects/Prairie_rattlesnake_Crotalus_viridis_genome_assembly_and_annotation/66560` and the recombination map[26] is at `https://figshare.com/articles/Rattlesnake_Recombination_Maps/11283224`.

##### 10.1 Sequencing PRDM9

We sequenced PRDM9 alleles from three wild caught samples of *C. viridis* kindly provided to us by Dr. Edward Myers from the American Museum of Natural History. Genomic DNA was extracted from the provided tissue samples as in section 2.1. We PCR amplified the zinc finger domain using primers described in Table S7, sequenced and assembled the resulting PCR products using Oxford Nanopore sequencing and canu as in section 3.1, and identified binding sites of these PRDM9 alleles as in 3.2.

##### 10.2 Recombination map and association genomic with features

We relied on the *Crotalus viridis* recombination map published by Schield *et al.* (2020)[26], which was generated using `LDhelmet`[41], and filtered out regions with implausibly high recombination rates as in section 5.1.4. Rattlesnakes have a $\hat{\boldsymbol{\theta}}_w$ value of .005 per base-pair[26], and so $\boldsymbol{\rho} = 6$ per bp is roughly 1 centimorgan per hotspot. We were unable to remove regions near contig boundaries due to the short contig N50 of the *Crotalus viridis* genome assembly (14.8KB)[65]; removing contig boundaries and 1 kb on either side from the recombination map, as done in section 5.1.4, resulted in the exclusion of more than 27% of the genome and over 80% of the hotspots reported in Schield et al. 2020.

In the absence of methylation information, we identified *C. viridis* CpG islands using the primary genome sequence, as in section 1.5.2. We examined the association of CpG islands and PRDM9 binding sites with recombination rate in the same way as in sections 6.2 and 6.3.

##### 10.3 Evaluating genome features at CpG islands

To evaluate the quality of the corn snakes and rattlesnake assemblies around CpG islands, which are repetitive in nature, we quantified mapping quality measures in the genomic regions surrounding them. To anchor our comparison between corn snakes and rattlesnakes, we also included garter snakes, for which the Vertebrate Genome Project has produced a high quality, chromosome-level assembly (`https://www.ncbi.nlm.nih.gov/assembly/GCF_009769535.1/`). For each assembly, we mapped short-read sequencing of the corresponding reference individual following the steps described in section 2.2. To allow for a straightforward comparison across assemblies, we identified CpG islands *de novo* using `cpgplot` with default parameters, as described in section 1.5.2. We then looked at the fraction of base-pairs covered, depth of coverage, and mean mapping quality around the CpG islands identified in each assembly using `samtools coverage` (Figure S14).

Given the high fragmentation of the rattlesnake genome, a possible explanation for the higher fraction of positions covered, depth of coverage, and mapping quality in rattlesnakes may be that paralogous sequences, such as the ones characteristic of CpG islands, are sometimes mapped into a single location in the reference genome. Such mapping errors present a challenge for analyses of LD-based recombination maps at CpG islands, as they will distort and likely decrease levels of linkage disequilibrium, resulting in upwardly biased estimates of recombination rates.

### 11 ZCWPW2 binding assay

The coding sequences of corn snake ZCWPW2 was identified based on the gene annotation and confirmed using the PacBio CCS reads collected in section 1.4.1. The gene was synthesized by GenScript (Piscataway, NJ).

To isolate the coding sequence of mouse ZCWPW2, we extracted total RNA from mouse testes provided to us by Franck Polleux (Columbia University) and stored in RNAlater (Thermo Scientific) at −80°C. RNA extraction was performed using an RNeasy kit as specified by the manufacturer using the hand-held pestle grinding and with the optional DNase step included (Qiagen, Valencia, CA). We synthesized cDNA from total RNA using the GoScript Reverse Transcription System kit as specified by the manufacturer with reaction volumes doubled (Promega, Madison, WI). We conducted PCR reactions from cDNA using LongAmp Taq with an annealing temperature of 66C and 30 cycles (New England Bio Labs, Ipswich, MA) and using primers designed to amplify the coding sequence of ZCWPW2 using NCBI Primer blast (Ye et al. 2023). We used the TOPO Blunt end Cloning kit to perform a cloning reaction, transform competent E. coli cells, and select colonies with the desired sequences as specified by the manufacturer (ThermoFisher Scientific, Waltham, MA). We used the QIAprep Spin Miniprep Kit to purify plasmid DNA as specified by the manufacturer (Qiagen, Valencia, CA). Plasmids were sequenced with Sanger sequencing at the Stanford University Protein and Nucleic Acid Facility to confirm that they contained the required sequences.

The coding sequences of ZCWPW2 from both mouse and corn snake were cloned into pcDNA3.1(+)-C-HA backbone vector to express ZCWPW2 protein fused with an HA-tag at the C-terminus (GenScript). These expression vectors were used for in vitro translation of HA-tagged mouse ZCWPW2 (mZCWPW2) and corn snake ZCWPW2 (sZCWPW2) using the TnT® Quick Coupled Transcription/Translation System for T7 promoter (Promega). For each reaction, 2 µg of plasmid DNA was used in a 50 µl reaction, and 1 µl of the translation product was utilized for Western blotting to confirm equal expression of mouse and snake proteins. Additionally, 20 µl of the translation product was used for histone peptide pull-down.

Biotinylated Histone 3 peptides with various modifications, including H3.1 aa1-43 non-modified, H3.1 aa1-43 K4me3, H3.1 aa1-43 K36me3, and H3.1 aa1-43 K4me3 K36me3, were obtained from EpiCypher. For each peptide binding reaction, 1 $\mu$g of H3 peptide was mixed with 20 $\mu$l of recombinant ZCWPW2 protein in 250 $\mu$l of binding buffer (50 mM Tris pH 7.5, 300 mM NaCl, 0.25% NP-40) and rotated at 4°C for 4 hours. Streptavidin Dynabeads™ M-280 (Invitrogen #11205D) were washed twice in 1 ml of binding buffer and resuspended in 25 $\mu$l of binding buffer. After the 4-hour incubation of H3 peptide and recombinant ZCWPW2 mixture, the resuspended beads were added to the mix and incubated for an additional hour with rotation at 4°C. Following incubation, the beads were washed three times with 1 ml of binding buffer and resuspended in 20 $\mu$l of 2x SDS sample buffer (Invitrogen). The beads-SDS mixture was then incubated at 70°C for 10 minutes and loaded onto an SDS PAGE Protein Gel (NuPAGE 4–12% Bis-Tris) in MES buffer (ThermoFisher). The proteins were subsequently transferred to nitrocellulose membranes and blocked overnight at 4°C in TBST with 5% milk. Primary antibody immunoblotting was performed for 1 hour at room temperature using an anti-HA rabbit antibody (Abcam ab9110, 1 $\mu$g/$\mu$l, 1:2000 dilution), and secondary immunoblotting was conducted using a goat anti-rabbit HRP (Thermofisher, 1/4000 dilution). The proteins were detected by enhanced chemiluminescence using a c600 Azure imager (Azure Biosystems).

### 12 Conservation analyses

#### 12.1 Compiling reptilian alignments

##### 12.1.1 ZCWPW1 and ZCWPW2 orthologs

To identify reptilian ZCWPW1 and ZCWPW2 orthologs, we followed a similar protocol to Cavassim *et al.* (2022)[66]. Using human MANE select transcripts downloaded from Ensembl release 109[67](translation IDs ENSP00000507762.1 and ENSP00000373278.2), we first performed `blastp` searches against the NCBI RefSeq database[68] and downloaded 500 GenPept files from reptilian taxa, filtering out Aves. We performed a similar search against the NCBI non-redundant protein sequence database, downloading only those files with >55% coverage and >40% identity. To these, we added representative sequences from a blast search against UniProt's UniRef100 database,[69] downloading representative reptilian sequences from the top 1000 hits. Finally, orthologs from species otherwise missing from our analysis were added from sequences annotated by Ensembl's gene tree pipeline. We removed duplicate sequences using `SeqKit's rmdup -s`[70] then annotated the sequences for the presence of zf-CW and PWWP domains using NCBI's `Batch-CD Search`[71,72] with an e-value cutoff of 1, drawing on NCBI-curated, Protein Clusters, Protein Families,[73] COG,[74] and TIGRFAMs databases[75]. Sequences were then divided into two groups: those with both zf-CW and PWWP domains and those with only one. Sequences with no domain annotations were discarded.

In `Geneious 2022.2.2`, we then aligned the sequences using `MUSCLE`[76] (default parameters) and extracted zf-CW and PWWP domains for domain-specific phylogenetic trees built with `RAxML`[77]. We used these trees to remove sequences that clustered with ZCWPW paralogs from the NSD and MORC families (MANE select or Ensembl canonical sequences from Ensembl). We then built full sequence trees using the remaining genes to differentiate ZCWPW1 from ZCWPW2. From those sequences with both domains, we obtained 83 complete, unique ZCWPW1 sequences for 32 reptilian species and 61 complete, unique ZCWPW2 sequences for 39 reptilian species. For alignments used in subsequent analyses, we chose one transcript per species based on its completeness and length. Final alignments were prepared using `MUSCLE`.

##### 12.1.2 PRDM9 orthologs

Similarly, PRDM9 orthologs were obtained via blast searches against NCBI RefSeq, non-redundant, and UniRef100 databases. As a query, we used the human MANE select PRDM9 sequence (downloaded from Ensembl, translation ID ENSP00000296682.4), truncated to remove the highly variable zinc finger domain. Duplicate sequences were removed using SeqKit, and domains were annotated using NCBI's Batch-CD search. We additionally performed a blast search against incompletely annotated reptilian sequences using SSXRD and KRAB domains from a close relative. This approach allowed us to identify additional complete sequences missed in the previous step. Sequences were grouped based on their completeness, and only those possessing SSXRD, KRAB, and SET domains were used in further analyses.

As before, we then aligned the sequences using `MUSCLE` and built domain-specific phylogenetic trees with `RAxML`, in order to remove paralogs of PRDM9 such as other members of the PRDM family and members of the SSX and KRAB-ZFP families. From this procedure, we obtained 24 complete reptilian PRDM9 sequences, which we later supplemented with an additional four snake sequences extracted from whole genome sequences in Cavassim *et al.* (2022).

###### 12.1.3 ANKRD31 orthologs

ANKRD31 orthologs were obtained via blast searches against NCBI RefSeq, non-redundant, and UniRef100 databases. In all cases, the human MANE select protein sequence was used as a query (translation IDs ENSP00000427262.2 and ENSP00000285106.6, respectively) and only sequences annotated as ANKRD31 were downloaded. Duplicate sequences were removed using SeqKit. We next built trees using `RAxML` and removed transcripts that grouped with human paralogs of the target genes such as ANKRD11 and ANKRD12. Final `MUSCLE` alignments were prepared using one sequence per species, chosen based on length and completeness.

#### 12.2 PAML *codeml* conservation analysis

`PAML's codeml` branch models allow estimation of dN/dS (w) across branches in a phylogeny, as well as likelihood-based comparison of models where these ratios are fixed for certain branches[78]. Because genes evolving under purifying selection are expected to have $\omega < 1$, we used `codeml` to test whether ZCWPW1, ZCWPW2, PRDM9, and ANKRD31 were conserved in snakes, i.e., if we could reject the null hypothesis that $\omega = 1$ for this lineage.

Where possible, coding sequences corresponding to the most complete protein sequence obtained in the ortholog analysis were downloaded from NCBI RefSeq or Ensembl. Otherwise, coding sequences from taxa with a confirmed copy of the gene were downloaded from orthologs identified by NCBI's Eukaryotic Genome Annotation pipeline. Mammalian coding sequences were also downloaded from NCBI Orthologs for comparison. Sequences were aligned and filtered using `Guidance2`[79] (default parameters, removing columns with an alignment score $< .93$). Prior to filtering in Guidance, codon-aware alignments of ANKRD31 sequences(prepared using `Pal2Nal`[80]) were additionally trimmed using `trimAl` to remove sites present in fewer than a third of the sequences (`trimAl -gt .33`)[81]. Zinc fingers were removed from PRDM9 sequences. We excluded taxa with confirmed absences of PRDM9 when analyzing ZCWPW1, ZCWPW2, and ANKRD31 and those missing ZCWPW1 or ZCWPW2 when analyzing PRDM9.

For each reptilian alignment, we first allowed two $\omega$ values: one for the snake branch and the other for the remaining reptiles in the alignment. We then compared two models, a null model in which the snake $\omega$ was fixed at 1 and an alternative model in which it was free to vary. The likelihoods of these models were compared using a log-ratio test with one degree of freedom. Conservation in corn snakes was tested in a similar manner, with a third $\omega$ permitted for the corn snake branch and fixed at 1 under the null model. This model was compared to the earlier alternative model using a log-ratio test with two degrees freedom.

### Supplemental Figures

S1 PCA . . . 34
S2 Genetic diversity . . . 35
S3 Corn snake PRDM9 predicted binding motifs . . . 36
S4 Conservation of PRDM9 and ZCWPW1 . . . 37
S5 Decay of genome-wide LD . . . 38
S6 LDhelmet and Pyrho hotspots . . . 39
S7 Pyrho recombination map rate correlation with genomic features . . . 40
S8 LDhelmet recombination map characterization . . . 41
S9 Correlation of recombination rate with predicted PRDM9 binding sites of individual alleles . . . 42
S10 Zinc finger segments shared among PRDM9 alleles . . . 43
S11 Prairie rattlesnake PRDM9 predicted binding motifs . . . 44
S12 Overlap between predicted PRDM9 binding sites and CpG islands . . . 45
S13 Prairie rattlesnake recombination map rate correlation with genomic features . . . 46
S14 Genomic characteristics around CpG islands . . . 47
S15 Characteristics of inferred hotspots . . . 48
S16 Overlap of hotspots with predicted PRDM9 binding sites and promoter-like features . . . 49
S17 Predicted PRDM9 binding site losses and gains . . . 50
S18 Similarity between motifs enriched in hotspots and predicted PRDM9 binding motifs . . . 51
S19 GC* at hotspots and coldspots . . . 52
S20 Comparison of crossovers and LD-based recombination map . . . 53
S21 Overlap between crossovers (¡20 kb) and predicted binding sites and promoter-like features . . . 54
S22 Overlap of crossovers with features divided by sex of parent . . . 55
S23 Downsampling crossovers on macrochromosomes . . . 56
S24 Identification of scaffolds putatively belonging to the sex chromosomes . . . 57
S25 Removal of contaminated libraries . . . 58
S26 Mendelian errors per scaffold . . . 59
S27 Distribution of the fraction of methylated reads at CpG sites . . . 60

### Supplemental Tables

S1 Corn snake sample information . . . 61
S2 PRDM9 Allele Sequencing information . . . 62
S3 Phylogenetic Analysis by Maximum Likelihood . . . 63
S4 Repetitive elements . . . 64
S5 Scaffolds in the corn snake assembly likely belonging to the sex chromosomes (see section 1.6.1). 65
S6 NCBI Accession information for RNA-seq data used for genome annotation . . . 66
S7 Primer Sequences . . . 67

### Supplemental Figures

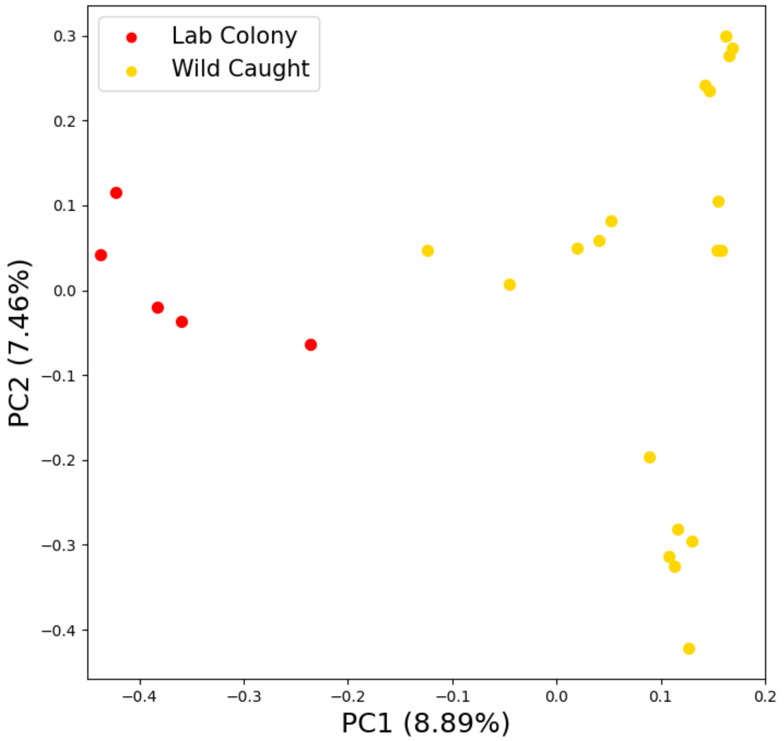

Figure S1: Principal component (PC) analysis of the "unrelated" corn snake samples. PCs were inferred from 583,817 SNPs (see Methods). Shown are PCs 1 and 2. The five samples in red are from a lab colony and the 19 samples in gold are wild caught.

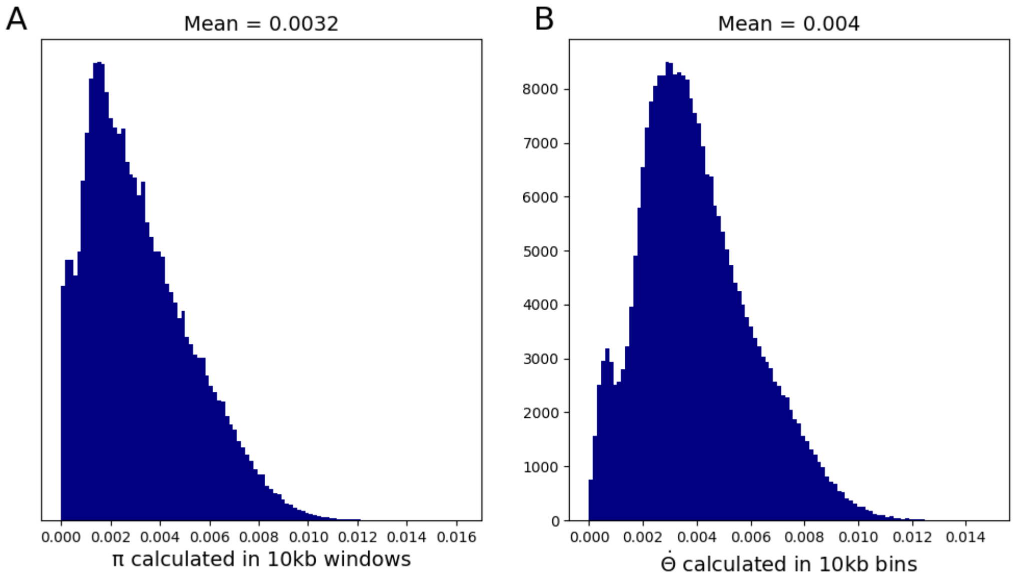

Figure S2: Genetic diversity in the "unrelated" corn snake samples. A) Heterozygosity ($\pi$) and B) Watterson estimator $\theta_w$, were calculated in 10 kb windows (see Methods).

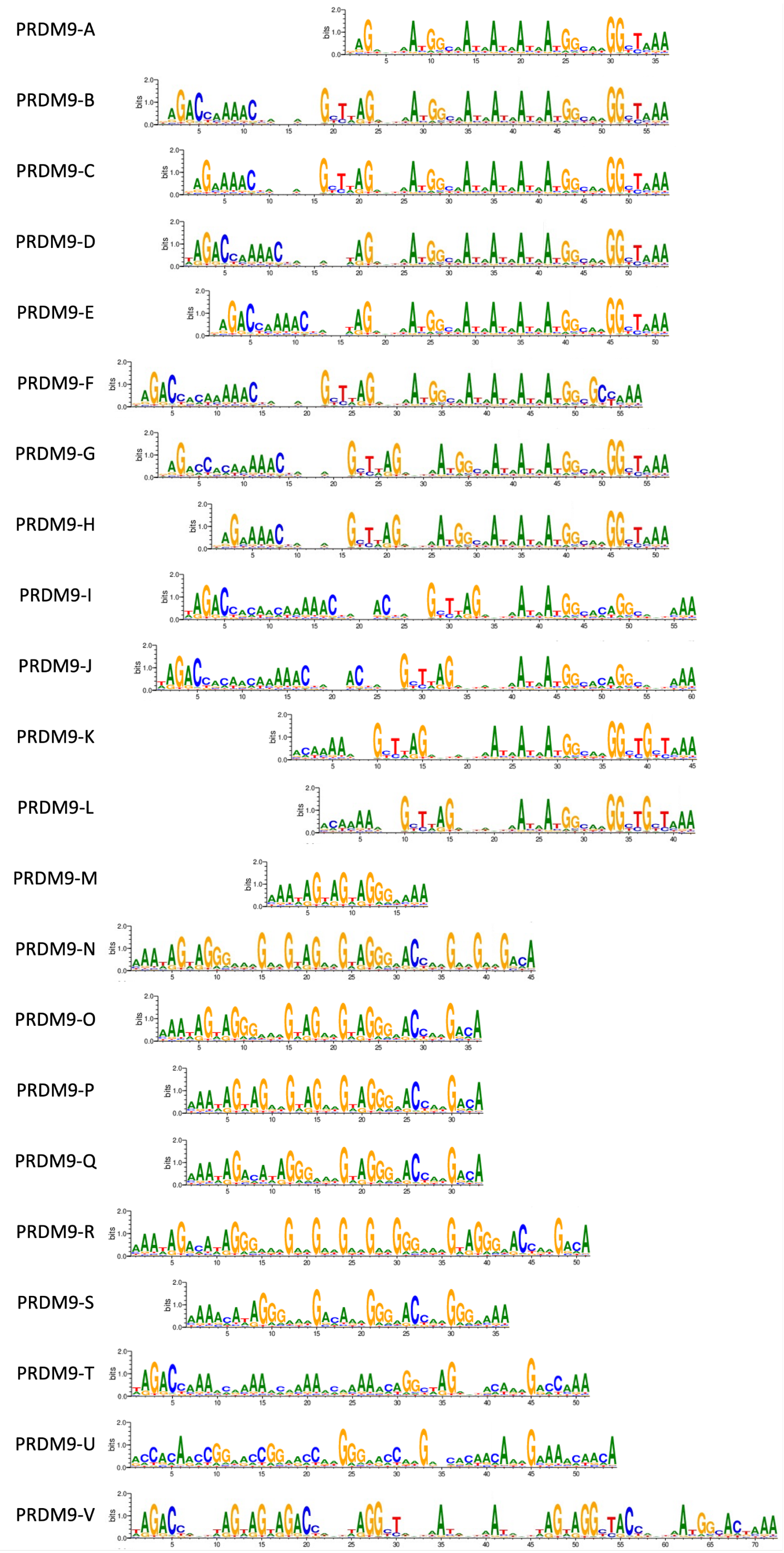

Figure S3: Position weight matrices representing predicted binding affinities for each of 22 PRDM9 alleles identified in corn snakes (Figure 1C).

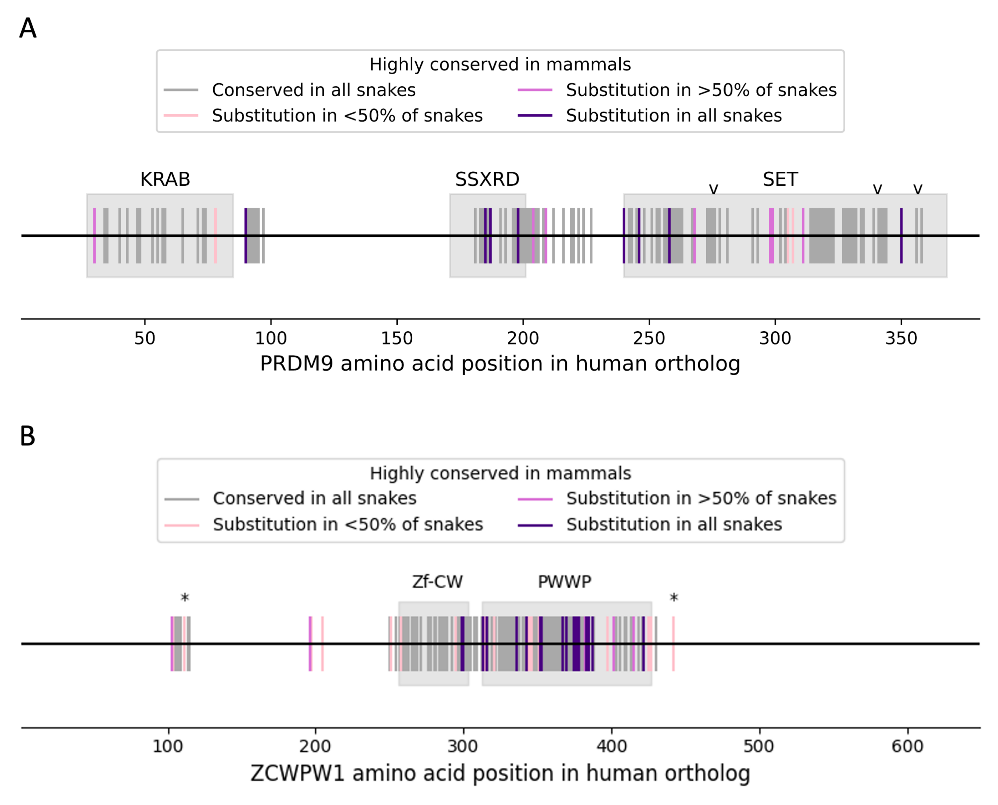

Figure S4: A) Conservation of PRDM9 amino acids between 135 mammals and 9 snakes, excluding the N-terminal coding exon (the zinc finger domain). There are no corn snake specific PRDM9 substitutions at residues that are highly conserved in mammals. The three catalytic tyrosine residues in the SET domain are annotated with a caret. B) Conservation of ZCWPW1 amino acids between 130 mammals and 7 snakes. Each tick mark denotes a site at which >95% mammals carry the same amino acid. Amino acids shown as gray ticks are completely conserved in the nine snakes; light pink are conserved in all but 1-4 snakes, dark pink are changed in 5-8 snakes, and dark purple amino acids are changed in all 9 snakes. Asterisks denote sites at which corn snakes are the only snakes with a substitution.

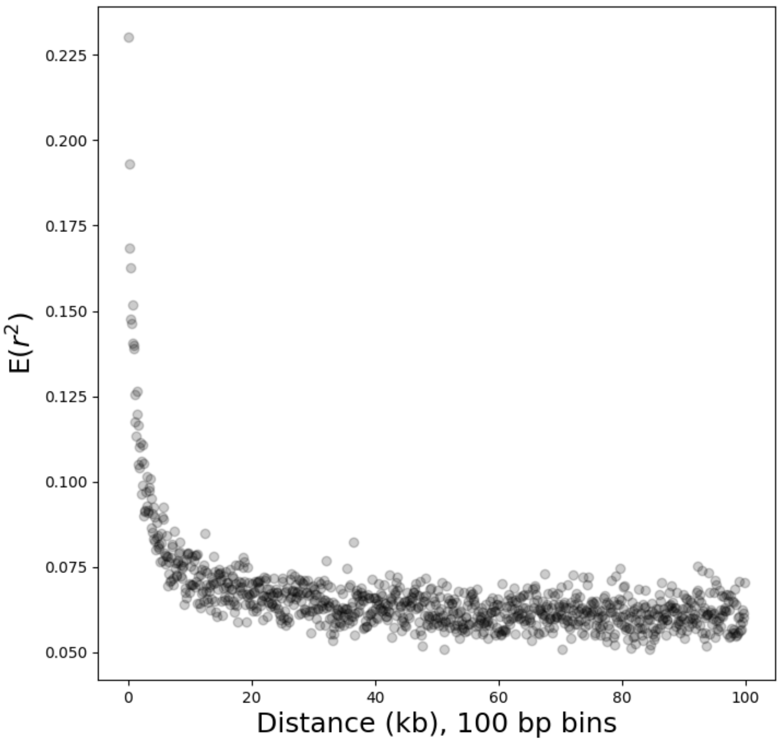

Figure S5: Decay of pairwise linkage disequilibrium (expected r2) for 1% of SNPs with minor allele frequency >5% across the genome, chosen at random. Points represent mean r2 between SNPs at a given distance apart, in bins of 100 bp.

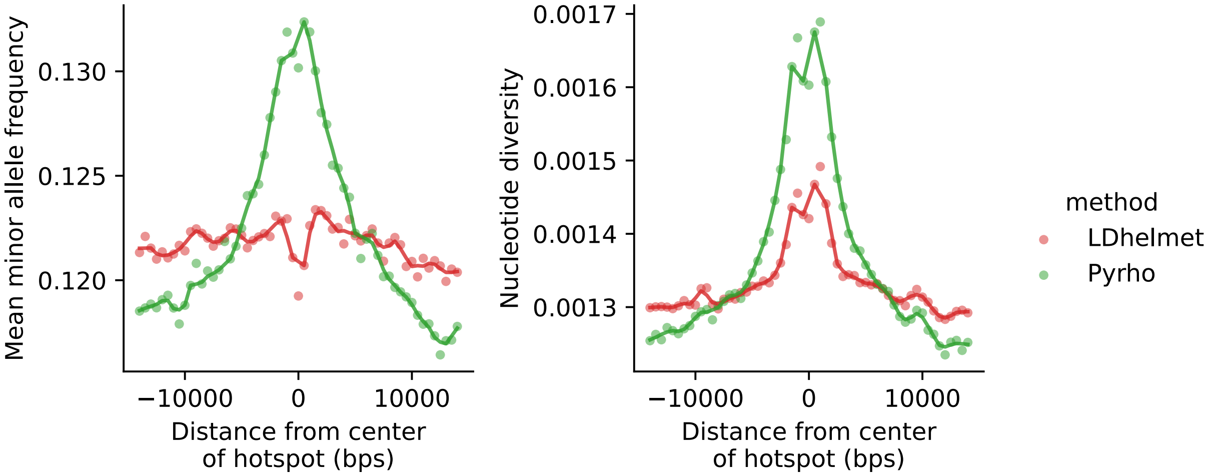

Figure S6: Allele frequencies and diversity levels around hotspots inferred by Pyrho or LDhelmet. Both programs infer the population recombination rate, $\rho$, from linkage disequilibrium patterns. The mean minor allele frequency and nucleotide diversity were calculated in sliding windows of 1 kb with an offset of 500 bps in the regions surrounding (15 kb added to each end) hotspots called from Pyrho (green) or LDhelmet (red). These statistics suggest that rho estimates from Pyrho may be inflated where the local gene genealogy is deep.

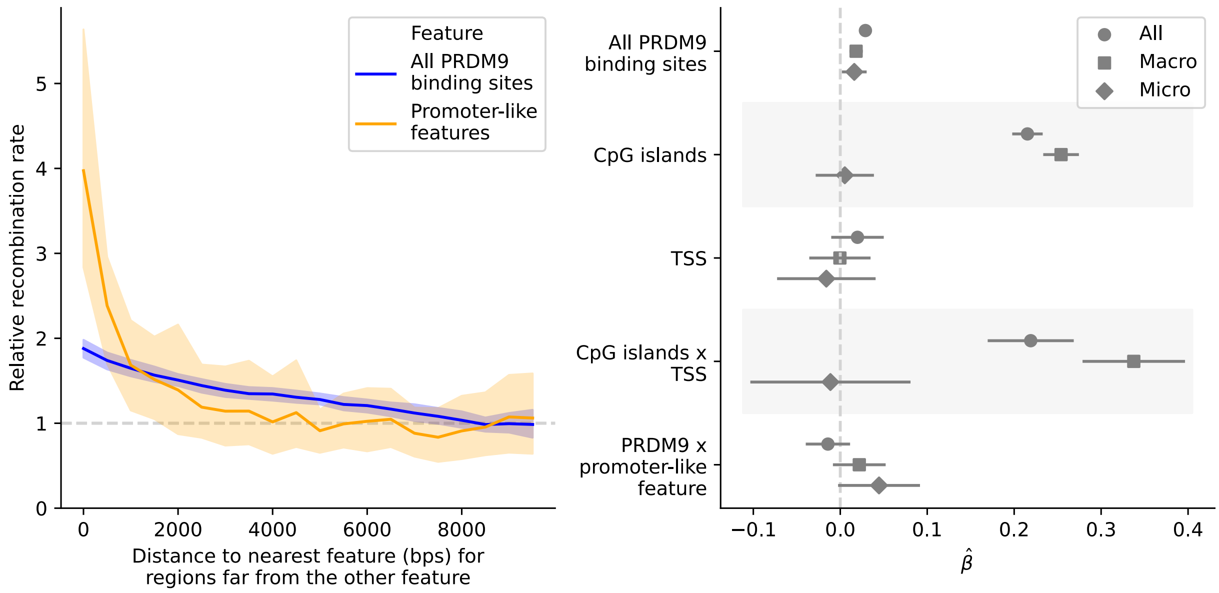

Figure S7: Qualitative conclusions remain the same when using Pyrho to infer rates instead of LDhelmet: an increase in the population recombination rate is seen around both PRDM9 binding sites and CpG islands. (Left panel) As in Figure 2A: Mean population recombination rate in 100 bp windows as a function of distance to the nearest predicted PRDM9 binding site (blue) or promoter-like feature (orange) (plotted for bins of 500 bp). When considering one feature, we condition on windows >10 kb from the other feature; thus, when focusing on predicted PRDM9 binding sites, we only consider windows that are ¿10 kb from promoter-like features (i.e., TSS or CpG island). The recombination rate is relative to the mean rate 8-10 kb away. Shaded regions represent the central 95% confidence interval obtained by bootstrapping (see Methods). (Right panel) As in Figure 2C: Point estimate and 95% CI for the coefficients of a linear model, in which the response variable is the (log) recombination rate in 1 kb windows (thinned to be 10 kb apart) and the predictors are the (binary) presence or absence of one or more predicted PRDM9 binding sites, TSS, or CpG islands. Covariates include the background recombination rate (1Mb scale) and GC content (see Methods). Results are reported for data from the autosomes (circles), only scaffolds assigned to macrochromosomes (squares), and only microchromosomes (diamonds).

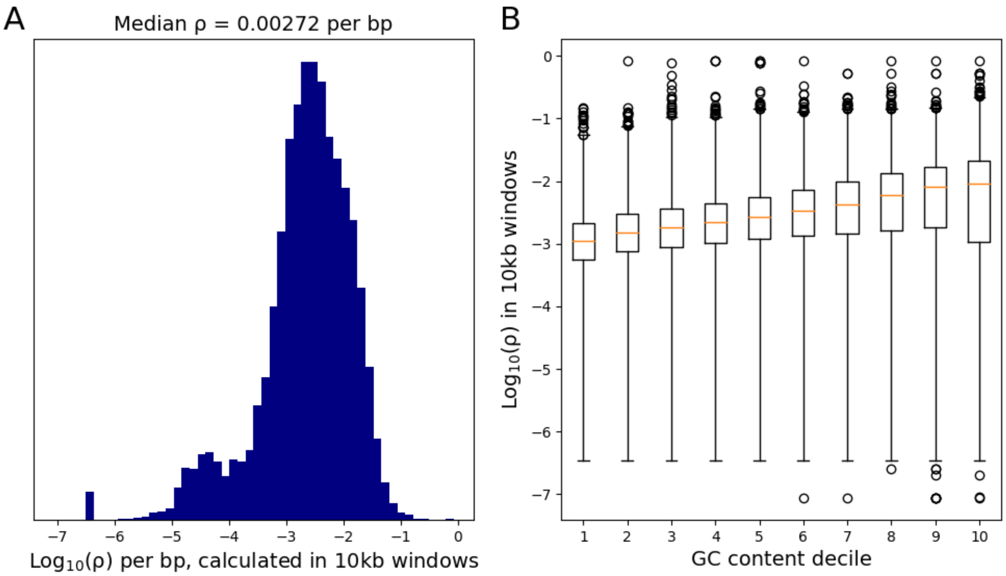

Figure S8: A) Mean recombination rate per bp on a log scale, calculated in 10 kb windows. B) Correlation between recombination rate and GC content. 10kb windows were divided into deciles based on GC content. Box and whiskers are plotted for each decile, where the whiskers exclude only the most extreme 0.1% of the data (circles) and the orange horizontal line represents the median for a given GC content decile.

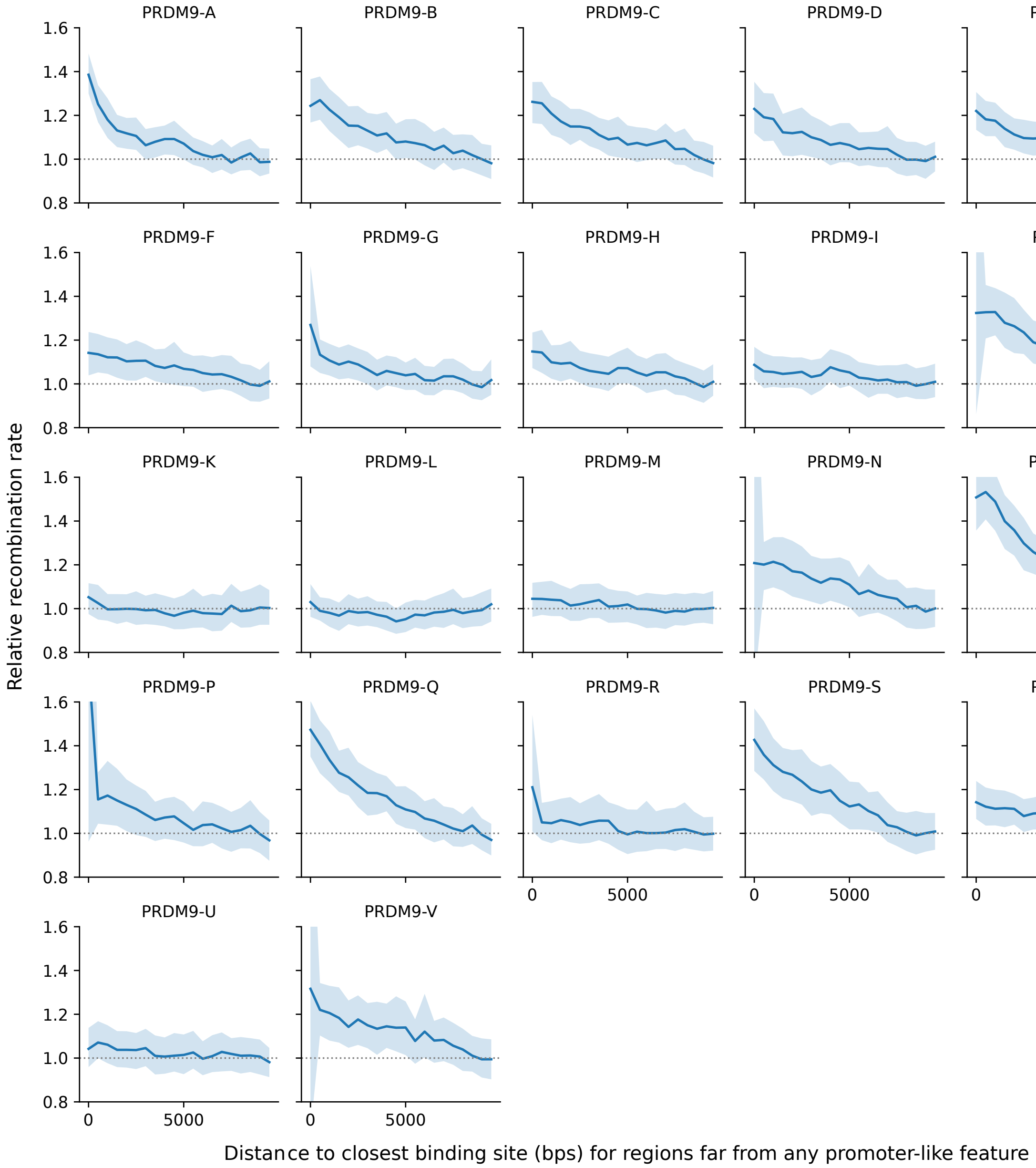

Figure S9: Population recombination rate around predicted PRDM9 binding sites for individual PRDM9 alleles. Solid lines represent the mean population recombination rate in 100 bp windows as a function of distance to the nearest predicted binding site (plotted for bins of 500 bp), for windows that are >10 kb from promoter-like features (i.e., a TSS or CpG island). The recombination rate is relative to the mean rate 8-10 kb away. Shaded regions represent the central 95% confidence interval obtained by bootstrapping

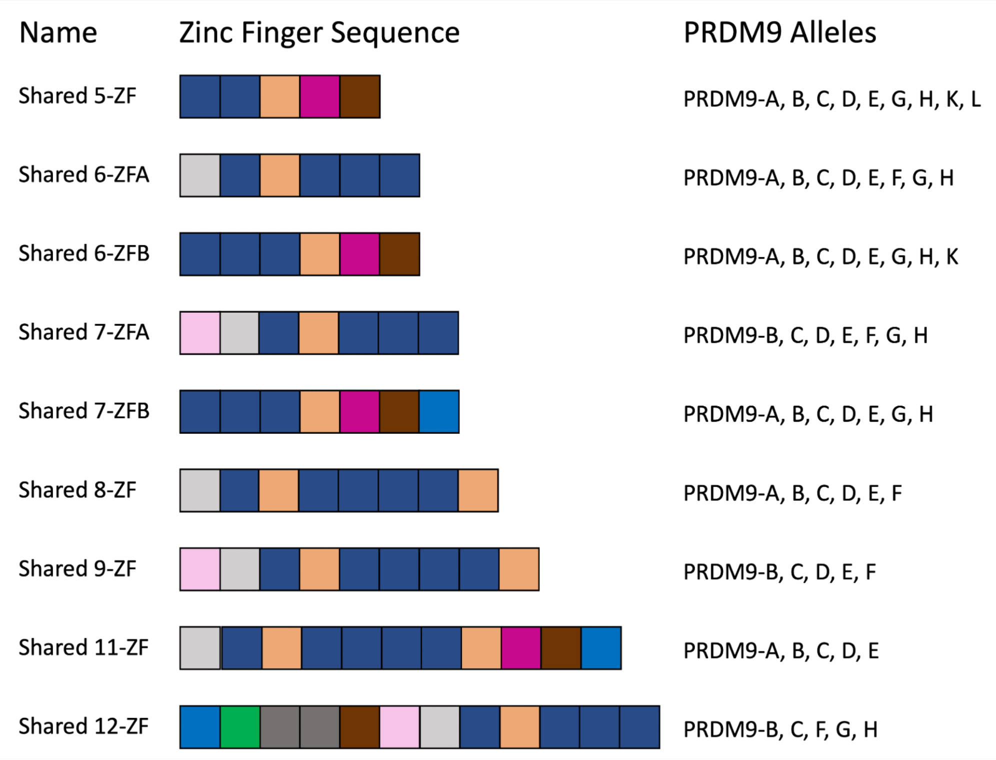

Figure S10: Representation of all sets of five or more consecutive fingers observed in at least five alleles. The color scheme of the zinc fingers is the same as in Figure 1C.

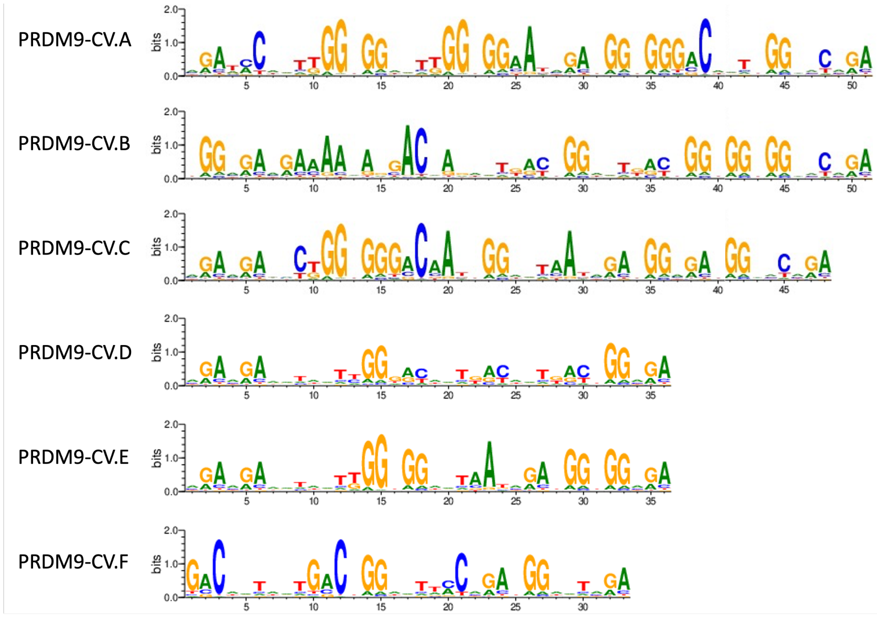

Figure S11: Position weight matrices representing predicted binding affinities for each of six PRDM9 alleles identified in three prairie rattlesnake individuals (see Methods).

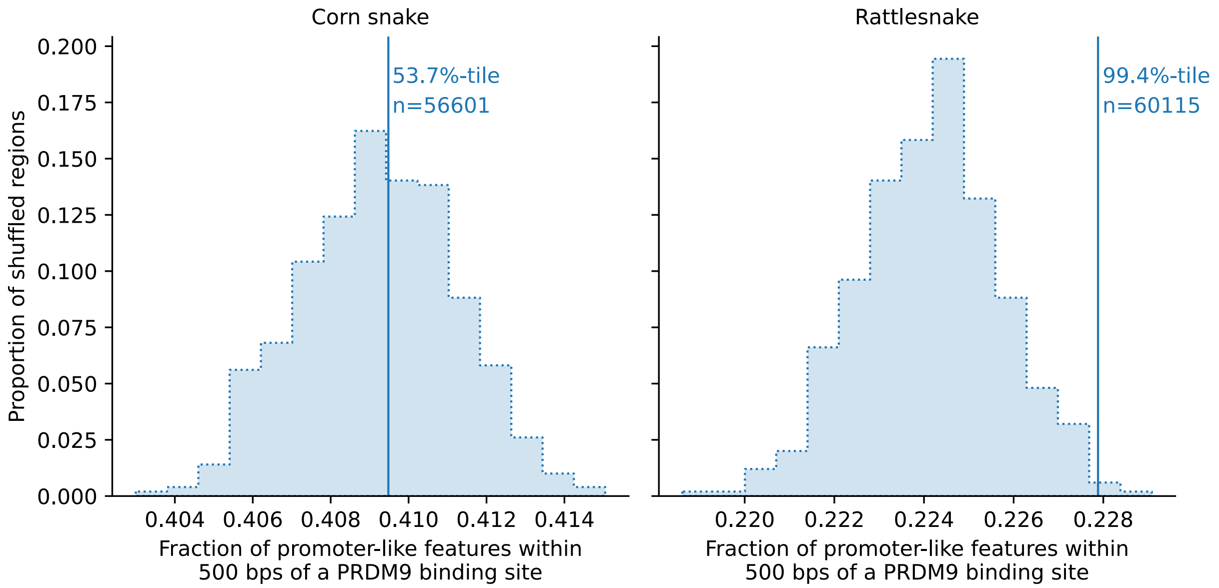

Figure S12: Overlap between predicted PRDM9 binding sites and promoter-like features. The observed overlap between predicted PRDM9 binding sites and promoter-like features (either CpG islands and/or TSS) are shown with vertical lines for corn snakes (left) and rattlesnakes (right). The overlap expected by chance is shown with the shaded distribution and is based on 500 replicates, in which each region was placed at random within the same scaffold of the original location, conditional on there not being a gap in the genome sequence.

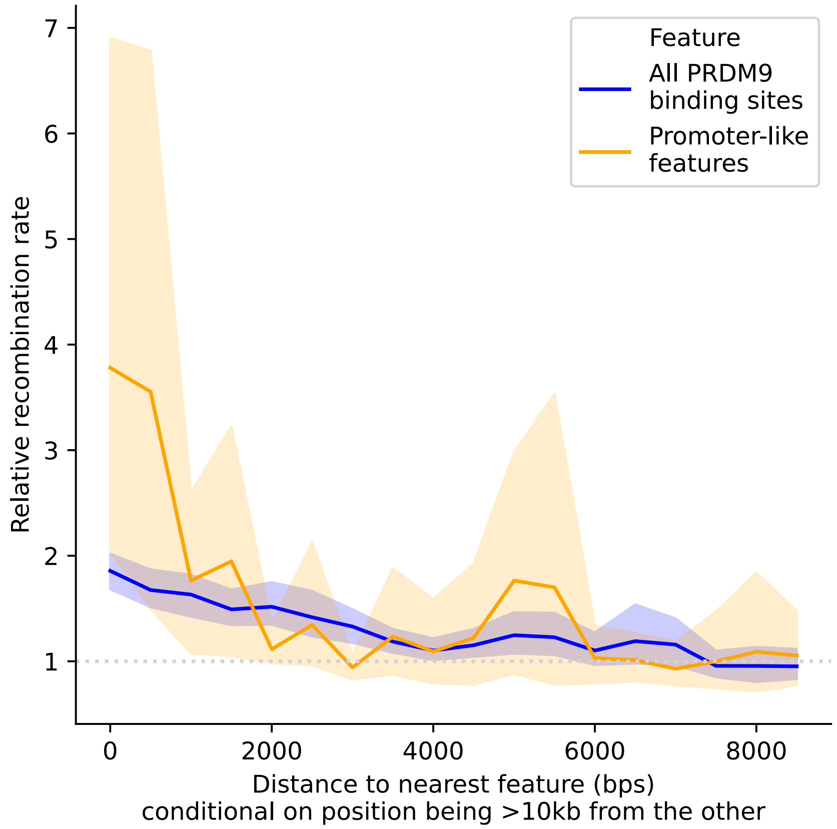

Figure S13: In prairie rattlesnake, the mean population recombination rate as a function of distance to the nearest predicted PRDM9 binding site (blue) or promoter-like feature (orange) (in 100 bp windows, plotted in bins of 500 bp). When considering one feature, we condition on the windows laying >10 kb from the other feature; thus, when focusing on predicted PRDM9 binding sites, we only consider windows that are >10 kb from promoter-like features (i.e., TSS or CpG island). The recombination rate is relative to the mean rate 8-10 kb away. Shaded regions represent the central 95% confidence interval obtained by bootstrapping.

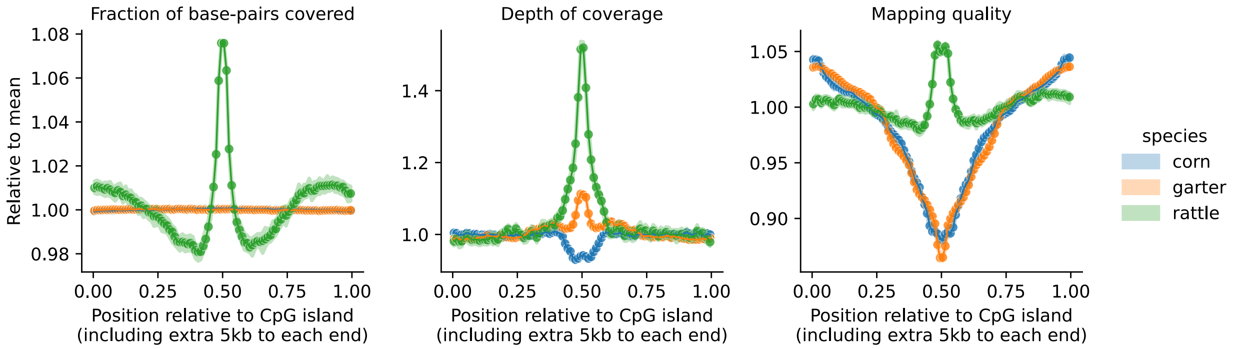

Figure S14: Various measures around CpG islands in corn snakes, garter snakes and rattlesnakes. CpG islands were identified in each assembly using `cpgplot` with default parameters (see section 10.3). Using alignments of the reference individual to its corresponding assembly, we measured the fraction of positions covered by at least one read (right), the mean depth of coverage (center), and mean mapping quality (left) around CpG islands plus an extra 5kb to each end, using sliding windows of 100 bps with an offset of 50 bps. Note that because the length of CpG islands is not constant, we plot the x-axis as the relative position of windows within the whole interval of the CpG island and the 5kb padding (i.e., 0.5 represent the middle point of the interval, the center of the CpG island). The metrics are normalized by the mean across all windows within a species (y axis). Together, these quality measures suggest that the rattlesnake genome may be less reliably assembled around CpG islands, perhaps because reads belonging to paralogous locations were sometimes collapsed.

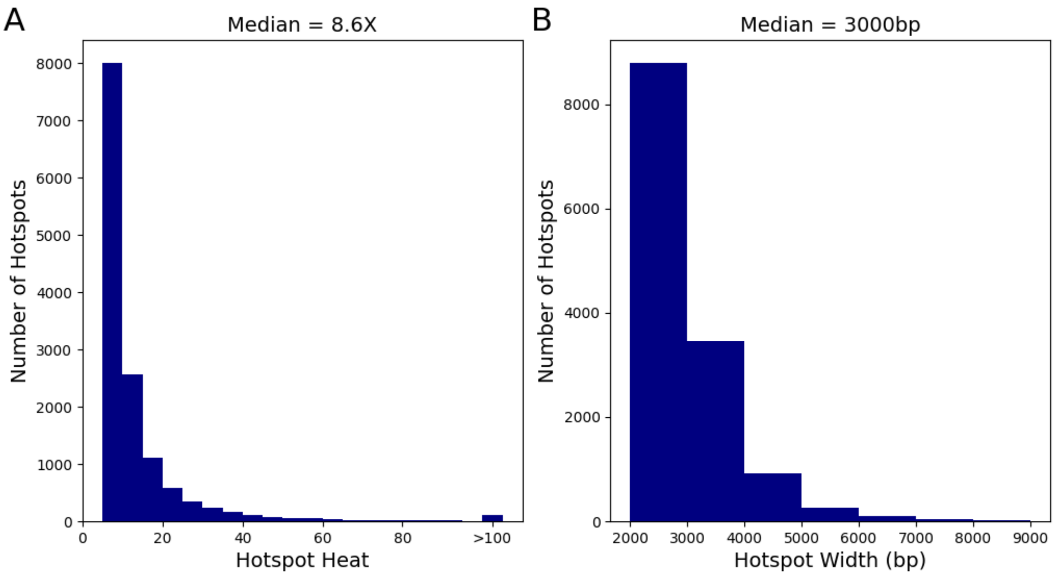

Figure S15: A) A histogram of estimated fold-change (heat) of hotspots; the ratio of the recombination rate within the hotspot to the recombination rate in the surrounding 40 kb (see Methods). B) A histogram of estimated hotspot widths. Hotspots (i.e., 2 kb windows with >5-fold heats) were merged if they were within 5 kb of each other; what is reported is the width of the resulting, merged hotspot.

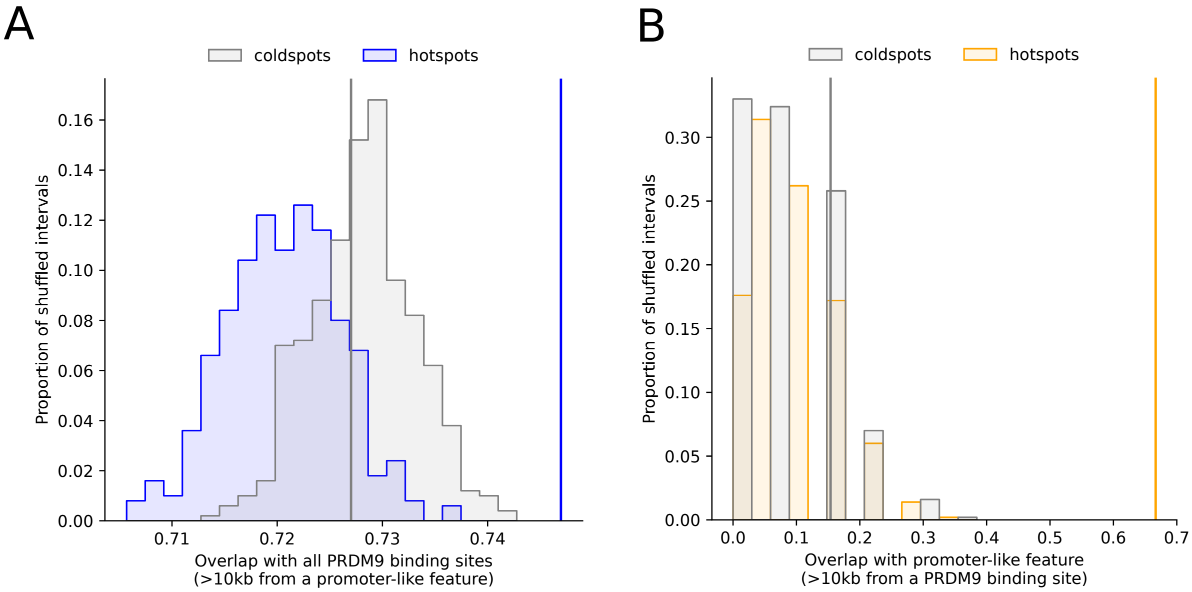

Figure S16: A) Overlap of hotspots (blue) and matched coldspots (gray) with the predicted binding sites for the all PRDM9 alleles, conditional on them being far from a promoter-like feature. The observed values are shown with solid lines. The overlap expected by chance is shown with the shaded distribution and is based on 500 replicates, in which each hotspot was placed at random within 5 Mb of the original location, conditional on there not being a gap in the genome sequence (see Methods). We note that while hotspots and coldspots are matched for base composition (see Methods), that need no longer be the case once we condition on them laying far PRDM9 binding sites or promoter-like features, driving the slight difference in null distributions. B) As in A, but for the overlap of hotspots (orange) and matched coldspots (gray) with promoter-like features far from any PRDM9 binding site.

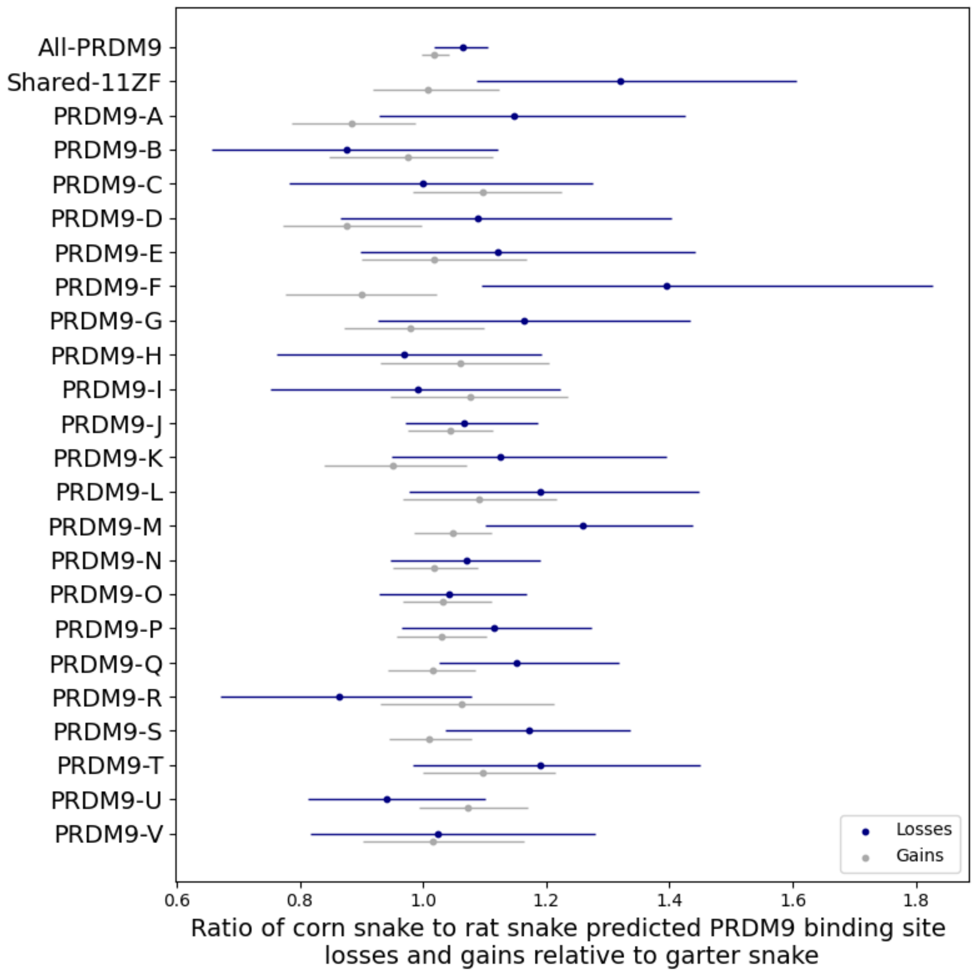

Figure S17: The ratio of losses in the corn snake lineage relative to the black rat snake (in blue) and the ratio of gains in the corn snake lineage relative to the black rat snake (in gray), for the predicted binding sites of all corn snake PRDM9 alleles together ("All-PRDM9") and for each allele individually. The 95% confidence intervals were obtained by bootstrapping over aligned regions of variable length that contain no repeats, insertions, or deletions, and which had at least one gain or loss event in either of the two lineages.

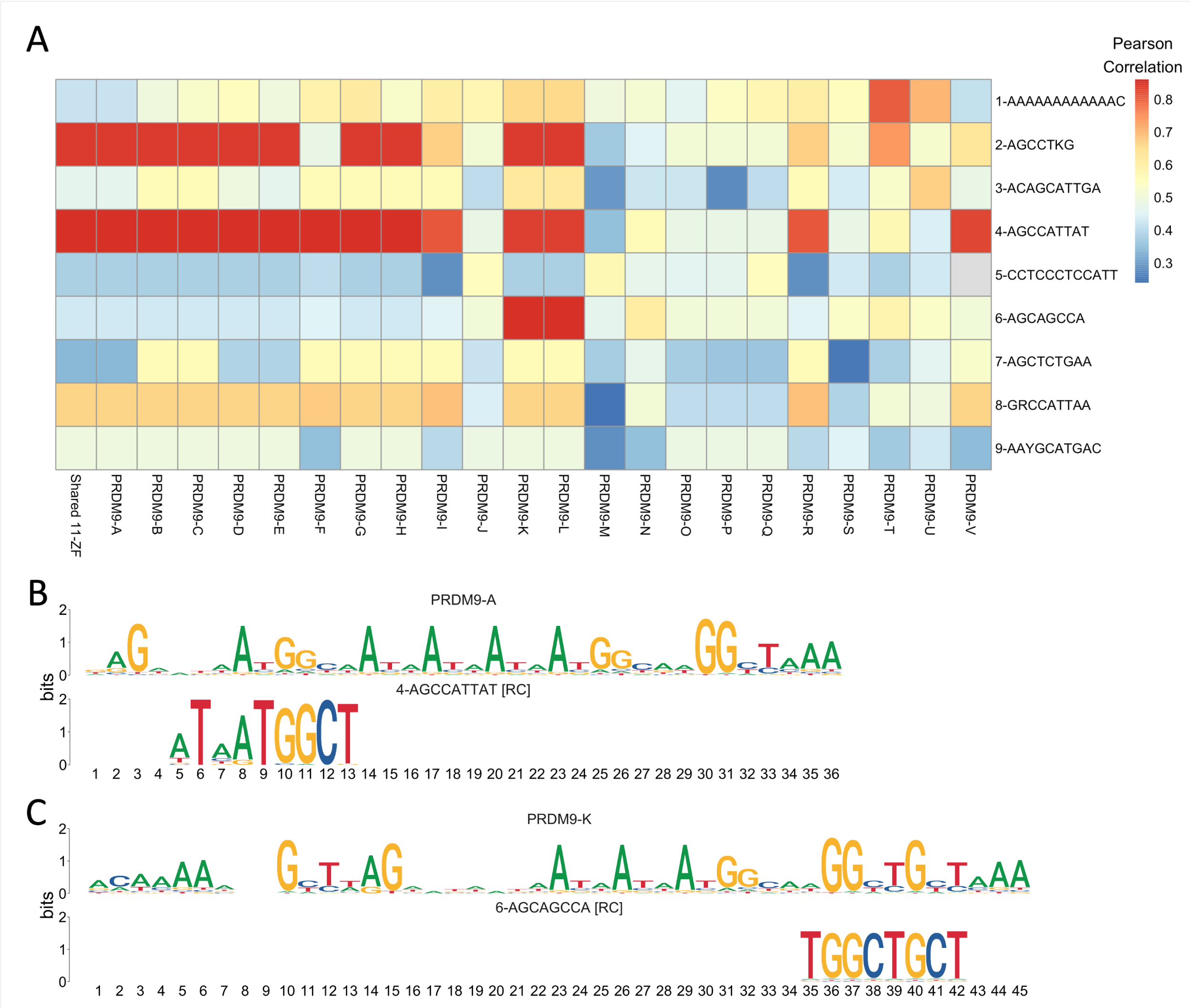

Figure S18: Motifs enriched in hotspots relative to coldspots are similar to predicted PRDM9 binding motifs. A) The color corresponds to the Pearson correlation coefficient associated with the best alignment between each motif (X-axis) and allele (Y-axis). Gray boxes represent positions where comparisons that failed due to low motif position information content (See Methods). B-C) Shown are the best alignments between two motifs enriched in hotspots (motifs 4 and 6) and the PRDM9 alleles with which they have the highest similarity (Pearson correlation coefficient ¿.85), alleles PRDM9-A and PRDM9-K, respectively.

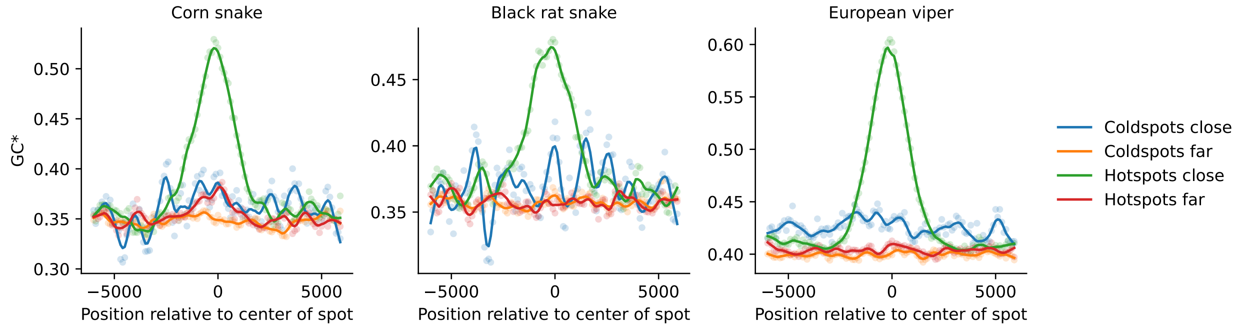

Figure S19: Flux to GC (GC*) as a function of distance from corn snake autosomal hotspots and coldspots, in sliding windows of 500 bp with a 100 bp offset, for the lineages leading to corn snake (left), black rat snake (center), and European viper (right). Note that rattlesnakes were included as an outgroup only (see Methods). GC* around hotspots (or coldspots) that are close (¡500 bp) to a promoter-like feature and far (>10 kb) from any promoter-like feature are shown. Local regression curves are shown for a span of 0.05.

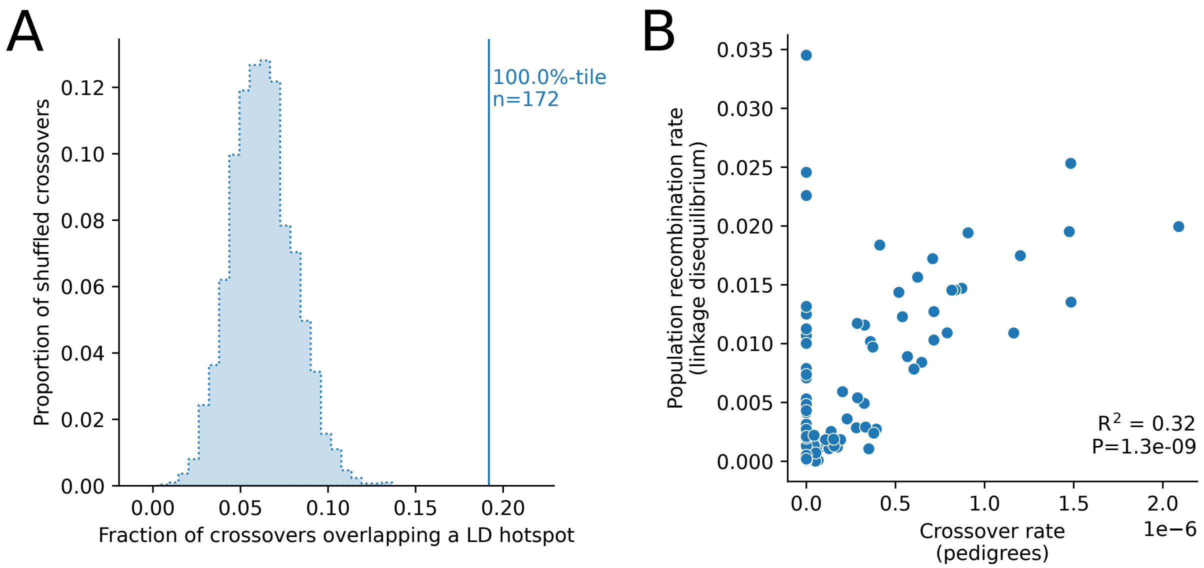

Figure S20: Comparison of LD-based recombination map and rate estimates based on crossovers identified in two pedigrees. A) Overlap of crossovers with LD hotspots. The solid line is the observed overlap for crossovers. The frequency distribution represents the overlap for 3000 sets of simulated crossovers obtained by placing the observed interval lengths down at random within 5 Mb of the original crossover interval, conditional on it containing at least two informative markers and there not being a gap in the genome sequence at that location (see section 9.2). B) Correlation between the mean population recombination rate per scaffold as estimated from LDHelmet and the crossover rate in two pedigrees, for scaffolds over 100 Kb. Conditioning on scaffolds with at least one crossover observed gets rid of the smear at 0, and greatly increases the variance explained (to $R^2$=0.67).

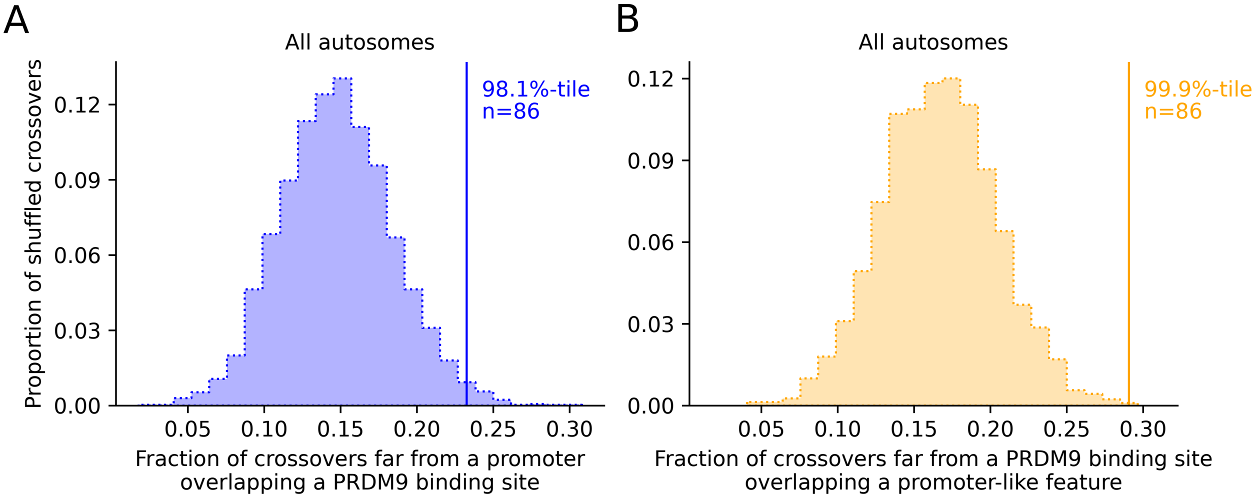

Figure S21: Crossovers identified from pedigree resequencing with a resolution of ¡20 kb overlap both predicted PRDM9 binding sites and promoter-like features significantly more often than expected by chance in corn snakes. A) Overlap of crossovers with the predicted binding sites of the PRDM9 alleles carried by the parent in which the crossover is inferred to have occurred, for binding sites far from a promoter-like feature (>10 kb). B) Overlap of crossovers with promoter-like features far (>10 kb) from the PRDM9 binding sites of alleles carried by the parents. The solid line is the observed overlap for the n crossovers that satisfy the criteria. The frequency distribution represents the overlap for 3000 sets of simulated crossovers obtained by placing the observed interval lengths down at random within 5 Mb of the original crossover interval, conditional on it containing at least two informative markers and there not being a gap in the genome sequence at that location (see section 9.2). The rank of the observation relative to realizations under the null is given as a percentile[82].

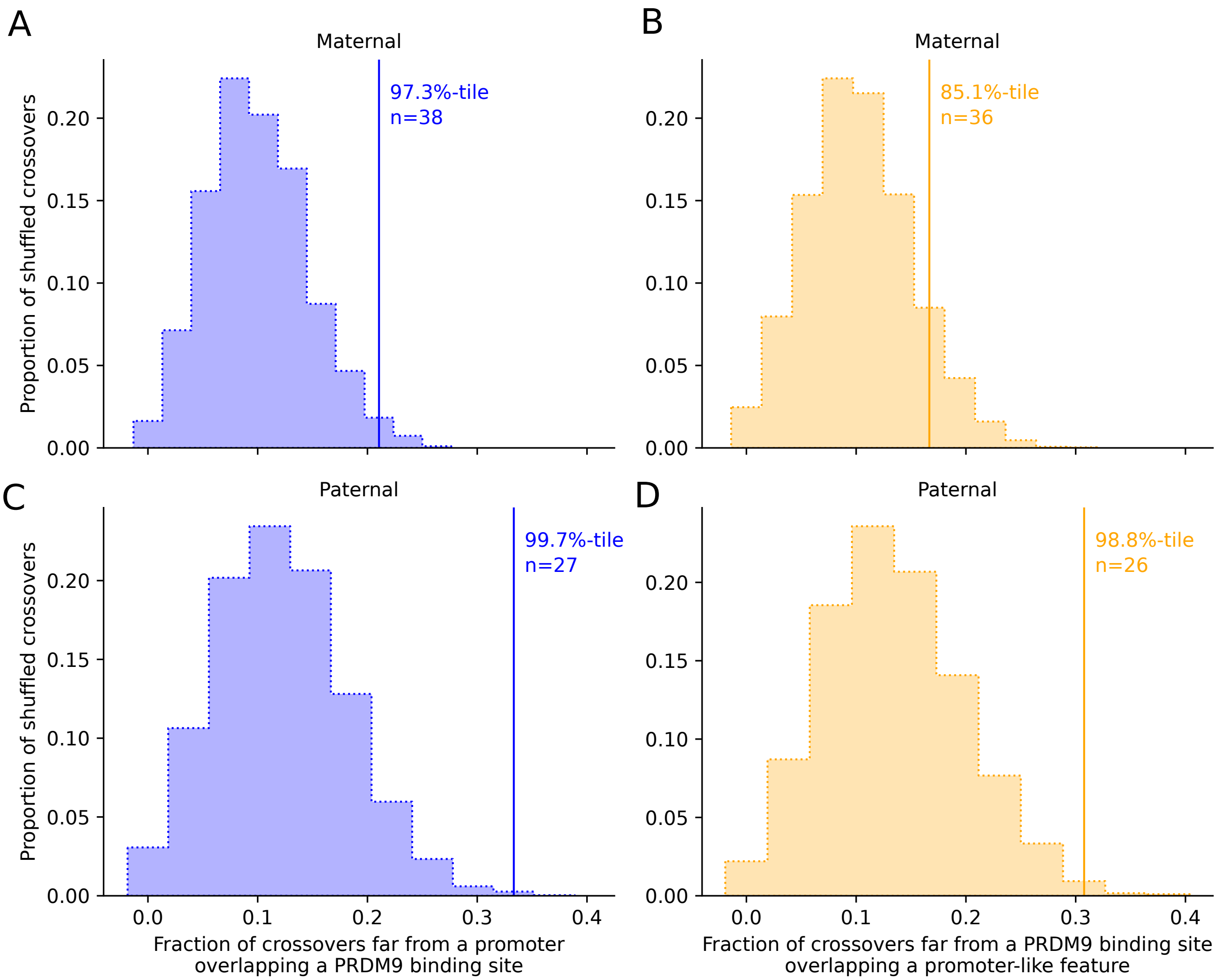

Figure S22: A) Overlap of crossovers with the predicted binding sites of the PRDM9 alleles carried by the mother in which the crossover is inferred to have occurred, for binding sites far from a promoter-like feature (>10 kb). The solid line is the observed overlap for the n crossovers that satisfy the criteria. The frequency distribution represents the overlap for 3000 sets of simulated crossovers obtained by placing the observed interval lengths down at random within 5 Mb of the original crossover interval, conditional on it containing at least two informative markers and there not being a gap in the genome sequence at that location (see section 9.2). The rank of the observation relative to realizations under the null is given as a percentile (North, Curtis, and Sham 2002). B) As in panel A, but for the overlap of crossovers with promoter-like features far (>10 kb) from the PRDM9 binding sites of alleles carried by the mothers. (C) and (D) As in panels A and B, but for crossovers that occurred in fathers.

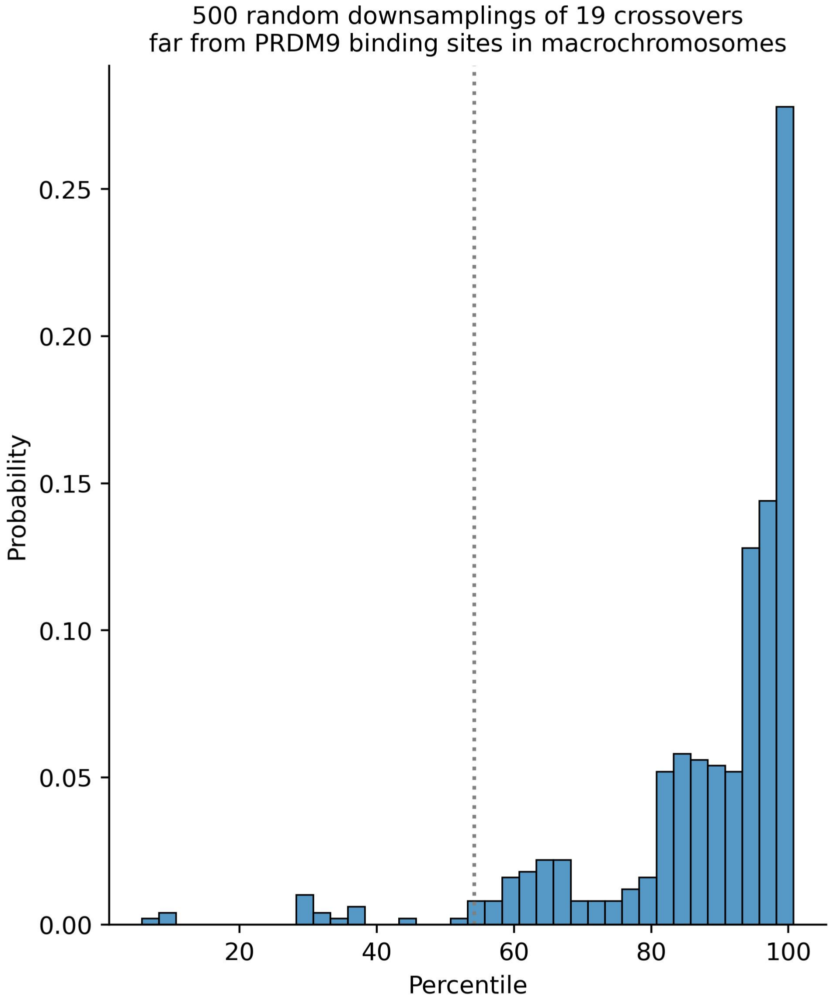

Figure S23: The number of crossovers in macrochromosomes far (>10kb away) from any PRDM9 binding site (n=43) was randomly downsampled 500 times, to match the number of crossovers far from any PRDM9 binding site found in microchromosomes (n=19). For each subsample, we calculated the percentile of the observed overlap in a null distribution of overlap obtained by shuffling crossover locations 3,000 times (see section 9.2). The distribution of percentiles across the 500 subsamples is shown as an histogram; the percentile observed for microchromosomes is shown as a dotted line. Thus, the difference between macrochromosomes and microchromosomes is unlikely to be due to sampling error.

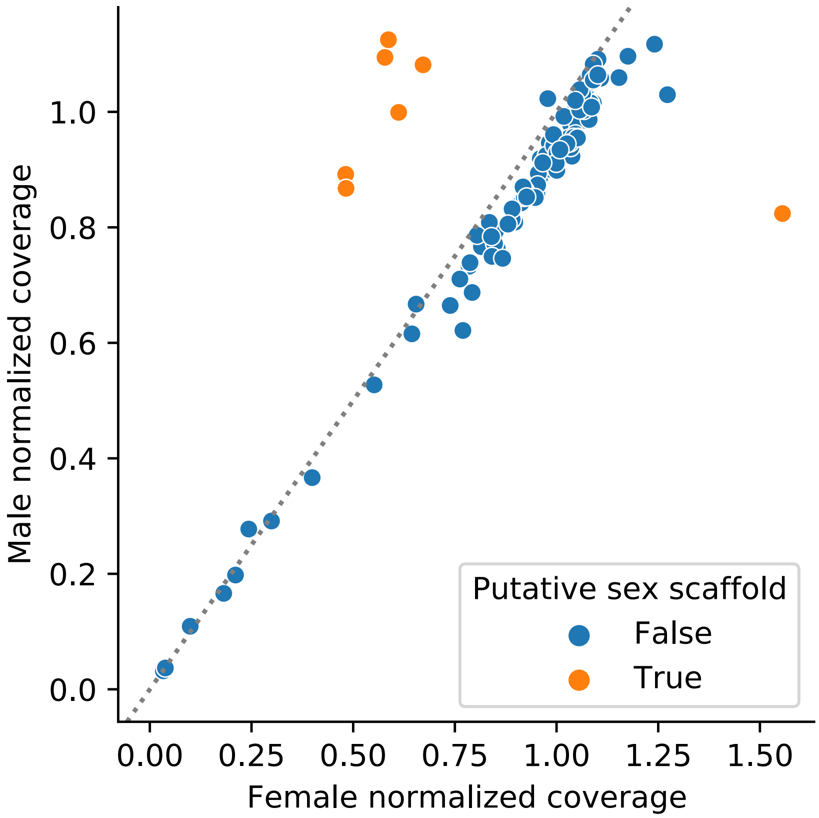

Figure S24: Identification of scaffolds putatively belonging to the sex chromosomes. We calculated the mean coverage per scaffold in the parents in the pedigrees, for which we know the sex. We classified scaffolds as non-autosomal (shown in orange) if the female-to-male depth of coverage ratio was $<0.7$ (putatively belonging to the Z) or $>1.3$ (putatively belonging to the W). Note that the values shown in the plot are normalized by the sex-specific mean depth of coverage across all scaffolds.

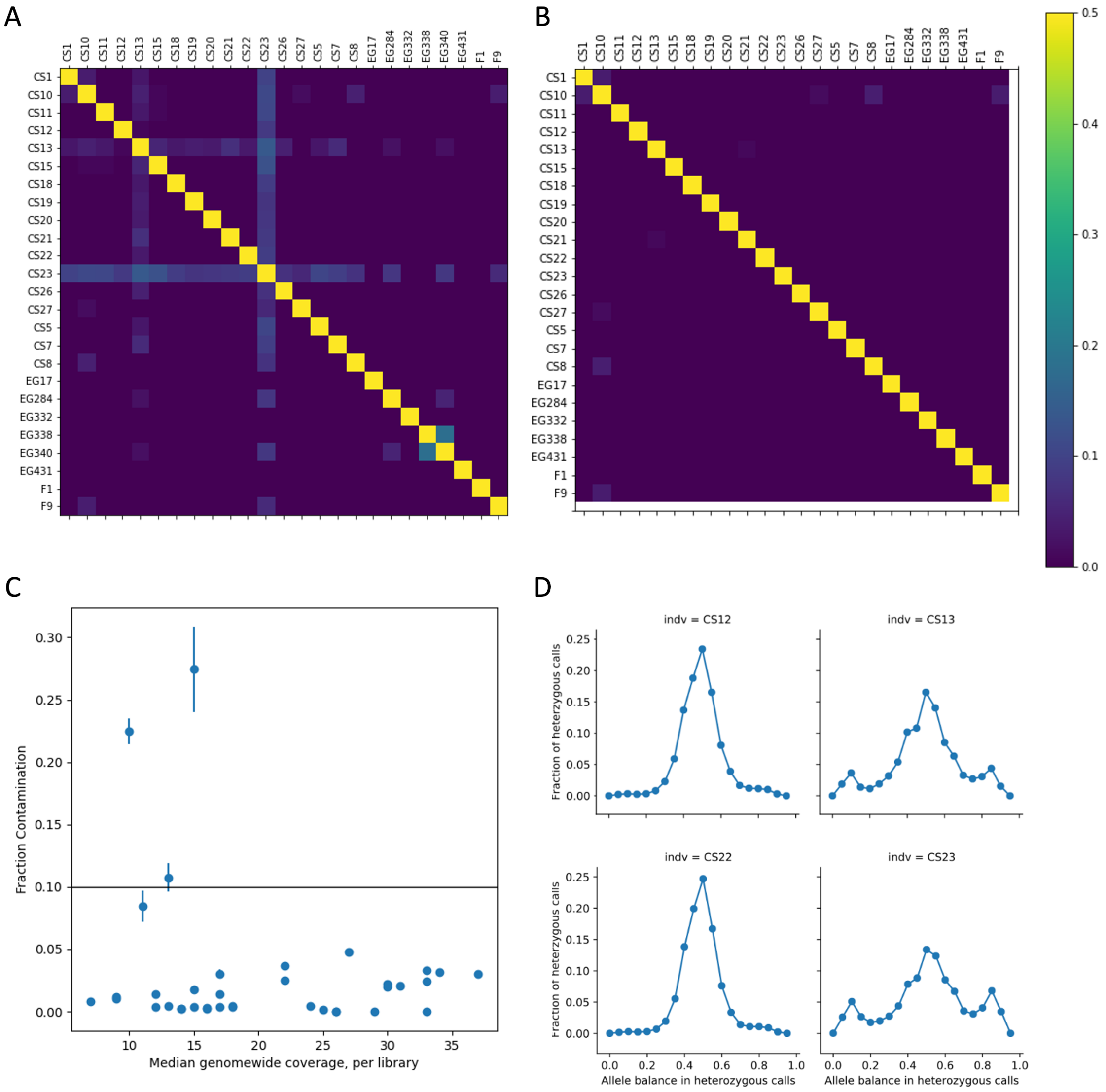

g

Figure S25: Identification and removal of contaminated libraries - A) After an initial round of variant calling, two samples showed unexpected relatedness (cite) to many other samples - samples CS13 and CS23. B) Exclusion of libraries with high estimated levels of contamination removed the signal of unexpected relatedness. C) Fraction of contamination per library (estimated by GATK CalculateContamination) after initial round of variant calling, as estimated by GATK CalculateContamination, verses per library genome-wide coverage (Picard CollectWgsMetrics). Libraries with more than 10% contamination were removed from subsequent analyses, including libraries for CS13 and CS23. D) Examples of the distribution of allelic balance in heterozygous calls prior to contaminated library removal, for two samples with very low levels of estimated contamination (CS12, CS22) and the two samples with high estimated levels of contamination (CS13, CS23).

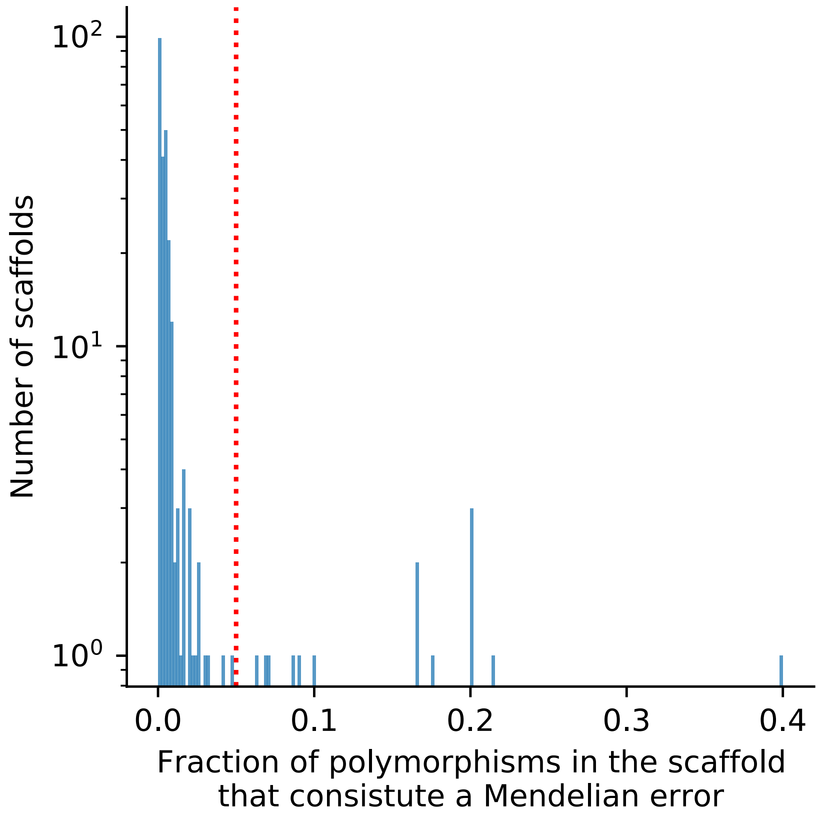

Figure S26: Fraction of polymorphisms in each scaffold that constitute a Mendelian error. Scaffolds with an unusually high rate of Mendelian errors (>5%, red line) were discarded (<.01% of the total sequence).

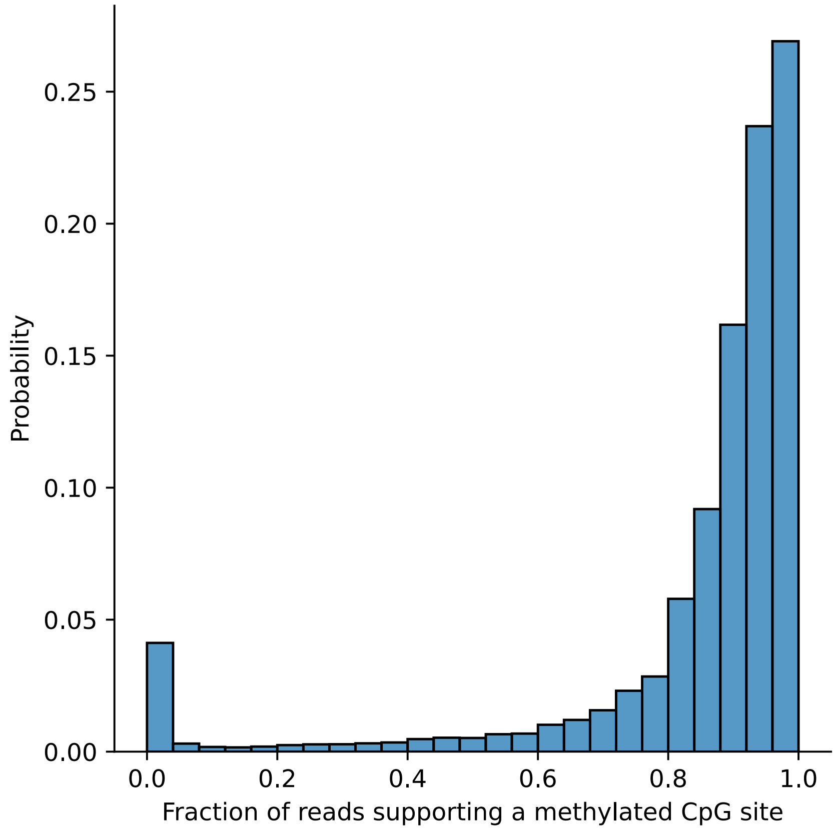

Figure S27: Distribution of the fraction of bisulfite sequencing reads that support a methylated cytosine across CpG sites in the corn snake genome (see section 1.5.2).

### References

[1] A. Ullate-Agote, I. Burgelin, A. Debry, C. Langrez, F. Montange, R. Peraldi, J. Daraspe, H. Kaessmann, M. C. Milinkovitch, and A. C. Tzika. Genome mapping of a lyst mutation in corn snakes indicates that vertebrate chromatophore vesicles are lysosome-related organelles. *Proceedings of the National Academy of Sciences*, 117(42):26307–26317, 2020.

[2] Asier Ullate-Agote, Michel C Milinkovitch, and Athanasia C Tzika. The genome sequence of the corn snake (pantherophis guttatus), a valuable resource for evodevo studies in squamates. *International Journal of Developmental Biology*, 58(10-11-12):881–888, 2015.

[3] S. Koren, B. P. Walenz, K. Berlin, J. R. Miller, N. H. Bergman, and A. M. Phillippy. Canu: scalable and accurate long-read assembly via adaptive k-mer weighting and repeat separation. *Genome research*, 27(5):722–736, 2017.

[4] M. J. Roach, S. A. Schmidt, and A. R. Borneman. Purge haplotigs: allelic contig reassignment for third-gen diploid genome assemblies. *BMC bioinformatics*, 19(1):1–10, 2018.

[5] Heng Li. New strategies to improve minimap2 alignment accuracy. *Bioinformatics*, 37(23):4572–4574, 2021.

[6] Mao Qin, Shigang Wu, Alun Li, Fengli Zhao, Hu Feng, Lulu Ding, and Jue Ruan. Lrscaf: improving draft genomes using long noisy reads. *BMC genomics*, 20(1):1–12, 2019.

[7] Mengyang Xu, Lidong Guo, Shengqiang Gu, Ou Wang, Rui Zhang, Brock A Peters, Guangyi Fan, Xin Liu, Xun Xu, Li Deng, et al. Tgs-gapcloser: a fast and accurate gap closer for large genomes with low coverage of error-prone long reads. *GigaScience*, 9(9):giaa094, 2020.

[8] Mahul Chakraborty, James G Baldwin-Brown, Anthony D Long, and JJ Emerson. Contiguous and accurate de novo assembly of metazoan genomes with modest long read coverage. *Nucleic acids research*, 44(19):e147–e147, 2016.

[9] Jennifer M Shelton, Michelle C Coleman, Nic Herndon, Nanyan Lu, Ernest T Lam, Thomas Anantharaman, Palak Sheth, and Susan J Brown. Tools and pipelines for bionano data: molecule assembly pipeline and fasta super scaffolding tool. *BMC genomics*, 16(1):1–16, 2015.

[10] Daren C Card, Richard H Adams, Drew R Schield, Blair W Perry, Andrew B Corbin, Giulia IM Pasquesi, Kristopher Row, Melissa J Van Kleeck, Juan M Daza, Warren Booth, et al. Genomic basis of convergent island phenotypes in boa constrictors. *Genome biology and evolution*, 11(11):3123–3143, 2019.

[11] Jullien M Flynn, Robert Hubley, Clément Goubert, Jeb Rosen, Andrew G Clark, Cédric Feschotte, and Arian F Smit. Repeatmodeler2 for automated genomic discovery of transposable element families. *Proceedings of the National Academy of Sciences*, 117(17):9451–9457, 2020.

[12] A.F.A. Smit, R. Hubley, and P. Green. Repeatmasker open-4.0. `http://www.repeatmasker.org`, 2013-2015.

[13] Robert Hubley, Robert D Finn, Jody Clements, Sean R Eddy, Thomas A Jones, Weidong Bao, Arian FA Smit, and Travis J Wheeler. The dfam database of repetitive dna families. *Nucleic acids research*, 44(D1):D81–D89, 2016.

[14] Stephen F Altschul, Warren Gish, Webb Miller, Eugene W Myers, and David J Lipman. Basic local alignment search tool. *Journal of molecular biology*, 215(3):403–410, 1990.

[15] Jessica Storer, Robert Hubley, Jeb Rosen, Travis J Wheeler, and Arian F Smit. The dfam community resource of transposable element families, sequence models, and genome annotations. *Mobile DNA*, 12(1):1–14, 2021.

[16] Manfred G Grabherr, Brian J Haas, Moran Yassour, Joshua Z Levin, Dawn A Thompson, Ido Amit, Xian Adiconis, Lin Fan, Raktima Raychowdhury, Qiandong Zeng, et al. Trinity: reconstructing a full-length transcriptome without a genome from rna-seq data. *Nature biotechnology*, 29(7):644, 2011.

[17] Thomas D Wu and Colin K Watanabe. Gmap: a genomic mapping and alignment program for mrna and est sequences. *Bioinformatics*, 21(9):1859–1875, 2005.

[18] Brian J Haas, Arthur L Delcher, Stephen M Mount, Jennifer R Wortman, Roger K Smith Jr, Linda I Hannick, Rama Maiti, Catherine M Ronning, Douglas B Rusch, Christopher D Town, et al. Improv-

ing the arabidopsis genome annotation using maximal transcript alignment assemblies. *Nucleic acids research*, 31(19):5654–5666, 2003.

[19] Brian J Haas, Steven L Salzberg, Wei Zhu, Mihaela Pertea, Jonathan E Allen, Joshua Orvis, Owen White, C Robin Buell, and Jennifer R Wortman. Automated eukaryotic gene structure annotation using evidencemodeler and the program to assemble spliced alignments. *Genome biology*, 9:1–22, 2008.

[20] Katharina J Hoff, Alexandre Lomsadze, Mark Borodovsky, and Mario Stanke. Whole-genome annotation with braker. *Gene prediction: methods and protocols*, pages 65–95, 2019.

[21] W James Kent. Blat—the blast-like alignment tool. *Genome research*, 12(4):656–664, 2002.

[22] Felix Krueger and Simon R Andrews. Bismark: a flexible aligner and methylation caller for bisulfite-seq applications. *bioinformatics*, 27(11):1571–1572, 2011.

[23] Aimée M Deaton and Adrian Bird. Cpg islands and the regulation of transcription. *Genes & development*, 25(10):1010–1022, 2011.

[24] Fábio Madeira, Matt Pearce, Adrian RN Tivey, Prasad Basutkar, Joon Lee, Ossama Edbali, Nandana Madhusoodanan, Anton Kolesnikov, and Rodrigo Lopez. Search and sequence analysis tools services from embl-ebi in 2022. *Nucleic acids research*, 50(W1):W276–W279, 2022.

[25] Francisco Antequera and Adrian Bird. Number of cpg islands and genes in human and mouse. *Proceedings of the National Academy of Sciences*, 90(24):11995–11999, 1993.

[26] Drew R Schield, Giulia IM Pasquesi, Blair W Perry, Richard H Adams, Zachary L Nikolakis, Aundrea K Westfall, Richard W Orton, Jesse M Meik, Stephen P Mackessy, and Todd A Castoe. Snake recombination landscapes are concentrated in functional regions despite prdm9. *Molecular Biology and Evolution*, 37(5):1272–1294, 2020.

[27] Olga Dudchenko, Sanjit S Batra, Arina D Omer, Sarah K Nyquist, Marie Hoeger, Neva C Durand, Muhammad S Shamim, Ido Machol, Eric S Lander, Aviva Presser Aiden, et al. De novo assembly of the aedes aegypti genome using hi-c yields chromosome-length scaffolds. *Science*, 356(6333):92–95, 2017.

[28] Michael A Quail, Harold Swerdlow, and Daniel J Turner. Improved protocols for the illumina genome analyzer sequencing system. *Current protocols in human genetics*, 62(1):18–2, 2009.

[29] Anthony M Bolger, Marc Lohse, and Bjoern Usadel. Trimmomatic: a flexible trimmer for illumina sequence data. *Bioinformatics*, 30(15):2114–2120, 2014.

[30] Heng Li. Aligning sequence reads, clone sequences and assembly contigs with bwa-mem. *arXiv preprint arXiv:1303.3997*, 2013.

[31] Heng Li, Bob Handsaker, Alec Wysoker, Tim Fennell, Jue Ruan, Nils Homer, Gabor Marth, Goncalo Abecasis, Richard Durbin, 1000 Genome Project Data Processing Subgroup, et al. The sequence alignment/map (sam) format and samtools. *Bioinformatics*, 25(16):2078–2079, 2009.

[32] Picard toolkit. `https://broadinstitute.github.io/picard/`, 2019.

[33] Ryan Poplin, Valentin Ruano-Rubio, Mark A DePristo, Tim J Fennell, Mauricio O Carneiro, Geraldine A Van der Auwera, David E Kling, Laura D Gauthier, Ami Levy-Moonshine, David Roazen, et al. Scaling accurate genetic variant discovery to tens of thousands of samples. *BioRxiv*, page 201178, 2017.

[34] Petr Danecek, Adam Auton, Goncalo Abecasis, Cornelis A Albers, Eric Banks, Mark A DePristo, Robert E Handsaker, Gerton Lunter, Gabor T Marth, Stephen T Sherry, et al. The variant call format and vcftools. *Bioinformatics*, 27(15):2156–2158, 2011.

[35] Ani Manichaikul, Josyf C Mychaleckyj, Stephen S Rich, Kathy Daly, Michèle Sale, and Wei-Min Chen. Robust relationship inference in genome-wide association studies. *Bioinformatics*, 26(22):2867–2873, 2010.

[36] Geraldine A Van der Auwera and Brian D O'Connor. *Genomics in the cloud: using Docker, GATK, and WDL in Terra*. O'Reilly Media, 2020.

[37] Petr Danecek, James K Bonfield, Jennifer Liddle, John Marshall, Valeriu Ohan, Martin O Pollard, Andrew Whitwham, Thomas Keane, Shane A McCarthy, Robert M Davies, et al. Twelve years of samtools and bcftools. *Gigascience*, 10(2):giab008, 2021.

[38] Heng Li. Tabix: fast retrieval of sequence features from generic tab-delimited files. *Bioinformatics*, 27(5):718–719, 2011.

[39] Olivier Delaneau, Jean-Francois Zagury, and Jonathan Marchini. Improved whole-chromosome phasing for disease and population genetic studies. *Nature methods*, 10(1):5–6, 2013.

[40] Christopher C Chang, Carson C Chow, Laurent CAM Tellier, Shashaank Vattikuti, Shaun M Purcell, and James J Lee. Second-generation plink: rising to the challenge of larger and richer datasets. *Gigascience*, 4(1):s13742–015, 2015.

[41] A. H. Chan, P.A. Jenkins, and Y. S. Song. Genome-wide fine-scale recombination rate variation in drosophila melanogaster. *PLoS genetics*, 8(12):e1003090, 2012.

[42] Zachary Baker, Molly Schumer, Yuki Haba, Lisa Bashkirova, Chris Holland, Gil G Rosenthal, and Molly Przeworski. Repeated losses of prdm9-directed recombination despite the conservation of prdm9 across vertebrates. *Elife*, 6:e24133, 2017.

[43] Anton V Persikov and Mona Singh. De novo prediction of dna-binding specificities for cys2his2 zinc finger proteins. *Nucleic acids research*, 42(1):97–108, 2014.

[44] Charles E Grant, Timothy L Bailey, and William Stafford Noble. Fimo: scanning for occurrences of a given motif. *Bioinformatics*, 27(7):1017–1018, 2011.

[45] Jerrod J Schwartz, David J Roach, James H Thomas, and Jay Shendure. Primate evolution of the recombination regulator prdm9. *Nature communications*, 5(1):4370, 2014.

[46] Jérôme Buard, Eric Rivals, Denis Dunoyer de Segonzac, Charlotte Garres, Pierre Caminade, Bernard de Massy, and Pierre Boursot. Diversity of prdm9 zinc finger array in wild mice unravels new facets of the evolutionary turnover of this coding minisatellite. *PloS one*, 9(1):e85021, 2014.

[47] Kazutaka Katoh, John Rozewicki, and Kazunori D Yamada. Mafft online service: multiple sequence alignment, interactive sequence choice and visualization. *Briefings in bioinformatics*, 20(4):1160–1166, 2019.

[48] Prdm9 is a major determinant of meiotic recombination hotspots in humans and mice. *Science*, 327(5967):836–840, 2010.

[49] Simon Myers, Rory Bowden, Afidalina Tumian, Ronald E Bontrop, Colin Freeman, Tammie S MacFie, Gil McVean, and Peter Donnelly. Drive against hotspot motifs in primates implicates the prdm9 gene in meiotic recombination. *Science*, 327(5967):876–879, 2010.

[50] Timothy L Bailey, James Johnson, Charles E Grant, and William S Noble. The meme suite. *Nucleic acids research*, 43(W1):W39–W49, 2015.

[51] Aaron R Quinlan and Ira M Hall. Bedtools: a flexible suite of utilities for comparing genomic features. *Bioinformatics*, 26(6):841–842, 2010.

[52] Bruce S Weir and C Clark Cockerham. Estimating f-statistics for the analysis of population structure. *evolution*, pages 1358–1370, 1984.

[53] Jonathan Terhorst, John A Kamm, and Yun S Song. Robust and scalable inference of population history from hundreds of unphased whole genomes. *Nature genetics*, 49(2):303–309, 2017.

[54] Lucie A Bergeron, Søren Besenbacher, Jiao Zheng, Panyi Li, Mads Frost Bertelsen, Benoit Quintard, Joseph I Hoffman, Zhipeng Li, Judy St. Leger, Changwei Shao, et al. Evolution of the germline mutation rate across vertebrates. *Nature*, 615(7951):285–291, 2023.

[55] Jeffrey P Spence and Yun S Song. Inference and analysis of population-specific fine-scale recombination maps across 26 diverse human populations. *Science Advances*, 5(10):eaaw9206, 2019.

[56] GA Watterson. On the number of segregating sites in genetical models without recombination. *Theoretical population biology*, 7(2):256–276, 1975.

[57] Joel Armstrong, Glenn Hickey, Mark Diekhans, Ian T Fiddes, Adam M Novak, Alden Deran, Qi Fang, Duo Xie, Shaohong Feng, Josefin Stiller, et al. Progressive cactus is a multiple-genome aligner for the thousand-genome era. *Nature*, 587(7833):246–251, 2020.

[58] Marc de Manuel, Felix L Wu, and Molly Przeworski. A paternal bias in germline mutation is widespread across amniotes and can arise independently of cell divisions. *bioRxiv*, 2022.

[59] Michael I Jensen-Seaman, Terrence S Furey, Bret A Payseur, Yontao Lu, Krishna M Roskin, Chin-Fu Chen, Michael A Thomas, David Haussler, and Howard J Jacob. Comparative recombination rates in the rat, mouse, and human genomes. *Genome research*, 14(4):528–538, 2004.

[60] Timothy L Bailey. Streme: accurate and versatile sequence motif discovery. *Bioinformatics*, 37(18):2834–2840, 2021.

[61] Tremblay BJ (2023). *universalmotif: Import, Modify, and Export Motifs with R*, 2023. R package version 1.18.1.

[62] Glenn Hickey, Benedict Paten, Dent Earl, Daniel Zerbino, and David Haussler. Hal: a hierarchical format for storing and analyzing multiple genome alignments. *Bioinformatics*, 29(10):1341–1342, 2013.

[63] Julien Meunier and Laurent Duret. Recombination drives the evolution of gc-content in the human genome. *Molecular biology and evolution*, 21(6):984–990, 2004.

[64] Graham Coop, Xiaoquan Wen, Carole Ober, Jonathan K Pritchard, and Molly Przeworski. High-resolution mapping of crossovers reveals extensive variation in fine-scale recombination patterns among humans. *science*, 319(5868):1395–1398, 2008.

[65] Drew R Schield, Daren C Card, Nicole R Hales, Blair W Perry, Giulia M Pasquesi, Heath Blackmon, Richard H Adams, Andrew B Corbin, Cara F Smith, Balan Ramesh, et al. The origins and evolution of chromosomes, dosage compensation, and mechanisms underlying venom regulation in snakes. *Genome research*, 29(4):590–601, 2019.

[66] Maria Izabel A Cavassim, Zachary Baker, Carla Hoge, Mikkel H Schierup, Molly Schumer, and Molly Przeworski. Prdm9 losses in vertebrates are coupled to those of paralogs zcwpw1 and zcwpw2. *Proceedings of the National Academy of Sciences*, 119(9):e2114401119, 2022.

[67] Fiona Cunningham, James E Allen, Jamie Allen, Jorge Alvarez-Jarreta, M Ridwan Amode, Irina M Armean, Olanrewaju Austine-Orimoloye, Andrey G Azov, If Barnes, Ruth Bennett, et al. Ensembl 2022. *Nucleic acids research*, 50(D1):D988–D995, 2022.

[68] Nuala A O'Leary, Mathew W Wright, J Rodney Brister, Stacy Ciufo, Diana Haddad, Rich McVeigh, Bhanu Rajput, Barbara Robbertse, Brian Smith-White, Danso Ako-Adjei, et al. Reference sequence (refseq) database at ncbi: current status, taxonomic expansion, and functional annotation. *Nucleic acids research*, 44(D1):D733–D745, 2016.

[69] Uniprot: the universal protein knowledgebase in 2023. *Nucleic Acids Research*, 51(D1):D523–D531, 2023.

[70] Wei Shen, Shuai Le, Yan Li, and Fuquan Hu. Seqkit: a cross-platform and ultrafast toolkit for fasta/q file manipulation. *PloS one*, 11(10):e0163962, 2016.

[71] Aron Marchler-Bauer and Stephen H Bryant. Cd-search: protein domain annotations on the fly. *Nucleic acids research*, 32(suppl_2):W327–W331, 2004.

[72] Aron Marchler-Bauer, Shennan Lu, John B Anderson, Farideh Chitsaz, Myra K Derbyshire, Carol DeWeese-Scott, Jessica H Fong, Lewis Y Geer, Renata C Geer, Noreen R Gonzales, et al. Cdd: a conserved domain database for the functional annotation of proteins. *Nucleic acids research*, 39(suppl_1):D225–D229, 2010.

[73] Jaina Mistry, Sara Chuguransky, Lowri Williams, Matloob Qureshi, Gustavo A Salazar, Erik LL Sonnhammer, Silvio CE Tosatto, Lisanna Paladin, Shriya Raj, Lorna J Richardson, et al. Pfam: The protein families database in 2021. *Nucleic acids research*, 49(D1):D412–D419, 2021.

[74] Michael Y Galperin, Yuri I Wolf, Kira S Makarova, Roberto Vera Alvarez, David Landsman, and Eugene V Koonin. Cog database update: focus on microbial diversity, model organisms, and widespread pathogens. *Nucleic acids research*, 49(D1):D274–D281, 2021.

[75] Wenjun Li, Kathleen R O'Neill, Daniel H Haft, Michael DiCuccio, Vyacheslav Chetvernin, Azat Badretdin, George Coulouris, Farideh Chitsaz, Myra K Derbyshire, A Scott Durkin, et al. Refseq: expanding the prokaryotic genome annotation pipeline reach with protein family model curation. *Nucleic acids research*, 49(D1):D1020–D1028, 2021.

[76] Robert C Edgar. Muscle: multiple sequence alignment with high accuracy and high throughput. *Nucleic acids research*, 32(5):1792–1797, 2004.

[77] Alexandros Stamatakis. Raxml version 8: a tool for phylogenetic analysis and post-analysis of large phylogenies. *Bioinformatics*, 30(9):1312–1313, 2014.

[78] Ziheng Yang et al. Paml: a program package for phylogenetic analysis by maximum likelihood. *Computer*

*applications in the biosciences*, 13(5):555–556, 1997.

[79] Itamar Sela, Haim Ashkenazy, Kazutaka Katoh, and Tal Pupko. Guidance2: accurate detection of unreliable alignment regions accounting for the uncertainty of multiple parameters. *Nucleic acids research*, 43(W1):W7–W14, 2015.

[80] Mikita Suyama, David Torrents, and Peer Bork. Pal2nal: robust conversion of protein sequence alignments into the corresponding codon alignments. *Nucleic acids research*, 34(suppl_2):W609–W612, 2006.

[81] Salvador Capella-Gutiérrez, José M Silla-Martínez, and Toni Gabaldón. trimal: a tool for automated alignment trimming in large-scale phylogenetic analyses. *Bioinformatics*, 25(15):1972–1973, 2009.

[82] Bernard V North, David Curtis, and Pak C Sham. A note on the calculation of empirical p values from monte carlo procedures. *The American Journal of Human Genetics*, 71(2):439–441, 2002.
